# Discovery and molecular basis of chloride as an allosteric activator and catalytic inhibitor for Class-D β-lactamases

**DOI:** 10.1101/2022.12.20.520813

**Authors:** Qi Zhou, Pablo Catalán, Helen Bell, Patrick Baumann, Rhodri Evans, Jianhua Yang, Zhen Zhang, Davide Zappalà, Ye Zhang, George Michael Blackburn, Yuan He, Yi Jin

## Abstract

Oxacillinase (OXA)-48-like carbapenemases are epidemic class D β-lactamases in *Enterobacterales*, resulting in high mortality. Though the chemical mechanism has been clearly established, for decades, the link between the biphasic kinetic behaviour of these enzymes, which significantly impacts antibiotic efficacy, and the state of carbamylated lysine has been elusive. Here, substituting *N-*carbamylated lysine73 with a chemically-stable *N*-acetyl lysine allows us to prove the origin of catalytic inhibition is not decarbamylation and enables us to capture an unprecedented inactive acyl-intermediate wedged in place by a chloride ion against the conserved residue arginine250. We here identify chloride as a “Janus effector” acting by allosteric activation of the burst phase and inhibition of the steady-state for a series of β-lactam substrates in kinetic assays. Chloride ions are necessarily present in both laboratory and clinical OXA activity assays and their inseparable role is now identified. Our finding suggests a new direction for the discovery of next-generation antibiotics specific for β-lactamases of Class D.

## Introduction

Antibiotic resistance is currently a major, global threat to health^1,2^. Carbapenems are last-line agents for treatment of serious infections^3,4^ as clinically-important β-lactam antibiotics with a broad spectrum of activity and high potency. However, with widespread dissemination of plasmid-encoded carbapenemases, resistance to carbapenems is posing a significant challenge to current clinical anti-infection therapy^5,6^. Three classes of carbapenemases have now been identified: Ambler Class A (e.g. KPC), Class B (e.g. NDM), and Class D (oxacillinases, OXA type)^7,8^. Class D β-lactamases have a post-translationally modified *N*-carbamylated catalytic lysine^†^ residue (KCX) formed reversibly from atmospheric, soluble CO2 not seen in Class A and C serine β-lactamases^9,10^. The carbamylated lysine-73 (*pK_a_* 5.6) provides orientation and general base catalysis for serine nucleophilic attack in the formation of the acyl intermediate in Step 1 and the nucleophilic water molecule in the often rate-limiting Step 2^10^. Compared to other β-lactamases, there has been slow progress on mechanism-based inhibitor design for Class D OXAs, not least because the mechanism of this large Class of enzymes is not well understood. OXAs-2, −10, −14, −16, −27, −50, −48 all show partial inactivation after repeated cycles of catalysis, displaying peculiar “biphasic kinetics” with β-lactams^9,11–16^. This partial inactivation has the capability to change the pharmacodynamics of antibiotics for effective treatment. Consequently, there has been much research seeking to identify the underlying molecular origin of this obscure behaviour.

For many years, a “partition ratio” *k*_active_/*k*_inactive_ has been used to describe the turnover number before the OXA enzyme is inactivated. Several studies attributed it to the ratio between the uncarbamylated inactive form and carbamylated active form of the OXA^9,10,17^. Because of this, addition of NaHCO_3_ has been used routinely to reverse inhibition and prevent decarbamylation of KCX during turnover^9,17^. Several studies that supported inactivation of OXA-48_WT_ by decarbamylation of K73 employed avibactam, a bicyclic diazabicyclooctane covalent inhibitor^18,19^. The use of ^13^C and ^19^F NMR to directly or indirectly monitor the ^13^C-labelled KCX state of the acylated OXA-avibactam complex suggested decarbamylation was caused by super-high concentrations (500 mM) of halide ions outcompeting the carbamylate group on Lys^18,19^. However, more recent studies have proposed that avibactam-bound β-lactamases intrinsically favour K73 decarbamylation^19,20^ (**Table S1**). Significantly, OXA-24 and also OXAs-25, −26, −27, −40, −48, −50, −58, and −163 *etc*. have been found to be partially inactivated by chloride ion *in vitro*^14,15,21–24^. The IC_50_ values of chloride ion for several OXAs were reported in the physiologically relevant mM range^23^. Chloride is an important electrolyte in life with a concentration between 15-120 mM^25–27^, though this can vary substantially in diseases^26–29^. NaCl is also a commonly used component in buffers for kinetic assays, in growth media for antibiotic susceptibility tests, and in diluents for antimicrobial treatment. Hence, elucidating how chloride ions modulate OXA activities and its relationship with biphasic kinetics is of particular importance for a comprehensive understanding of the catalytic mechanism and disease treatment.

There are two main challenges for establishing the molecular basis of biphasic OXA kinetics. Firstly, although the activity of OXAs can be affected by the KCX state of K73, there is no consensus on whether decarbamylation occurs during turnover, making it an urgent subject for debate^30^. Decarbamylation is susceptible to many factors, including pH^20,31,32^, meaning that in techniques, *e.g*. crystallography and NMR, that call for millimolar substrate concentration, carbamylation can dramatically diminish because of acidification from protons released by β-lactam hydrolysis^33,34^. Secondly, state-of-the-art studies of OXAs have focused mainly on measuring initial reaction rates in biphasic kinetics using UV-vis spectroscopy, thus overlooking more profound kinetic processes later in the reaction.

To capture subtle rate changes throughout the biphasic kinetics of OXA-48 reaction, which has a characteristic fast hydrolytic first phase followed by a quick transition to a slow but steady second phase^34,35^, we have employed a previously developed and highly sensitive Isothermal Titration Calorimetry (ITC) assay that detects Carbapenemase-Producing *Enterobacteriaceae* (CPE) efficiently. Moreover, we have shown ITC is a very accurate and reproducible method for monitoring β-lactam cleavage reactions^16^. By this means, we observed the first phase rate-acceleration increased when the NaHCO_3_ concentration was increased from 10 mM to 50 mM, well beyond the dissociation concentration of CO_2_ (0.23 μM)^9,16^. That result is inconsistent with decarbamylation during catalysis. Rather, it indicates that the proposed mechanism of KCX decarbamylation during turnover needs reexamination and it establishes that the relationship between biphasic kinetics, KCX state, and chloride during catalysis remains poorly understood.

Herein, we focus on OXA-48, one of the most noxious OXAs, that is often encoded by high-death-causing carbapenem-resistant *Enterobacterales*^8,36–38^. To test directly whether KCX undergoes decarbamylation during turnover, we have incorporated the non-natural amino acid *N-*acetyl-lysine (AcK) into position-73 of OXA-48, as a mimic of an un-decarbamylatable OXA-48. We utilized our continuous ITC assay to trace the complete reaction course of OXA-48 under various conditions (*e.g*., enzyme and substrate concentrations, halide concentrations, *etc.*) and identified halogen ions, namely chloride, bromide and iodide, as the ultimate determinants of biphasic kinetics in OXA-48 reactions with a series of β-lactam substrates. X-ray crystallography complemented by intrinsic fluorescence enabled us to discover an “inactive mode” of the reaction intermediate in the OXA-48AcK73-imipenem complex structure, in which the halide ion is “wedged” between the acyl-imipenem intermediate and active site residue R250. Mutagenesis and molecular docking further established R250 as an essential residue for enzyme inhibition. Our mutagenesis study has also identified that binding of chloride to three other surface residues can activate the enzyme allosterically. These observations together prove that chloride ion is a “Janus effector” of OXA-48 responsible for its biphasic kinetics. Based on all our above evidence and mathematical simulation, we propose a new catalytic inhibition reaction pathway involving chloride. Since chloride ions are universally present, our work provides a molecular definition for guiding consistent kinetics and minimum inhibitory concentration (MIC) results in both laboratory and clinical settings. It also draws attention to the effect of chloride on pharmacokinetics and efficacy of antibiotic drugs during *in vivo* treatment. Our findings are informative both for future design of new class of OXAs inhibitors and for preserving the effectiveness of current antibiotics and equally for CPE detection.

## Results

### Chloride affects bacterial growth and CPE detection

We monitored the growth curves of three prevalent CPE *Klebsiella pneumoniae* strains producing OXA-48 (Class D), KPC (Class A) and NDM (Class B), respectively, by measuring turbidity at 600 nm (**Fig. 1**). The growth rate of all three bacteria showed a slight inhibition in terrific broth (TB) supplied with chloride ions, a stronger inhibition effect in TB with the antibiotics, and co-addition of chloride ions and antibiotics resulted in the strongest inhibition effect. Such synergistic effect of chloride and imipenem is most significant for *K. pneumoniae* encoding OXA-48 showing absolutely no growth within 12 h (**Fig. 1a**). This implies that the detrimental effect of chloride on the growth of OXA-48-encoding *K. pneumoniae* could be related to the effect of chloride on OXA-48 carbapenemase activity. We then carried out living-cell ITC studies^16^ to compare calorimetric curves of imipenem hydrolysis by OXA-48-encoding *K. pneumoniae* under chloride-present and chloride-free conditions. The calorimetric curves were recorded when imipenem was titrated directly into *K. pneumoniae* cell suspensions. The inclusion of either 100 or 400 mM chloride in a phosphate buffer led to a biphasic curve of OXA-48-encoding *K. pneumoniae* (**Fig. 1d**), but showed no influence on the curves for both KPC-encoding and NDM-encoding *K. pneumoniae* (**Fig. 1e, f**). This demonstrates that biphasic kinetics is uniquely caused by chloride in Class D OXA-48 and is relevant *in vivo* for imipenem antibiotic use.

**Figure 1.**
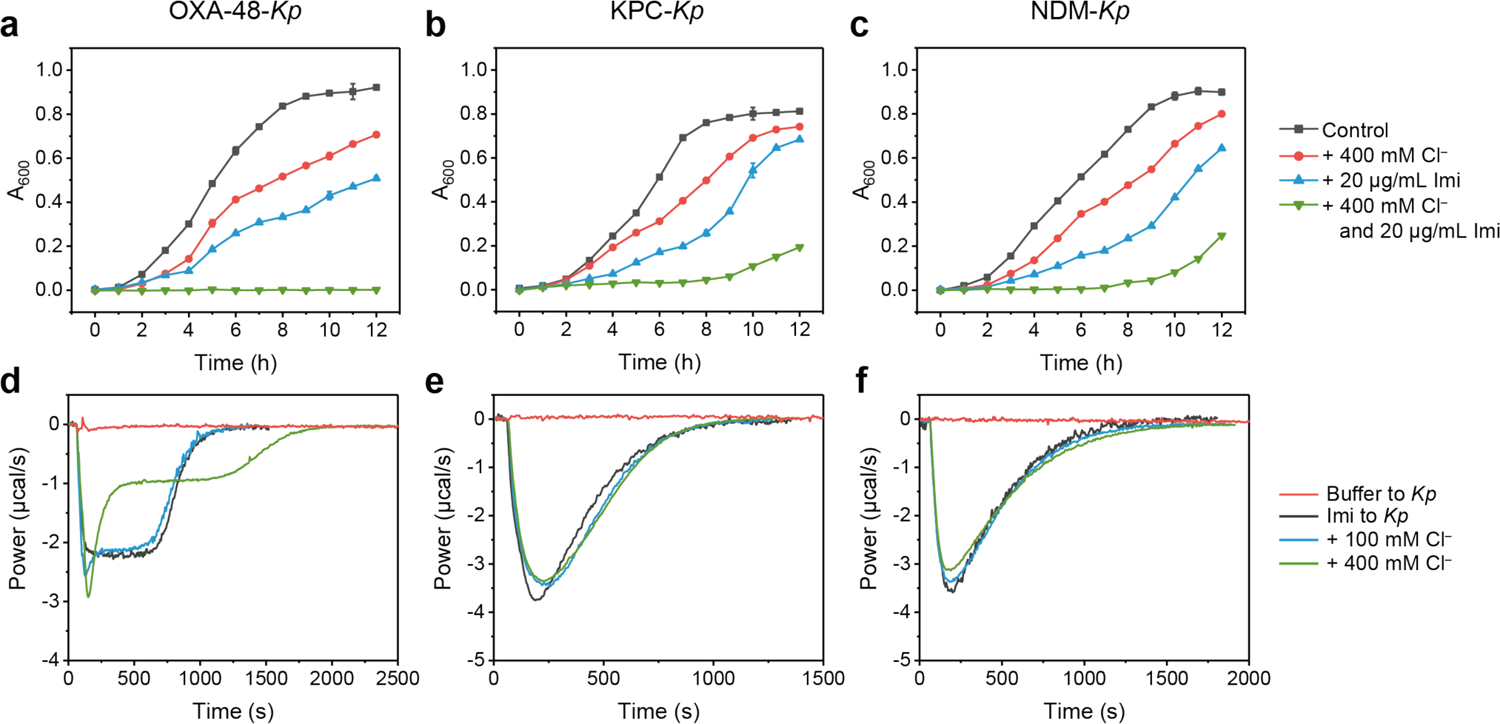
Growth curves of (**a**) Class D OXA-48-*Kp* bacteria, (**b**) Class A *KPC-Kp* bacteria, and (**c**) Class B NDM-*Kp* bacteria under variable conditions (400 mM NaCl and 20 μg/mL imipenem (Imi) in TB buffer). Thermogram curves of imipenem hydrolysis in the absence and presence of chloride by cell suspensions of (**d**) OXA-48-*Kp*, (**e**) KPC-*Kp* and (**f**) NDM-*Kp*, in 50 mM NaPi buffer, pH 7.5, and 0.1 mM ZnSO4.

### Chloride is the cause of biphasic kinetics

Given the role of chloride is at the heart of this study, we excluded every possible introduction of chloride in all steps of OXA-48 protein preparation. To this end, TB medium without supplement of NaCl was used for recombinant gene expression and 50 mM Na_2_HPO_3_-NaH_2_PO_3_ pH 7.5 was used as the buffer throughout protein purification and activity studies in this study. Studies of the state of carbamylation on lysine have always been challenging, given its labile nature^39^. To confirm that biphasic kinetics is not caused by chloride-dependent decarbamylation, we synthesized *N*-acetylated lysine (AcK, **Fig. 2a**), a structurally similar but non-hydrolyzable mimic of the carbamylated lysine, and incorporated it into OXA-48_AcK73_ variant using genetic code expansion technology. High-resolution mass spectrometric (HR-MS) analysis showed 100% AcK incorporation (**Fig. S1**). Michaelis-Menten kinetics for OXA-48_AcK73_ variant and OXA-48_WT_ by UV-vis spectroscopy (**Fig. S2**) show a similar *K*_M_ for OXA-48_AcK73_ (13.89 μM) and OXA-48_WT_ (13.79 μM), meaning that AcK substitution does not interfere with substrate binding. However, the 40-fold decrease of *k*_cat_ for OXA-48_AcK73_ (0.13 s^-1^) compared to OXA-48_WT_ (5.13 s^-1^) shows the important contribution of KCX to catalysis.

**Figure 2.**
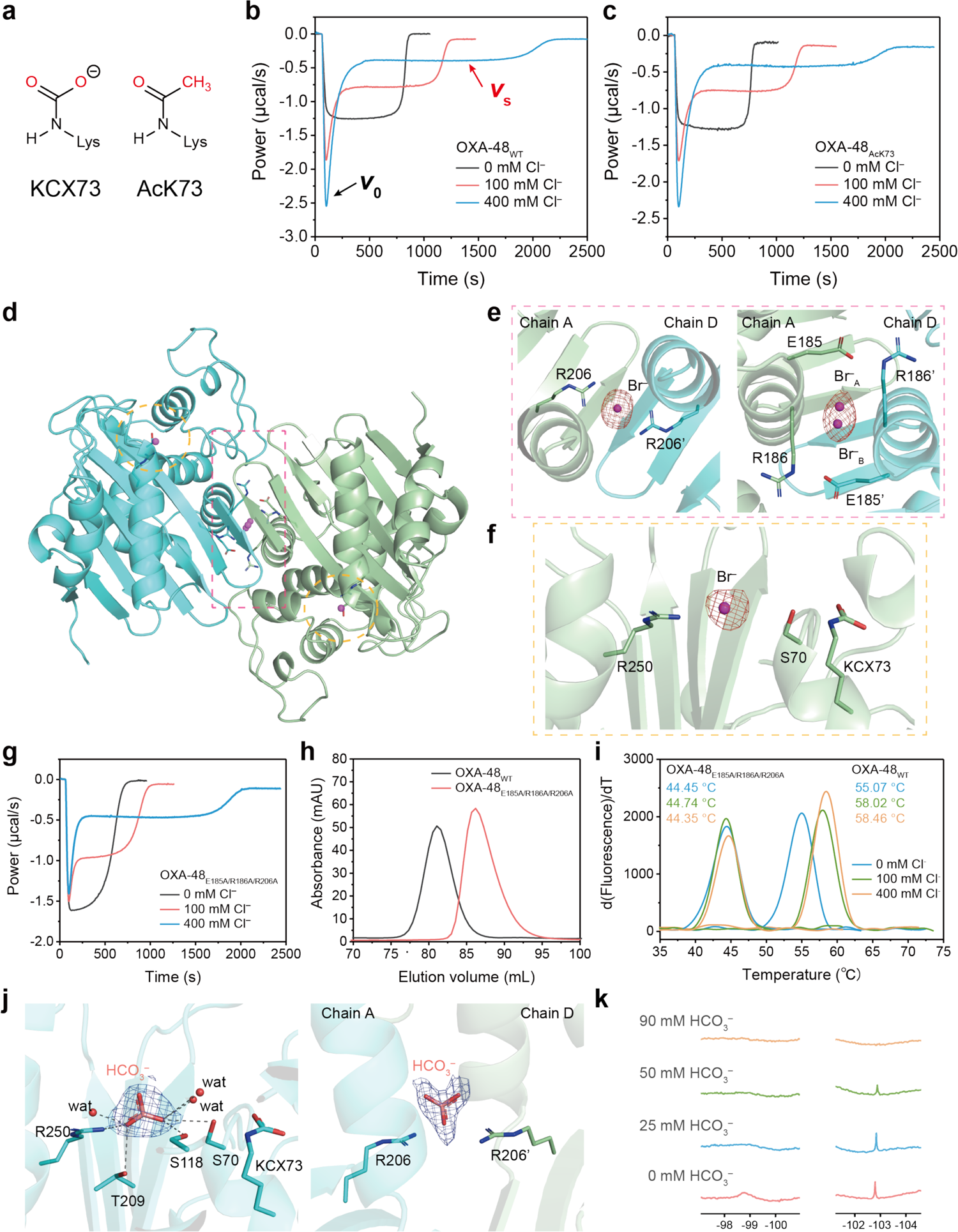
(**a**) Comparison of the structures of carbamylation (KCX73) and the acetylation (AcK73) modifications of K73. Effect of [Cl^-^] on the catalytic activity of (**b**) 100 nM OXA-48_WT_ and (**c**) 8 μM OXA-48_AcK73_ variant with 200 μM imipenem in a calorimetry assay, measured in 50 mM NaPi buffer, pH 7.5 and 1 mM NaHCO_3_. (**d**) A cartoon to show the dimeric assembly from crystal structure of OXA-48_WT_-Br^-^ complex; bromide ions bound on the dimer interface and in the active sites are shown in purple spheres. (**e**) Binding sites for halide ions at OXA-48_WT_ enzyme dimer interface (Anomalous map of Br is shown as red mesh, contoured at 3σ). (**f**) Binding site for halide ion at the OXA-48_WT_ active site (anomalous density map of Br is shown as red mesh, contoured at 3σ). (**g**) Kinetic curves of 200 μM imipenem hydrolysis by 100 nM interface variant OXA-48_E185A/R186A/R206A_ with various [Cl^-^] in 50 mM NaPi buffer, pH 7.5 and 1 mM NaHCO_3_. (**h**) Size exclusion chromatography of OXA-48_WT_ and the interface variant OXA-48_E185A/R186A/R206A_, run on a HiLoad 16/600 Superdex 200 pg column (GE Healthcare) in buffer 50 mM NaPi, pH 7.5. (**i**) Protein thermal shift melting curves for OXA-48_E185A/R186A/R206A_ and OXA-48_WT_ with different concentrations of chloride ion. (**j**) Crystal structure of OXA-48_WT_-HCO_3_^-^ complex shows HCO_3_^-^ ion bound to R250 (left) and at the dimer interface R206-R206’ (right). The initial unbiased 2*F*_o_-*F*_c_ electron density map of HCO_3_^-^ is shown as blue mesh, contoured at 1*σ* (0.3625 e^-^/Å ^3^). (**k**) ^19^F NMR spectra of a sample of 500 μM OXA-48_WT_ in the buffer of 500 mM NH4F, 100 mM NaPi, pH 7.5, 10% D_2_O titrated with 0-90 mM NaHCO_3_ to show HCO_3_^-^ ion outcompetes fluoride at lower concentrations for at least two binding sites.

ITC directly monitors changes in heat release rate associated with enzymatic activity as a function of time. This means that subtle changes in enzyme activity can be more easily and accurately observed by ITC than by UV-vis (**Fig. S3**). The heat profile curves of both OXA-48_WT_ and OXA-48_AcK73_ in 50 mM phosphate buffer, pH 7.5 at 25 °C, consistently showed a typical monophasic calorimetric curve under Cl^-^-free conditions but a distinct biphasic calorimetric curve with chloride present (**Fig. 2b, c**). These observations demonstrate that chloride is the cause of biphasic kinetics, and the variable activity is not due to KCX decarbamylation.

A closer investigation of the biphasic curves showed that increasing [Cl^-^] accelerates the first burst phase while it inhibits the second steady phase (**Fig. S4**). We next determined EC_50_ and IC_50_ for chloride and other halogen ions deduced from the initial burst phase rate *v*_0_ and the steady inhibitory phase rate *v*_s_^11^. We found that chloride and bromide ions show a similar enhancement in activity for the burst phase (**Fig. S5, 6, Table S2**) while fluoride only raises the steady-state rate (EC_50_ >222 mM). However, steady-state IC_50_ values vary according to the size of the halide ion, with larger ions displaying stronger inhibition and F^-^ not showing any inhibition. These data suggest that Cl^-^, Br^-^ and I modulate enzyme activity by a different mechanism from F^-^, not simply by the change of ionic strength.

### Chloride is a positive allosteric effector

To ensure all crystal structures are physiologically relevant with full occupancy of KCX, all OXA-48 proteins in this paper were crystallized at neutral pH under Cl^-^-free conditions. The OXA-48_WT_ apo structure was solved as a dimer (1.92 Å, **Table S3**). We then soaked NaBr into OXA-48_WT_ apo-crystals for anomalous data collection and the best structure of OXA-48_WT_ with bromide was solved at 1.81 Å resolution (**Fig. 2d, Table S3**). The anomalous signal (**Fig. 2e**, red mesh) clearly identified two bromides bound at the previously identified OXA-48 dimer interface^40^, namely between side chains of R206-R206’ and the side chains of E185-R186-E185’-R186’ (**Fig. 2d, e**). Moreover, a bromide was found next to R250 in the active site (**Fig. 2f**).

To relate these halide-binding residues to the observed kinetics, we also prepared interface variant OXA-48_E185A/R186A/R206A_ apo crystals and soaked them with 500 mM NaBr. The 1.9 Å resolution structure (**Table S3**) showed no electron density of bromide at the dimer interface (**Fig. S7**). The OXA-48_E185A/R186A/R206A_ variant showed similar monophasic activity as OXA-48_WT_ in Cl^-^-free buffer (**Fig. 2g**), yet a much smaller burst phase *vo* in biphasic kinetics was still observed with chloride (**Fig. 2g and S8**), suggesting that chloride binding to these interfacial residues accounts for the positive allosteric effect. Size exclusion chromatography revealed that OXA-48_WT_ is a strongly stabilized dimer, while interface variant OXA-48_E185A/R186A/R206A_ exists as a monomer or as a weakly-associated dimer in solution (**Fig. 2h**). Protein thermal shift assays also showed that *T*_m_ for OXA-48_WT_ is 55 °C in the pH 7.5 Cl^-^-free buffer, *i.e*., 10 °C higher than that of OXA-48_E185A/R186A/R206A_ (**Fig. 2i**). The addition of chloride further increases the *T*_m_ of OXA-48_WT_ by ~3.5 °C but does not do so for OXA-48_E185A/R186A/R206A_ (**Table S4**). Our data suggest that chloride binding to E185, R186 and R206 at the enzyme dimer interface endows the enzyme with higher stability and also leads to a greater allosteric effect in the burst phase.

We then obtained a structure of OXA-48_WT_ apo crystal soaked with NaHCO_3_, in which HCO_3_^-^ also interacts with R206-R206’ at the dimer interface as do halides (**Fig. 2j**). To test whether HCO_3_^-^ is an allosteric effector, we tested the activity of both OXA-48_WT_ and OXA-48_AcK73_ and showed the imipenem hydrolytic rate for both enzymes doubled when the [HCO_3_^-^] was increased from 0 mM to 400 mM (**Fig. S9**). This shows that the “apparent rescuing” effect of HCO_3_^-^ report in the literature is not due to blocking decarbamylation of KCX. Instead, it has a similar positive allosteric effect to F^-^ (**Fig. S5**). We also did a competitive binding assay by solution ^19^F NMR to show an increase of [HCO_3_^-^] from 25 mM to 100 mM totally outcompeted the surface-bound F^-^ at 500 mM for the same binding sites (**Fig. 2k**).

### Crystal structure reveals unique inactive acyl-intermediate conformation

To provide further molecular insight into the mechanism of chloride-induced biphasic kinetics, we added 500 mM NaBr and 32 mM imipenem to the apo OXA-48_WT_ crystal drop. That resulted in a clear Br^-^ anomalous signal overlapping with the density of imipenem to establish Br^-^ and imipenem compete for binding to R250 (**Fig. 3a**, 1.53 Å resolution structure, undeposited). Based on this observation, we then carried out further structural investigation of OXA-48_AcK73_ crystals for its significantly slow catalysis that might facilitate trapping of key reaction intermediates. We soaked imipenem and bromide into OXA-48_AcK73_ crystals at neutral pH and solved the best structure to 2.15 Å resolution (**Table S3**), showing four copies of dimers in the asymmetric unit (ASU). All chains showed clear electron density for the unbiased *2F*_o_ - *F*_c_ maps of a covalent bond between imipenem and S70-OH (**Fig. 3b**). Significantly, imipenem was bound in all eight chains but with three different binding modes. In the C, D and E chains, no Br^-^ anomalous signal was detectable and imipenem was in a similar conformation to that observed in previously published OXA-48 acyl-enzyme intermediate structures (PDB 6P97^41^, 6PTU^32^ and 5QB4^42^)- The backbone amides of S70 and Y211 form an oxyanion hole and are H-bonded to the acyl carbonyl oxygen resulting from S70 attack on imipenem C7 (**Fig. 3b)**. The imipenem carboxylate group forms strong salt bridges with R250 sidechain and an H-bond with T209-OH. Hence the core structure of the imipenem does not block the previously proposed “deacylating water channel”, consisting of L158 and V120^20^, for the acylation step^32,43^. Since this conformation has no bound bromide and is conducive to hydrolysis of the intermediate^44^, we describe it as an *“active intermediate”*. By contrast, in the A, B and H chains, the acyl-intermediate shows imipenem flipped 180° compared to the active intermediate, with its carboxylate group forming a H-bond with R214 and blocking the “deacylating water channel” (**Fig. 3c**). Furthermore, in both A and B chains the orientation of the imipenem carboxylate group leads to disorder and even disruption of the Ω-loop, which is also considered to be detrimental to efficient diacylation^44^ and leads to a “catalytic dead-end”. Moreover, a bromide ion bound to the positive R250 side chain with partial occupancy. Thus, we describe this new bromide-bound structure an *“inactivate intermediate”*. Finally, we observed the dual-occupancy electron density of both intermediates in the F and G chains (**Fig. 3d**), which provided evidence that the inactivate and active intermediates are interconvertible. This was further confirmed in a structure obtained by soaking imipenem into Cl^-^-free OXA-48_AcK73_ crystals, in which we saw both *inactivate* and *active* acyl-intermediates (**Fig. S10**).

**Figure 3.**
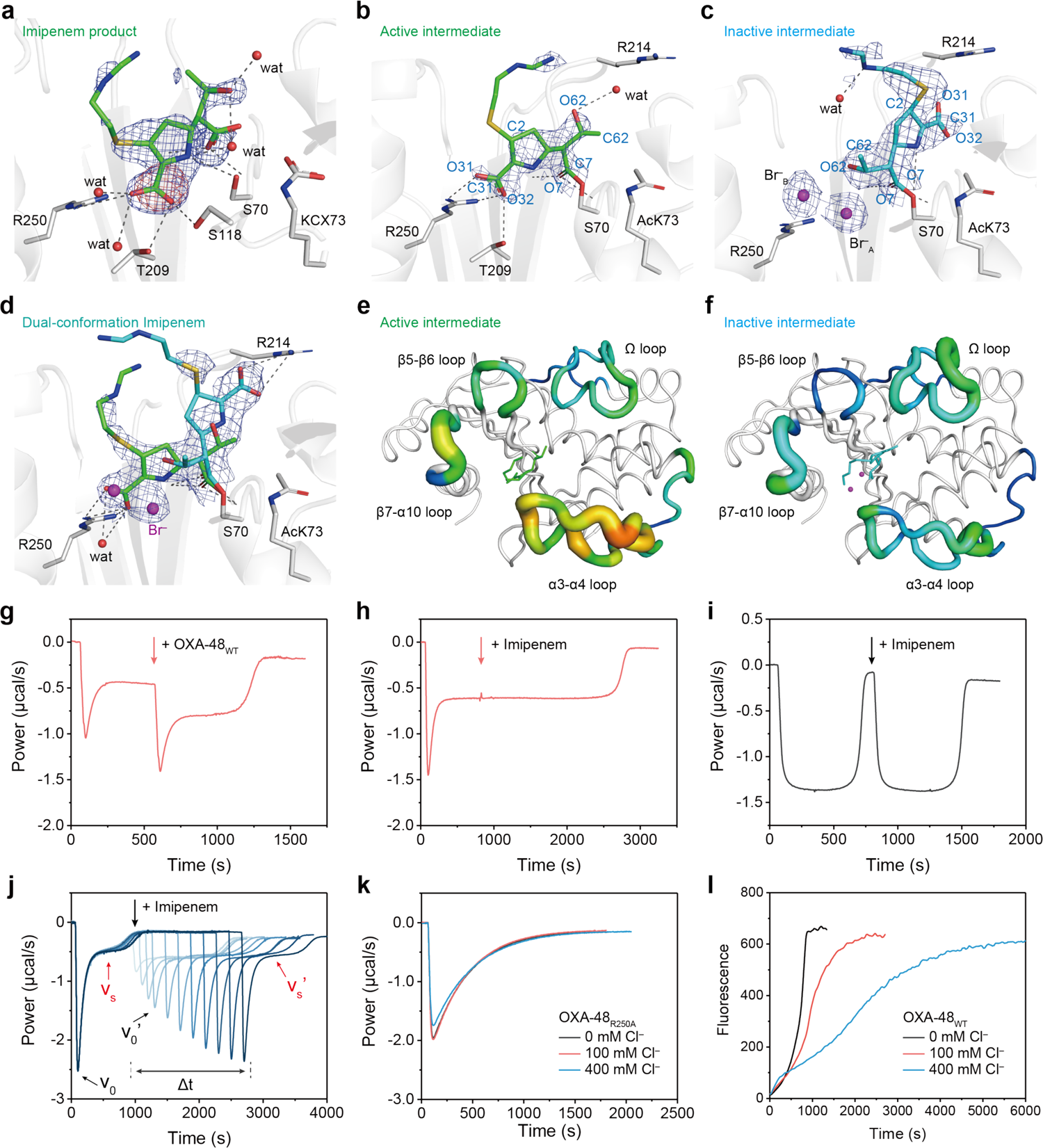
(**a**) Crystal structure of OXA-48wτ-imipenem-Br^-^ product complex obtained by soaking 500 mM NaBr and 32 mM imipenem into the apo OXA-48_WT_ crystal together. This structure contains an anomalous signal for Br (red mesh, contoured at 3σ) overlapped with hydrolyzed imipenem density, showing the active intermediate conformation competes with the halide ion for R250 binding. The initial *F*_o_-*F*_c_ electron density map of hydrolyzed imipenem is shown as blue mesh, contoured at 3σ (0.2687 e^-^/Å^3^). (**b,c**) The crystal stuctures are captured in (b) active intermediate and (c) inactive intermediate states stabilized by bromide in OXA-48_AcK73_. Unbiased 2*F*_o_-*F*_c_ electron density map of imipenem is shown as blue mesh, contoured at 1σ. Occupancies are 0.35 for Br_A_ and 0.4 for Br_B_. (**d**) Crystal structure of OXA-48_AcK73_-imipenem-Br^-^ acyl-intermediate complex with dual-occupancy density from both active and inactive conformations in chain G. The initial unbiased 2*F*_o_-*F*_c_ electron density map of imipenem is shown as blue mesh, contoured at 1σ (0.2330 e^-^/Å ^3^). (**e**) Active intermediate and (**f**) inactive intermediate monomers are drawn in cartoon putty representations for the Ω loop, β5-β6 loop, β7-α10 loop and α3-α4 loop; blue represents the lowest and red the highest B-factor value. The size of the tube reflects the value of the B-factor: the better the *B*-factor is, the thicker the tube is. Structures in other regions are coloured in white and displayed in cartoon tube representation, where the size of the tube is independent of B-factors. (**g**) Effects of adding fresh enzyme or (**h**) fresh substrate to the steady-state phase of the reaction on the biphasic kinetics with 100 mM Cl^-^. **(i**) Two successive additions of 200 μM fresh imipenem into 100 nM OXA-48_WT_ solution in Cl^-^-free buffer. (**j**) Overlay of ITC kinetic curves for titration of the time interval between two additions of 140 μM fresh imipenem into 100 nM OXA-48_WT_ solution with 400 mM Cl^-^. (**k**) Kinetic curves of 200 μM imipenem hydrolysis by 2 μM OXA-48_R250A_ variant with various [Cl^-^]. (**l**) Effect of [Cl^-^] on the intrinsic fluorescence curves of 200 nM OXA-48 hydrolysis of 400 μM imipenem. The fluorescence did not recover to the original intensity as the hydrolysed imipenem quenches the intrinsic fluorescence in a concentration-dependent fashion (shown in **Fig. S16**). All buffers used for ITC and fluorescence assays are 50 mM NaPi pH 7.5 with 1 mM NaHCO_3_.

These two intermediate conformations exhibit marked differences in their dynamic fluctuations near the active site, as presented by their B-factors^45^. The three chains containing the *active intermediate* conformation show higher B-factors in the β5-β6 loop, β7-α10 loop and α3-α4 loop regions around the active site^46^ (**Fig. 3e**), suggesting an association with a higher catalytic activity of OXA-48. By contrast, the three chains containing the *inactive intermediate* show higher B-factors in the Ω loop due to the spatial clashing with the C3 carboxylate group (**Fig. 3f**), and it has been suggested previously that a more disordered Ω loop conformation may decrease deacylation^44^. These results clearly show that *active-inactive* intermediate conformations induced by chloride ions are associated with distinct active site dynamics.

To establish that the inactive conformation observed *in crystallo* depends on the presence of chloride during turnover, we first allowed the reaction to progress to the steady state. At this point, the addition of fresh OXA-48 enzyme reinitiated the reaction with the kinetics of a biphasic calorimetric curve (**Fig. 3g**), whereas adding fresh imipenem did not initiate a biphasic process (**Fig. 3h**), meaning the equilibrium of the two intermediate conformations with different reactivities was established at the steady state. We next carried out double injection ITC experiments to explore the shift in this equilibrium. When there is no chloride in the reaction solution, the calorimetric curves of the two injections are almost identical (**Fig. 3i**), which indicates the turnover capacity of the enzyme remains the same and there is no product inhibition. By contrast, with 100 mM and 400 mM Cl^-^ in the reaction solution, the addition of fresh imipenem results in time-dependent changes in the calorimetric curves. When there is no time interval between the two reactions, the initial rate of the new reaction (*v*_0_’) is approximately the same as the previous steady-state rate *v*_s_ (**Fig. 3j**, light blue line, **Fig. S11**). As the time interval is increased, *v*_0_ of the second reaction gradually increases until it fully recovers to *v*_0_ (**Fig. 3j**, dark blue line), which suggests that the concentration of free enzyme increases with time, leading to apparent enzyme reactivation. This further confirms that chloride-induction of biphasic kinetics is related to a slow redistribution of the equilibrium between the two intermediate conformations with different reactivities.

### R250 plays a key role in halide-induced biphasic kinetics

Active site residue R250 has been proposed to play an important role in catalysis by binding to the carboxylate of the substrates^47^. We have observed clear binding of Br^-^ or I^-^ ion interacting with R250 in our halide-soaked OXA-48_WT_ and OXA-48_AcK73_-imipenem-Br^-^ structures (**Fig. 2f, Fig S12**). However, no electron density for bromide was found near A250 (**Fig. S13**) for the OXA-48_R250A_ variant structure (1.85 Å resolution) following soaking OXA-48_R250A_ apo crystals with 500 mM NaBr. More importantly, the ITC curve of OXA-48_R250A_ variant no longer exhibits biphasic features and does not respond to the presence of chloride (**Fig. 3k**). By comparison, active site R214 interacts with the carboxylate of imipenem in the inactive conformation, but OXA-48_R214A_ variant shows chloride-dependent biphasic kinetics (**Fig. S14**). These data show that halide ion binding to R250 is essential for generating biphasic kinetics during imipenem hydrolysis.

Combining the structural evidence with our mutagenesis data, we propose that for the chloride-free reaction of OXA-48, the *active intermediate* predominates and delivers turnover. After supplementation of chloride, binding of a halide ion wedged between imipenem intermediate and R250 results in progressive rate-retardation from the accumulation of the *inactive intermediate* formed during catalysis until a new *active-inactive* equilibrium is established to deliver the slower, steadystate kinetic behaviour.

### Direct observation of conformational changes by intrinsic fluorescence

To monitor chloride-induced conformational changes of OXA-48 directly in real-time, we used the intrinsic fluorescence of tryptophan, which has previously been employed for detecting substrate binding to NDM-1^48^ and ligand binding to OXA-1^49^. OXA-48 has several tryptophan residues near the active site, including W105, 157, 221, and 222. In particular, W105 on the α3-α4 loop has a direct interaction with imipenem. When we excited OXA-48 solution at λ_ex_ = 280 nm, it showed a stable fluorescence intensity at 340 nm (**Fig. S15**). Upon addition of substrates at *t* = 0, the intrinsic fluorescence of OXA-48 was immediately quenched due to substrate binding (**Fig. 3i**). Under Cl^-^-free conditions, the fluorescence quickly recovered as soon as the substrate was consumed at around 750 s, indicating that protein dynamics are synchronized with substrate binding and turnover (**Fig. 3i**, black curve). In contrast, when chloride was present, corresponding to the burst phase of the ITC curve, we observed a fluorescence recovery initially faster but dramatically slowed down after 500 s until 100 min. This demonstrates, in complementation to ITC, intrinsic fluorescence detects more subtle underlying dynamics from the two intermediate conformations in OXA-48, even after the time when substrate has been completely consumed (reflected by ITC, **Fig. 3j**). It identifies an unexpected long time-lag for OXA-48 to recover fully its original conformation under physiological conditions.

### Chloride is a *Janus Effector*

Based on our above results, we now propose a new mechanism with Cl^-^ as a “positive allosteric effector” and “catalytic inhibitor” to explain the observed biphasic kinetics^50^ (**Fig. 4a**). Chloride ions readily bind at the OXA-48 dimer interface as a positive allosteric effector generating a faster E^Cl^ and the fast formation of an acylated *active intermediate* E^Cl^A at rate constant *k*_1_’. As acylation (at *k*_2_’) is chemically rate-limiting, Cl^-^ ion(s) also compete to bind at R250, driving the acylated intermediate E^Cl^A towards the formation of an *inactive intermediate* E^Cl^A•Cl complex at *k*_3_’. The inactivated E^Cl^A•Cl then reverts to active E^Cl^A conformation in order to re-establish acyl-intermediate hydrolysis at *k*-3’.

**Figure 4.**
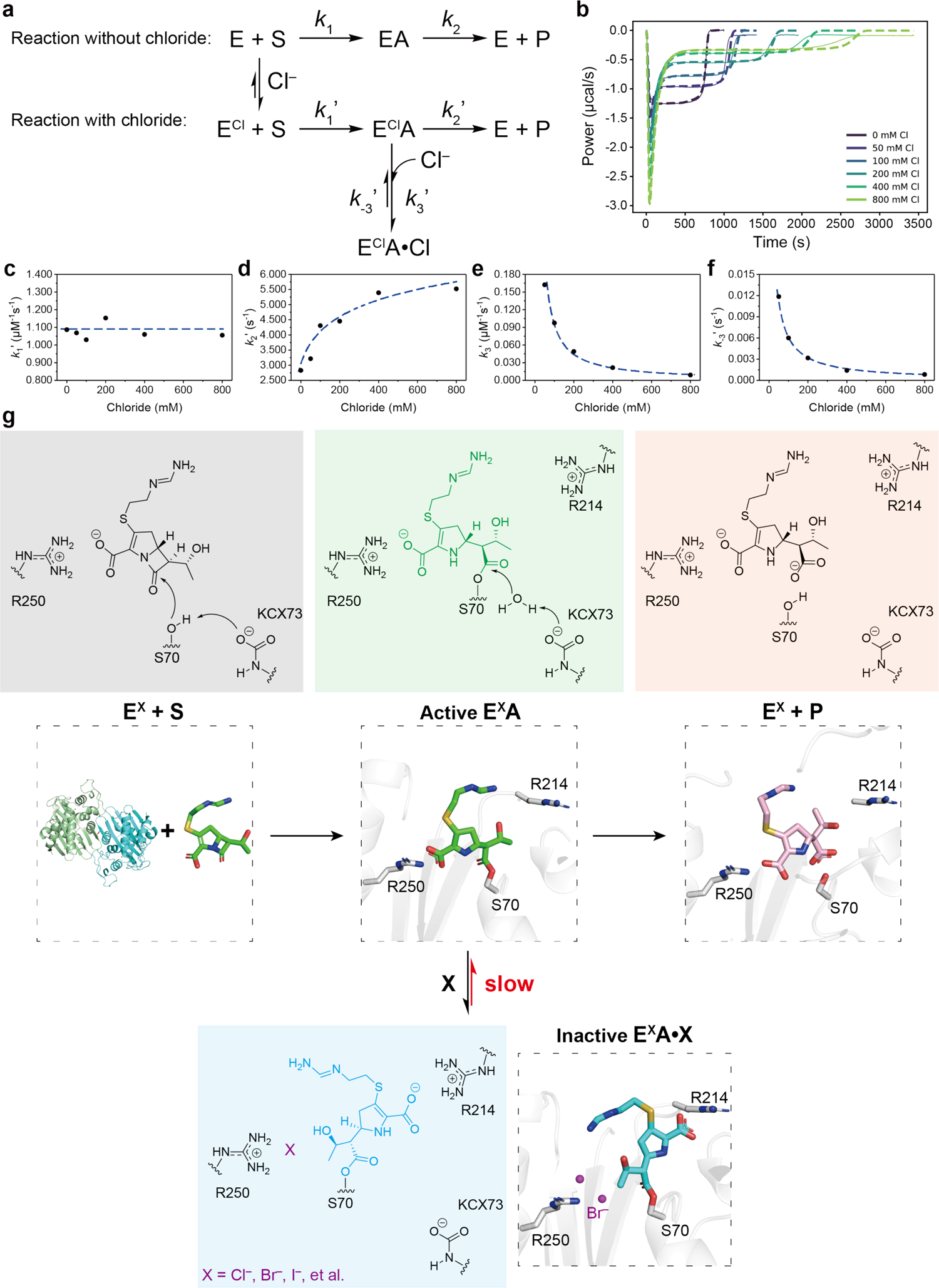
(**a**) Mechanism explaining biphasic hydrolysis with catalytic inhibition by chloride. E, enzyme; S, substrate. E^Cl^, chloride bound at the interface of the OXA-48 dimer; S, substrate; P, product; Active E^Cl^A, acyl-enzyme intermediate; Inactive E^Cl^A•Cl, chloride bound at R250 at an inactive conformation. (**b**) Kinetic modelling and simulation for biphasic hydrolysis. (**c-f**) Apparent rate constants *k*_1_’, *k*_2_’, *k*_3_’, *k*_4_’ and their dependence on [Cl^-^]. (**g**) Model showing biphasic hydrolysis with catalytic inhibition by halide. E, enzyme; S, substrate. E^X^, chloride bound at the interface of the OXA-48 dimer; S, substrate; P, product; Active E^X^A, acyl-enzyme intermediate; Inactive E^X^A•Cl, chloride bound at R250 at an inactive conformation.

With this model mechanism, we used computer analysis to obtain the apparent rates involved in each step of the reaction using our high-quality ITC data from **Fig 4b** and **S4d,** to reveal the step(s) that are [Cl^-^]-dependent (**Figs. S16-21**). Curve fitting shows *k*_1_’ is not correlated to [Cl^-^] (**Fig. 4c** and **Table 1**). This means that chloride does not interfere with substrate binding to the enzyme and therefore a path to E^Cl^A•Cl directly from E and S is unlikely. However, the magnitude of *k*_2_’ for turnover increases with chloride concentration because of a more positive allosteric effect (**Fig. 4d**). Hence, at the burst phase nearly all enzyme is in the active intermediate conformation E^Cl^A. This is followed by a sharp heat reduction during the steady-state reaction, which is due to the conversion of E^Cl^A + Cl into E^Cl^A•Cl at an apparent rate constant of *k*3’, that ranges from 0.16 μM^-1^ s^-1^ at 50 mM Cl^-^ to 0.009 μM^-1^ s^-1^ at 800 mM Cl^-^ (**Fig. 4e**). The apparent rate constant *k*-3’ for conversion of E^Cl^A•Cl into E^Cl^A + Cl is also inversely related to chloride concentration, ranging from 2 x 10^-4^ s^-1^ at 50 mM Cl^-^ to 10^-6^ s^-1^ at 800 mM Cl^-^ (**Fig. 4f**). Our catalytic inhibition model explains why higher [Cl^-^] prolongs the steady-state phase in the biphasic curve. The recovery of enzyme activity from E^Cl^A•Cl to E^Cl^A at *k*_-3_’ is the rate-limiting step, which explains how in intrinsic fluorescence experiments the inactive conformation had not fully transformed into the active conformation until after all substrate had been consumed (**Fig. 3l**). Furthermore, the direct hydrolysis path from inactive E^Cl^A•Cl to product is also shown to have a negligible rate (**Fig. S22**).

**Table 1.** The apparent rate constants for OXA-48_WT_ and OXA-48_E185A/R186A/R206A_.

| | **[CI]**<br>(mM) | **$k_1'$**<br>($\mu\text{M}^{-1} \text{s}^{-1}$) | **$k_2'$**<br>($\text{s}^{-1}$) | **$k_3'$**<br>($\mu\text{M}^{-1} \text{s}^{-1}$) | **$k_{-3}'$**<br>($\text{s}^{-1}$) |
| --- | --- | --- | --- | --- | --- |
| OXA-48<sub>WT</sub> | 0 | 1.09 | 2.83 | - | - |
|  | 100 | 1.03 | 4.35 | 0.11 | 0.0071 |
|  | 400 | 1.15 | 4.47 | 0.026 | 0.0034 |
| OXA-48<sub>E185A/R186A/R206A</sub> | 0 | 0.31 | 4.51 | - | - |
|  | 100 | 0.30 | 4.71 | 0.12 | 0.0163 |
|  | 400 | 0.32 | 3.38 | 0.026 | 0.0046 |

When we compare our derived kinetic parameters for OXA-48_E185A/R186A/R206A_ to OXA-48_WT_ (**Fig. S23, Table 1**), all *k*_1_’ values of OXA-48_WT_ are three times greater than for OXA-48_E185A/R186A/R206A_. This shows the allosteric effect on the surface increases the substrate affinity by three-fold regardless of chloride. This positive allosteric effect propagated from the surface also affects the acylation, reflected by the overall increase in *k*_2_’ values of OXA-48_WT_’ while *k*_2_’ values of OXA-48_E185A/R186A/R206A_ decrease when [Cl^-^] increases. By contrast, both *k*_3_’ and *k*_-3_’ are in the same order of magnitude for OXA-48_WT_ and OXA-48_E185A/R186A/R206A_. This suggests that changes in surface-related allostery have little effect on catalytic inactivation by Cl^-^.

### The novel mechanism is general

To verify the generality of our new mechanism, a series of β-lactam substrates were chosen for kinetic studies with OXA-48, including cefamezin, oxacillin, penicillin and cefotaxime. Our data show conclusively that, unless chloride ions are included in the buffer, there are no biphasic kinetics of OXA-48 with these substrates, even for oxacillin with a considerably large substituent chain group^9,51^ (**Fig. 5a-e**). We performed molecular docking using the H-chain of the OXA-48_AcK73_-imipenem structure containing a bromide ion, using MOE^52^. As expected, oxacillin displays an inactive intermediate binding mode with its α-carbon substituent group on the lactam forming more prominent hydrophobic interactions with residues I102, Y117 and S244 (**Fig. S24, S25**), and similar to imipenem in its inactive intermediate. Oxacillin in the *inactive conformation* (**Fig. S24b**) further blocks the “deacylating water channel”. This is in marked contrast to the binding mode of oxacillin in our active acyl-intermediate-OXA-48-_AcK73_ structure in the absence of halide (**Fig. S24a**). Together, our data confirm that when reacting with other β-lactams, halides also induce biphasic kinetics through interacting with the R250 sidechain to promote the formation of an inactivated acyl-intermediate complex. Therefore, our new model for catalytic inhibition by chloride is likely to be widely represented in OXA-48 hydrolysis of a variety of substrates.

**Figure 5.**
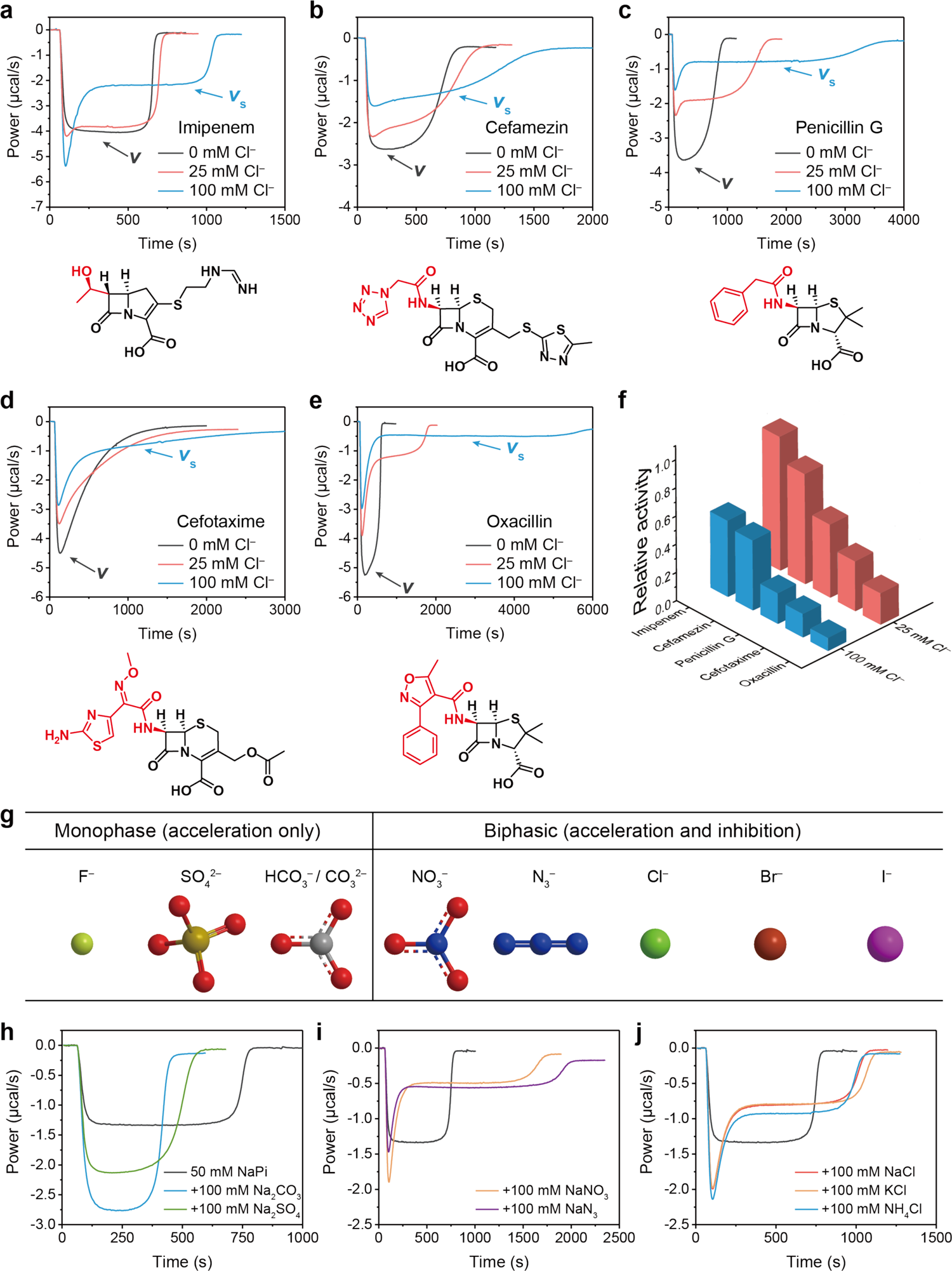
The generality of halide-induced biphasic kinetics. (**a-e**) Kinetic curves for a series of substrates hydrolysis by OXA-48_WT_ with 0, 25 mM or 100 mM Cl^-^. The side chain groups of different antibiotic substrates are shown in red. (**f**) The relative steady-state activities *v_s_/v* of OXA-48_WT_ hydrolyzing different antibiotics in the presence of 25 mM or 100 mM Cl^-^. (**h, i**) Effect of various anions on the kinetic rates. 200 μM imipenem was hydrolyzed by 100 nM OXA-48_WT_ with the supplement of 100 mM various sodium salts. (**j**) Effect of different cations on the kinetic curves. 200 μM imipenem was hydrolyzed by 100 nM OXA-48_WT_ with the supplement of 100 mM chloride salts of sodium, potassium, and ammonium. The black curve is the control without any chloride salt. All buffers used for ITC assays are 50 mM NaPi pH 7.5 with 1 mM NaHCO_3_.

## Discussion

The current ineffectiveness of inhibitors for Class D carbapenemases is a result of their extraordinary ability to mutate and expand their spectrum of action^53^. The investigation of the catalytic mechanism of carbapenemases under physiological conditions is important for the development of novel antibiotics and detection methods to tackle antibiotic resistance. However, the molecular mechanism related to inactivation during catalysis^9,54^ of the highly clinically important OXA-48-like enzymes is still unresolved. It is imperative to understand how these carbapenemases work in order to design useful inhibitors and specific detection assays^8,37,55^.

Hitherto, the observed biphasic kinetics have been attributed to several different causes. One is that β-lactams with sterically encumbered side chains at their a-carbon, such as cloxacillin, oxacillin and carbenicillin, can induce a conformational change in OXA-10^9^, OXA-27^14^, and OXA-2^11^, as well as in some other β-lactamases^51^. Moreover, in the protein dimer/monomer equilibrium, established for OXA-14^12^, OXA-10^56^ and OXA-48^40^, the dimer is regarded as the more active form whose concentration-dependency is mediated by cations. By testing a series of substrates such as oxacillin with sterically encumbered side chains at α-carbon (**Fig. 5a-e**) and comparing both the dimeric OXA-48_WT_ and monomeric OXA-48_E185/R186/R206_ variant in the presence/absence of halides (**Fig. 2g**), our analysis of chloride-induced biphasic kinetics of OXA-48 provides a complete molecular mechanism to explain this peculiar kinetical behaviour. Our analysis draws attention to re-evaluating the influence of these mechanistic components on the efficacy of antibiotics. For example, therapeutic injection of the antibiotic ceftazidime uses Na_2_CO_3_ at a concentration of 11.8% w/w to facilitate dissolution, with NaCl solution often as the preferred diluent. Given that chloride is common in physiological and experimental conditions, our study also underlines the need to pay careful attention while studying OXA enzymes in order to avoid inconsistent results resulting from variation in chloride concentrations in the buffer solution and in cell assays, including Michaelis-Menten kinetics measurements and MIC assays.

We show that the change in the state of catalytic lysine carbamylation is not the origin of biphasic kinetics during turnover by comparing thermodynamic profiles of OXA-48_WT_ and chemically stable OXA-48_AcK73_ with a series of β-lactam antibiotics at neutral pH. We also solved a 1.60 Å co-crystallized OXA-48_WT_-avibactam-Cl^-^ structure under neutral pH conditions with decarbamylated K73 (**Fig. S26**). This further proves that avibactam is not a good acyl-intermediate mimic for studying the behaviour of decarbamylation in Class D β-lactamases. While we show that chloride ion binding to the enzyme dimer interface stabilizes its quaternary structure, more significantly, chloride interacts with the active site R250 side chain leading directly to an *inactivated* form of the acyl intermediate via a “door wedge” mode, thereby accounts for the retarded rate in the observed biphasic kinetics.

Our kinetic analysis further shows that HCO_3_^-^ is a weak but positive allosteric effector. That is supported by an OXA-48_WT_-HCO_3_^-^ structure in which HCO_3_^-^interacts with R206-R206’ at the dimer interface. This structure also shows how HCO_3_^-^ interacts with R250 in the active site (**Fig. 2j**). However, unlike Cl^-^, Br^-^, and I^-^ ions, bicarbonate cannot act like a wedge between R250 and the inactive acyl-intermediate for catalytic inhibition, given that HCO_3_^-^ is much bigger than the halide anions and would clash with the intermediate (**Fig. S27**). Thus, numerous deductions from activity rescue experiments with NaHCO_3_ assuming KCX73 was prevented from decarbamylation in many early studies now need to be re-evaluated. As for HCO_3_^-^, we also showed that SO4^2-^ is exclusively a positive allosteric effector with no inhibitory effect, whereas nitrate and azide induce biphasic kinetics similar to those for chloride (**Fig. 5g-j**). These differences may well be due to variable energy of desolvation that each ionic species has to overcome in binding to the protein, and which could significantly affect their ability to act as a wedge^57^. Cations, *e.g*., Na^+^, K^+^ and NH_4_+, do not have any observed effect on the kinetics (**Fig. 5j, Fig. S28**). We also showed by ^13^C NMR, using ^13^C-NaHCO_3_ labelled OXA-48_WT_, that addition of 50 mM or more Cl^-^ does not cause the KCX or N-terminus decarbamylation, but can change the chemical shift of KCX (**Fig S29**).

Our computer analysis identifies *k*_-3_’ to be an order of magnitude slower than *k*_3_’ and to be [Cl^-^]-dependent (**Fig. 4e, f**). This chloride-dependent attenuation of reactivity by Cl wedging in place against the conserved residue R250 raises a puzzling question: Why do some bacteria encode Class D carbapenemase OXAs when that appears to put them at a competitive disadvantage by keeping R250 in the active site? Perhaps this is because of a trade-off between carbapenem substrate binding and chloride’s catalytic inhibition, during a relatively short evolutionary period of OXAs since the use of synthetic β-lactam antibiotics, especially carbapenems. In that regard, we note that the environmental *Shewanella xiamenensis* that has been proposed to carry the *bla*OXA-48 gene was only discovered early this century^8,58,59^.

It is well-established that metal cations influence the structure and function of metalloenzymes. However, the influence of anions on enzyme catalysis has been relatively little investigated^60^. Of only a few examples, chloride binding to a-amylase has been shown to promote catalytic activity by interacting with active site carboxyl groups^60^. Our present results show chloride is a “Janus effector” of OXA-48: It manifests allosteric activation, by binding at the dimer interface, to be followed by catalytic inhibition, through mis-orientation of a reaction intermediate in the active site. This finding greatly widens our understanding of the role of anions, particularly halides, in modulating enzyme catalysis.

In conclusion, our study identifies chloride as trigger for the biphasic kinetics of OXA-48 with a series of β-lactam substrates. It elucidates the binding site of halides on OXA-48, and it reveals the molecular mechanism of their effect on β-lactam hydrolysis. This finding is informative for the future design of new halide-tolerant antibiotics capable of preserving the effectiveness of existing antibiotics. The identification of a significant role for the highly conserved R250 in chloride binding (**Fig. S30, Table S5**) may also provide an additional target site for designing new antimicrobial agents. Moreover, halide ion-induced biphasic kinetics is a typical feature of carbapenemase OXAs and may be employed to develop new assays for highly specific detection of the carbapenemase-producing bacteria.

## Acknowledgement

We thank Dr. Pierre Rizkallah from Cardiff University Medical School for assistance with the X-ray data collection, the Diamond Light Source for access to beamline I03 under proposal number mx20147, and Shanghai Synchrotron Radiation Facility for access to beamline BL17B1 at National Facility for Protein Science in Shanghai (NFPS). G.M.B thanks the Leverhulme Trust for an Emeritus Fellowship (EM-2022-057). We thank Northwest University for a visiting studentship for Z.Z. and EPSRC for a studentship for D.Z. P.C. was supported by the Ministerio de Ciencia, Innovación y Universidades/FEDER (Spain/UE) (PGC2018-098186-B-I00 (BASIC) and PID2019-109320GB-I00). Y.H. was supported by the National Natural Science Foundation of China (31400663), Shaanxi Provincial Department of Education Funds (22JP081) and the Open Fund of Engineering Research Center of Artificial Organs and Materials, Jinan University (ERCAOM202210). We thank Wellcome Trust Seed Award (209057/Z/17/Z) for supporting Y.J. and H.B., and The Academy of Medical Sciences Springboard Award (SBE003\1154) for supporting Y.J. and P.B. Y.J. is a Wellcome Trust Sir Henry Dale Fellow (218568/Z/19/Z).

## Author contributions

Q.Z., Y.H. and Y.J. drafted the manuscript with contributions from all authors. Y.H. and Y.J. coordinated the study. Y.H. and Y.J. designed experiments and analyzed the data. Q.Z. and Y.Z. performed biochemical and microbiological experiments. Q.Z., and Y.Z. performed ITC and UV-Vis assays. H.B., and Q.Z. performed mutagenesis of OXA-48. H.B. performed acetyllysine in cooperation. R.E. and D.Z. performed the synthesis of acetyllysine. H.B., P.B. and J.Y. and Y.J. prepared the proteins and performed NMR analysis. Q.Z., H.B., P.B., Z.Z. and Y.J. performed crystallography and determined the structures. Q.Z. performed bioinformatic analyses and molecular docking. P.C. performed mathematical simulation.

## Footnotes

† This is K73 for OXA-48. The numbering can be variable for different OXAs.

