## Supplementary Information for "Discovery and molecular basis of chloride as an allosteric activator and catalytic inhibitor for Class-D β-lactamases"

**Table of Tables:**

**Table of Figures:**

**Supplementary Figure 20** The analytical approximations are good fits to the data at different chloride concentrations. Solid lines represent experimental data, while dashed lines are simulations of the model. 28
**Supplementary Figure 25** Ligplot diagrams show the hydrogen bonding and hydrophobic interactions between OXA-$48_{AcK73}$ with the inactive acyl-intermediate conformations of (**a**) oxacillin and (**b**) imipenem. 31

**Supplementary Table 1** Survey of all OXA-48 structures in Protein Data Bank (PDB) (up to November 2022). The correlation of the pH, resolution, the carbamylation status of the Lys73, and any presence of halide ions.

| Name | | PDB | Resolution (Å) | pH | Lysine73 carbarmylated (KCX) | Any or position of halide ions |
|---|---|---|---|---|---|---|
| Apo | | 6P96 | 1.6 | ~4 | no | K73, R174, R206 |
| | | 3HBR | 1.9 | 7.5 | yes | none |
| | | 4S2P | 1.7 | 7.5 | yes | none |
| | $^{19}$F-labelled | 6RJ7 | 1.73 | 8 | yes | R206 |
| | S70A | 5HAP | 1.89 | 8.5 | yes | R206 |
| | designed loop18 | 6HOO | 2.38 | | yes | K94 |
| | S70G | 5HAQ | 2.14 | | yes | none |
| | R189A | 6GOA | 2.55 | 9.5 | yes | R206 |
| | | 5OFT | 3.20 | 9 | yes | none |
| | Homocitrullinated | 7LXG | 2.20 | 5.5 | Homocitrulline | R206 |
| | | 6I5D | 1.75 | 7 | no | R206 |
| Imipenem | | 7KH9 | 2.29 | 8.5 | K73A | W157, R189, R206 |
| | | 6P97 | 1.8 | ~4 | no | K73, R174, R206 |
| | | 6PK0 | 1.75 | 4.6 | yes | R206 |
| | | 5QB4 | 1.95 | 7.5 | yes | R206 |
| | | 6PTU | 2.00 | 4.6 | no | K73, R206 |
| | K73A | 7KH9 | 2.29 | 8.5 | K73A | W157, R189, R206 |
| Avibactam | | 4S2J | 2.54 | 6.5 | no | none |
| | P68A | 6Q5B | 2.22 | 6.5 | partially | Q91, R186, R206 |
| | | 4S2K | 2.1 | 7.5 | no | none |
| | | 4WMC | 2.3 | 7.5 | no | none |
| | | 4S2N | 2.0 | 8.5 | partially | none |
| Other antibiotics | Meropenem | 6P98 | 1.75 | ~4 | no | K73, R174, R186, R206 |
| | Ertapenem | 6P99 | 2.25 | ~4 | no | K73, R206 |
| | Doripenem | 6P9C | 1.9 | ~4 | no | K73, R174, R206, |
| | Cefotaxime | 6PQI | 2.05 | 4.6 | no | K73, R206 |
| | Faropenem | 6PSG | 2.13 | 4.6 | no | R206 |
| | Meropenem | 6PT1 | 2.00 | 4.6 | no | K73, R206 |
| | Cefoxitin | 6PT5 | 2.30 | 4.6 | no | R206 |
| | Ceftazidime | 6Q5F | 2.5 | 6.5-7.5 | partially | Q91, W157, R186, R206 |
| | K73A Meropenem | 7KHQ | 2.00 | | K73A | R174, R206, R214 |
| | K73A Doripenem | 6PXX | 1.50 | 7.5-8.5 | K73A | W157, R206 |
| | L67F Ceftazidime | 7ASS | 1.91 | 9 | yes | R206, R214, R250 |
| | Ertapenem | 6ZRJ | 1.94 | 7.5 | partially | R206 |
| | Meropenem | 6ZRP | 1.74 | 7.5 | partially | R206 |
| Other ligands | | 7AW5 | 1.65 | 7.5 | yes | R206 |
| | | 7AUX | 2.05 | 7.5 | yes | R206 |
| | | 6V1O | 1.80 | 8.5 | yes | R206 |
| | | 7JHQ | 2.00 | 8.5 | yes | none |
| | | 6XQR | 2.20 | 8.5 | yes | R206 |
| | | 7K5V | 2.80 Å | 8.5 | yes | R206 |
| | | 7L8O | 2.70 | 8.5 | yes | R186, Q193, R206, K208 |
| | | 7R6Z | 2.10 | 8.5 | yes | S40, R206 |
| | | 5DTS | 1.94 | 7.5 | yes | R206 |
| | | 5DTK | 1.60 | 7.5 | yes | R206 |
| | | 5DTT | 2.10 | 7.5 | yes | R206 |

| | | | | | | |
|---|---|---|---|---|---|---|
| | | 5DVA | 2.50 | 7.5 | yes | R206 |
| | | 6UVK | 2.20 | 8 | yes | N110, R206 |
| | | 7DML | 1.94 | 7.5 | yes | none |
| | | 6ZXI | 1.85 | 7.5 | no | R206 |
| | | 5QA4 | 1.95 | 7.5 | yes | R174, R206 |
| | | 5QA5 | 1.95 | 7.5 | yes | Y177, R206 |
| | | 5QA6 | 1.95 | 7.5 | yes | R206 |
| | | 5QA7 | 1.82 | 7.5 | yes | R206 |
| | | 5QA8 | 2.50 | 7.5 | yes | R206 |
| | | 5QA9 | 1.90 | 7.5 | yes | R206 |
| | | 5QAA | 1.95 | 7.5 | yes | R206 |
| | | 5QAB | 2.15 | 7.5 | yes | R206 |
| | | 5QAC | 2.00 | 7.5 | yes | R206 |
| | | 5QAD | 1.75 | 7.5 | yes | R206 |
| | | 5QAE | 2.10 | 7.5 | yes | R206 |
| | | 5QAF | 2.15 | 7.5 | yes | R206 |
| | | 5QAG | 2.30 | 7.5 | yes | R206 |
| | | 5QAH | 1.95 | 7.5 | yes | R206 |
| | | 5QAI | 1.90 | 7.5 | yes | R206 |
| | | 5QAJ | 2.00 | 7.5 | yes | R206 |
| | | 5QAK | 1.90 | 7.5 | yes | R206 |
| | | 5QAL | 1.95 | 7.5 | yes | R206 |
| | | 5QAM | 1.87 | 7.5 | yes | R206 |
| | | 5QAN | 2.30 | 7.5 | yes | R206 |
| | | 5QAO | 2.00 | 7.5 | yes | R206 |
| | | 5QAP | 1.79 | 7.5 | yes | R206 |
| | | 5QAQ | 2.40 | 7.5 | yes | R206 |
| | | 5QAR | 2.10 | 7.5 | yes | R206 |
| | | 5QAS | 1.90 | 7.5 | yes | none |
| | | 5QAT | 1.90 | 7.5 | yes | R206 |
| | | 5QAU | 1.75 | 7.5 | yes | R206 |
| | | 5QAV | 1.72 | 7.5 | yes | R206 |
| | | 5QAW | 2.20 | 7.5 | yes | R206 |
| | | 5QAX | 2.31 | 7.5 | yes | R206 |
| | | 5QAY | 1.70 | 7.5 | yes | R206 |
| | | 5QAZ | 2.20 | 7.5 | yes | R206 |
| | | 5QB0 | 1.95 | 7.5 | yes | R206 |
| | | 5QB1 | 1.80 | 7.5 | yes | R206 |
| | | 5QB2 | 1.75 | 7.5 | yes | R206 |
| | | 5QB3 | 2.00 | 7.5 | yes | R206 |
| | | 5FAQ | 1.96 | 7.5 | no | R206 |
| | | 5FAS | 1.74 | 7.5 | no | R206 |
| | | 5FAT | 2.09 | 7.5 | no | R206 |

**Supplementary Table 2** $EC_{50}$ and $IC_{50}$ Values of OXA-48 for all four halide ions.

| | **$EC_{50}$ (mM)[a]** | **$IC_{50}$ (mM)[b]** |
|---|---|---|
| **$F^-$** | >222 | ND[c] |
| **$Cl^-$** | 259 ± 29 | 24 ± 2 |
| **$Br^-$** | >253 | 2.9 ± 0.3 |
| **$I^-$** | ND[c] | 2.25 ± 0.03 |

[a] Determined with the initial burst rate $v_0$. [b] Determined with the steady-state rate $v_s$. [c] The initial rate acceleration phase for $I^-$ and the inhibitory phase for $F^-$ are not significant enough to be measured accurately.

**Supplementary Table 3** Data collection and refinement statistics

| | OXA-48$_{WT}$ apo | OXA-48$_{WT}$-$Br^{-}$ | OXA-48$_{WT}$-$I^{-}$ | OXA-48$_{E185A/R186A/R206A}$-$Cl^{-}$ | OXA-48$_{R250A}$-$Br^{-}$ |
|---|---|---|---|---|---|
| **PDB** | **7PEH** | **7O5T** | **7NRJ** | **7PEI** | **7PGO** |
| **Data collection** | | | | | |
| Wavelength (Å) | 0.815342 | 0.918374 | 0.979150 | 0.979150 | 0.918395 |
| Space group | P 21 21 21 | P 21 21 21 | P 1 21 1 | P 62 | P 21 21 21 |
| Cell dimensions | | | | | |
| *a*, *b*, *c* (Å) | 83.00, 106.98, 124.61 | 82.57, 107.26, 124.62 | 46.76, 126.86, 111.10 | 146.19, 146.19, 54.89 | 84.55, 108.20, 124.61 |
| α, β, γ (°) | 90.00, 90.00, 90.00 | 90.00, 90.00, 90.00 | 90.00, 98.35, 90.00 | 90.00, 90.00, 120.00 | 90.00, 90.00, 90.00 |
| Resolution (Å) | 53.84-1.92 (1.96-1.92) | 81.30-1.81 (1.84-1.81) | 19.78-1.67 (1.70-1.67) | 19.69-1.90 (1.94-1.90) | 58.75-1.85 (1.88-1.85) |
| $R_{sym}$ or $R_{merge}$ (within I+/I-) | 0.313 (2.917) | 0.193 (2.259) | 0.088 (1.847) | 0.125 (4.749) | 0.381 (3.815) |
| $R_{sym}$ or $R_{merge}$ (all I+ and I-) | 0.324 (3.030) | 0.199 (2.347) | 0.095 (1.885) | 0.127 (4.858) | 0.388 (3.890) |
| $I$ / σ$I$ | 5.8 (1.0) | 7.4 (1.0) | 11.0 (1.0) | 16.1 (1.0) | 6.6 (1.1) |
| Completeness (%) | 99.9 (99.8) | 99.9 (99.7) | 99.9 (99.9) | 99.9 (100.0) | 100.0 (99.8) |
| Multiplicity | 12.0 (11.4) | 13.3 (12.6) | 6.8 (6.9) | 20.6 (20.5) | 23.2 (23.7) |
| **Refinement** | | | | | |
| Resolution (Å) | 53.90-1.92 | 81.43-1.81 | 19.74-1.67 | 19.70-1.90 | 58.82-1.85 |
| No. reflections all / free | 85147 / 4389 | 101119 / 5098 | 147791 / 7450 | 52978 / 2657 | 97955 / 5022 |
| $R_{work}$ / $R_{free}$ | 0.218 / 0.250 | 0.187 / 0.261 | 0.166 / 0.197 | 0.177 / 0.211 | 0.205 / 0.234 |
| No. atoms | | | | | |
| Protein | 7907 | 8009 | 7958 | 3944 | 7978 |
| Ligand / ion | 83 / | 54 / 7 | 119 / 13 | 73 / | 72 / 3 |
| Water | 455 | 709 | 923 | 258 | 902 |
| *B*-factors | | | | | |
| Protein | 25.46 | 28.86 | 24.98 | 40.39 | 21.76 |
| Ligand / ion | 28.77 / | 27.45 / 39.68 | 31.43 / 34.46 | 42.44 / | 22.48 / 28.92 |
| Water | 26.63 | 36.28 | 37.83 | 46.82 | 28.28 |
| R.m.s. deviations | | | | | |
| Bond lengths (Å) | 0.0075 | 0.0099 | 0.0098 | 0.0089 | 0.0088 |
| Bond angles (°) | 1.415 | 1.490 | 1.602 | 1.583 | 1.435 |

*Values in parentheses are for highest-resolution shell.

| | OXA-48$_{AcK}$-imipenem-Br$^{-}$ | OXA-48$_{AcK}$-imipenem | OXA-48$_{AcK}$-oxaciline | OXA-48$_{WT}$-imipenem-Br$^{-}$ | OXA-48$_{WT}$-$HCO_3^{-}$ |
|---|---|---|---|---|---|
| **PDB** | **7Q14** | **7PFN** | **7PSE** | **Not deposited** | **7O9N** |
| **Data collection** | | | | | |
| Wavelength (Å) | 0.976269 | 0.976292 | 0.976270 | 0.918381 | 0.976254 |
| Space group | P 1 21 1 | P 2 21 21 | P 2 21 21 | P 21 21 21 | P 21 21 21 |
| Cell dimensions | | | | | |
| *a*, *b*, *c* (Å) | 63.40, 161.93, 107.78 | 44.80, 105.00, 124.75 | 44.83, 105.48, 124.96 | 81.31, 105.97, 124.68 | 81.81, 106.40, 124.71 |
| α, β, γ (°) | 90.00, 90.51, 90.00 | 90.00, 90.00, 90.00 | 90.00, 90.00, 90.000 | 90.00, 90.00, 90.00 | 90.00, 90.00, 90.000 |
| Resolution (Å) | 80.97-2.15 (2.19-2.15) | 62.45-1.80 (1.84-1.80) | 53.76-2.32 (2.40-2.32) | 80.74-1.53 (1.56-1.53) | 81.81-1.97 (2.01-1.97) |
| $R_{sym}$ or $R_{merge\ (within\ I+/I-)}$ | 0.165 (1.628) | 0.172 (2.448) | 0.262 (2.266) | 0.132 (2.755) | 0.340 (2.727) |
| $R_{sym}$ or $R_{merge\ (all\ I+\ and\ I-)}$ | 0.181 (1.747) | 0.179 (2.545) | 0.276 (2.421) | 0.139 (2.854) | 0.351 (2.820) |
| $I / \sigma I$ | 7.3 (1.1) | 9.9 (1.1) | 7.0 (1.0) | 12.0 (1.1) | 6.0 (1.0) |
| Completeness (%) | 99.6 (99.1) | 100.0 (100.0) | 99.3 (95.3) | 98.3 (96.9) | 100.0 (100.0) |
| Multiplicity | 6.7 (6.7) | 11.4 (11.5) | 10.9 (8.8) | 13.6 (14.1) | 13.2 (13.6) |
| **Refinement** | | | | | |
| Resolution (Å) | 81.10-2.15 | 62.45-1.80 | 53.81-2.32 | 62.42-1.53 | 68.50-1.97 |
| No. reflections all / free | 117142 / 5895 | 55487 / 2745 | 26280 / 1266 | 159336 / 7990 | 77539 / 3748 |
| $R_{work}$ / $R_{free}$ | 0.210 / 0.250 | 0.178 / 0.214 | 0.235 / 0.289 | 0.147 / 0.199 | 0.195 / 0.286 |
| No. atoms | | | | | |
| Protein | 15708 | 3976 | 3934 | 8113 | 8005 |
| Ligand / ion | 372 / 4 | 94 / | 100 / 1 | 124 / 7 | 68 / |
| Water | 518 | 365 | 64 | 1020 | 616 |
| *B*-factors | | | | | |
| Protein | 45.90 | 26.51 | 50.37 | 21.46 | 27.15 |
| Ligand / ion | 42.48 / 37.13 | 32.94 / | 41.91 / 27.56 | 28.77 / 36.17 | 24.28 / |
| Water | 35.38 | 33.15 | 34.31 | 32.38 | 32.01 |
| R.m.s. deviations | | | | | |
| Bond lengths (Å) | 0.0081 | 0.0096 | 0.0075 | 0.0091 | 0.0090 |
| Bond angles (°) | 1.464 | 1.539 | 1.529 | 1.480 | 1.466 |

*Values in parentheses are for highest-resolution shell.

| | OXA-$48_{WT}$-imipenem *product* | OXA-$48_{WT}$-imipenem *intermediate* | OXA-$48_{WT}$-avibactam-$Cl^{-}$ |
|---|---|---|---|
| **PDB** | **7PEP** | **7PSF** | **7O5N** |
| **Data collection** | | | |
| Wavelength (Å) | 0.918381 | 0.918380 | 0.979191 |
| Space group | P 21 21 21 | P 21 21 21 | P 21 21 21 |
| Cell dimensions | | | |
| *a*, *b*, *c* (Å) | 83.95, 107.93, 124.42 | 83.42, 107.60, 124.59 | 89.75, 107.96, 125.68 |
| α, β, γ (°) | 90.00, 90.00, 90.00 | 90.00, 90.00, 90.00 | 90.00, 90.00, 90.000 |
| Resolution (Å) | 69.59-1.70 (1.73-1.70) | 49.39-2.10 (2.15-2.10) | 81.89-1.60 (1.63-1.60) |
| $R_{sym}$ or $R_{merge\ (within\ I+/I-)}$ | 0.170 (2.436) | 0.282 (3.195) | 0.090 (2.665) |
| $R_{sym}$ or $R_{merge\ (all\ I+\ and\ I-)}$ | 0.177 (2.535) | 0.294 (3.331) | 0.092 (2.726) |
| $I$ / σ$I$ | 9.8 (1.0) | 8.2 (1.0) | 12.1 (1.2) |
| Completeness (%) | 100.0 (99.7) | 99.9 (99.9) | 99.9 (99.99) |
| Multiplicity | 13.3 (12.7) | 13.6 (14.0) | 13.2 (13.5) |
| **Refinement** | | | |
| Resolution (Å) | 69.69-1.70 | 49.44-2.10 | 73.14-1.60 |
| No. reflections all / free | 124488 / 6286 | 66012 / 3258 | 160736 / 7894 |
| $R_{work}$ / $R_{free}$ | 0.199 / 0.231 | 0.241 / 0.278 | 0.188 / 0.216 |
| No. atoms | | | |
| Protein | 8033 | 7932 | 8043 |
| Ligand / ion | 213 / 2 | 106 / | 123 / 2 |
| Water | 881 | 231 | 695 |
| *B*-factors | | | |
| Protein | 24.58 | 39.36 | 34.9 |
| Ligand / ion | 33.47 /43.84 | 39.56 | 41.07 / 27.47 |
| Water | 32.61 | 32.64 | 45.33 |
| R.m.s. deviations | | | |
| Bond lengths (Å) | 0.0089 | 0.0089 | 0.0098 |
| Bond angles (°) | 1.518 | 1.523 | 1.589 |

*Values in parentheses are for highest-resolution shell.

**Supplementary Table 4** $T_m$ value of OXA-$48_{WT}$ and OXA-$48_{E185A/R186A/R206}$ with chloride.

| | $[Cl^-]$ (mM) | Tm (°C) | | | Mean ± SD |
|---|---|---|---|---|---|
| OXA-$48_{WT}$ | 0 | 54.91 | 55.06 | 55.24 | 55.1 ± 0.2 |
| | 40 | 57.45 | 58.30 | 58.33 | 58.0 ± 0.4 |
| | 100 | 58.10 | 58.69 | 58.58 | 58.5 ± 0.3 |
| OXA-$48_{E185A/R186A/R206}$ | 0 | 44.86 | 44.16 | 44.33 | 44.5 ± 0.4 |
| | 40 | 44.53 | 44.26 | 44.27 | 44.4 ± 0.1 |
| | 100 | 45.15 | 44.45 | 44.62 | 44.7 ± 0.4 |

**Supplementary Table 5** Sequence alignment for OXAs that have been reported to display biphasic kinetics

| | Enzyme | Type | Kinetics (Substrate) | | Inhibited by halide | Sequence (Align to OXA-48) | | | |
|---|---|---|---|---|---|---|---|---|---|
| | | | | | | **70-73** | **250** | **214** | **GenBank ID** |
| 1 | OXA-48 | CHDL | biphasic | imipenem<br>... | Y | STFK | R | R | >API82700.1 OXA-48 (plasmid) [*Klebsiella pneumoniae*] |
| 2 | OXA-22 | Narrow spectrum | biphasic | cephalosporins<br>penicillins<br>... | Unknown | STFK | R | - | >QCO89828.1 OXA-2 (plasmid) [*Escherichia coli*] |
| 3 | OXA-103 | Narrow spectrum | biphasic | ampicillin<br>carbenicillin<br>oxacillin<br>cloxacillin<br>cephaloridine | Y | STFK | R | - | >ACI28891.1 OXA-10 (plasmid) [*Pseudomonas aeruginosa*] |
| 4 | OXA-144 | Expanded spectrum | biphasic | oxacillin<br>penicillin G<br>cephaloridine<br>carbenicillin | Y | STFK | R | - | >WP_064056056.1 OXA-10 family oxacillin-hydrolyzing class D extended-spectrum $\beta$-lactamase OXA-14 [*Pseudomonas aeruginosa*] |
| 5 | OXA-165 | Expanded spectrum | biphasic | penicillin G<br>ampicillin<br>carbenicillin<br>oxacillin<br>cloxacillin<br>cephaloridine<br>cefotaxime | Unknown | STFK | R | - | >AAB97924.1 $\beta$-lactamase OXA-16, partial [*Pseudomonas aeruginosa*] |
| 6 | OXA-276 | CHDL | biphasic | oxacillin | Y | STFK | R | - | >AAG35609.2 $\beta$-lactamase OXA-27 [*Acinetobacter baumannii*] |
| 7 | OXA-507 | Narrow spectrum | biphasic | ampicillin<br>cefsulodin<br>piperacillin | Y | ST**Y**K | R | - | >WP_225025027.1 OXA-50 family oxacillin-hydrolyzing class D $\beta$-lactamase [*Pseudomonas aeruginosa*] |

| | | | | | | | | | |
|---|---|---|---|---|---|---|---|---|---|
| 8 | OXA-24 | CHDL | | | Y | STFK | R | - | >ACV72170.1 OXA-24 class D β-lactamase (plasmid) [*Acinetobacter baumannii*] |
| 9 | OXA-25 | CHDL | | | Y | STFK | R | - | >AAG35607.1 beta-lactamase OXA-25 [*Acinetobacter baumannii*] |
| 10 | OXA-26 | CHDL | | | Y | STFK | R | - | >AAG35608.1 β-lactamase OXA-26 [*Acinetobacter baumannii*] |
| 11 | OXA-40 | CHDL | | | N | STFK | R | - | >ADB28893.1 class D β-lactamase OXA-40 [*Acinetobacter baumannii*] |
| 12 | OXA-58 | CHDL | | | Y | STFK | R | - | >AAW57529.1 OXA-58 [*Acinetobacter baumannii*] |
| 13 | OXA-163 | Expanded spectrum | | | Y | STFK | R | - | >ADY06444.1 β-lactamase OXA-163 [*Enterobacter cloacae*] |

CHDL - Carbapenem-hydrolyzing class D β-lactamase

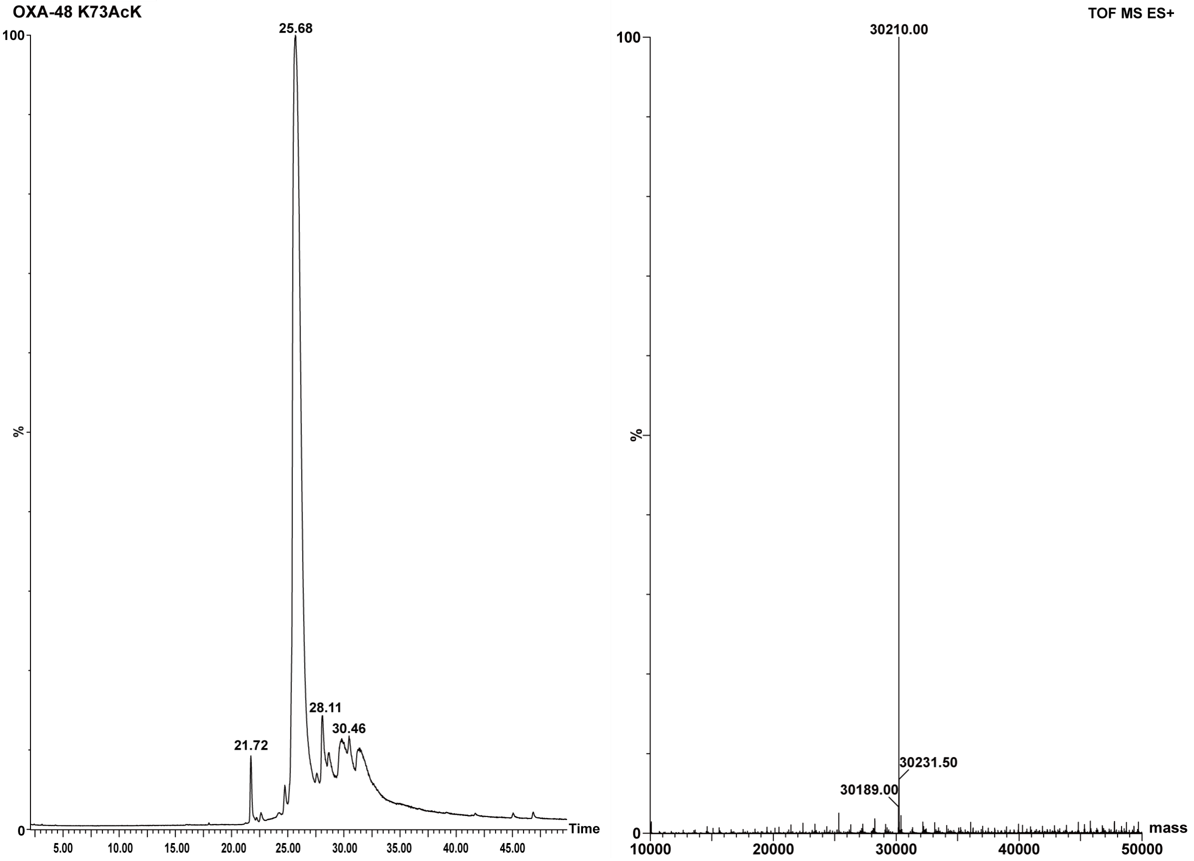

**Supplementary Figure 1** Mass spectrometry of OXA-48AcK73.

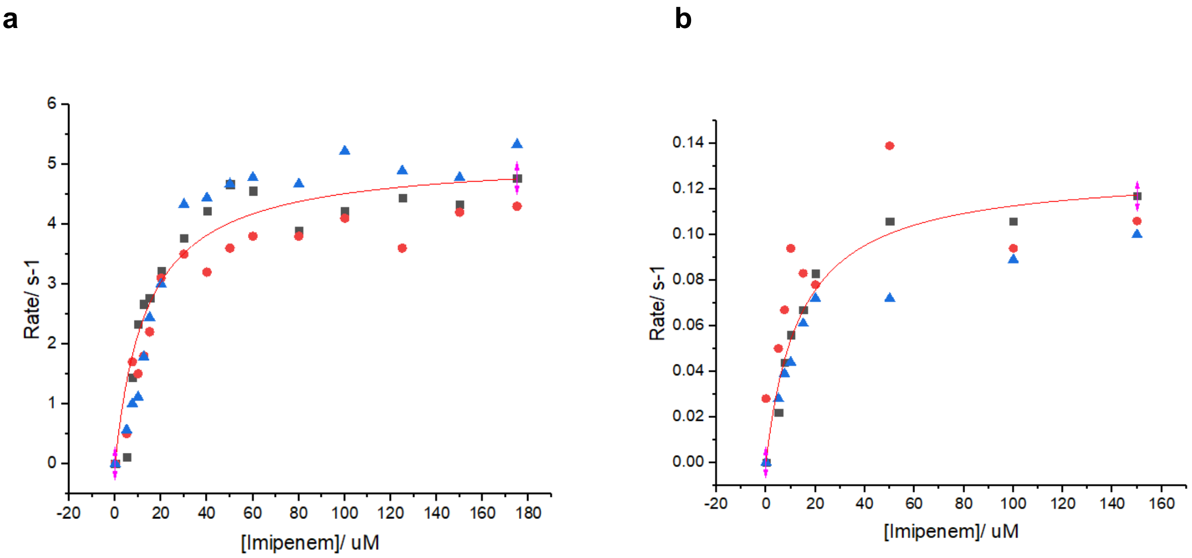

**Supplementary Figure 2** Michaelis-Menten kinetics were measured in triplicate for OXA-$48_{AcK73}$ variant (**b**) to compare with OXA-$48_{WT}$ (**a**).

For OXA-$48_{WT}$, $K_M$ = 13.79 μM, $k_{cat}$ = 5.13 $s^{-1}$, and $k_{cat}/K_M$ = 3.72 x $10^5$ $M^{-1}s^{-1}$. For OXA-$48_{AcK73}$, $K_M$ = 13.89 μM and $k_{cat}$ = 0.13 $s^{-1}$, and $k_{cat}/K_M$ = 9.36 x $10^3$ $M^{-1}s^{-1}$.

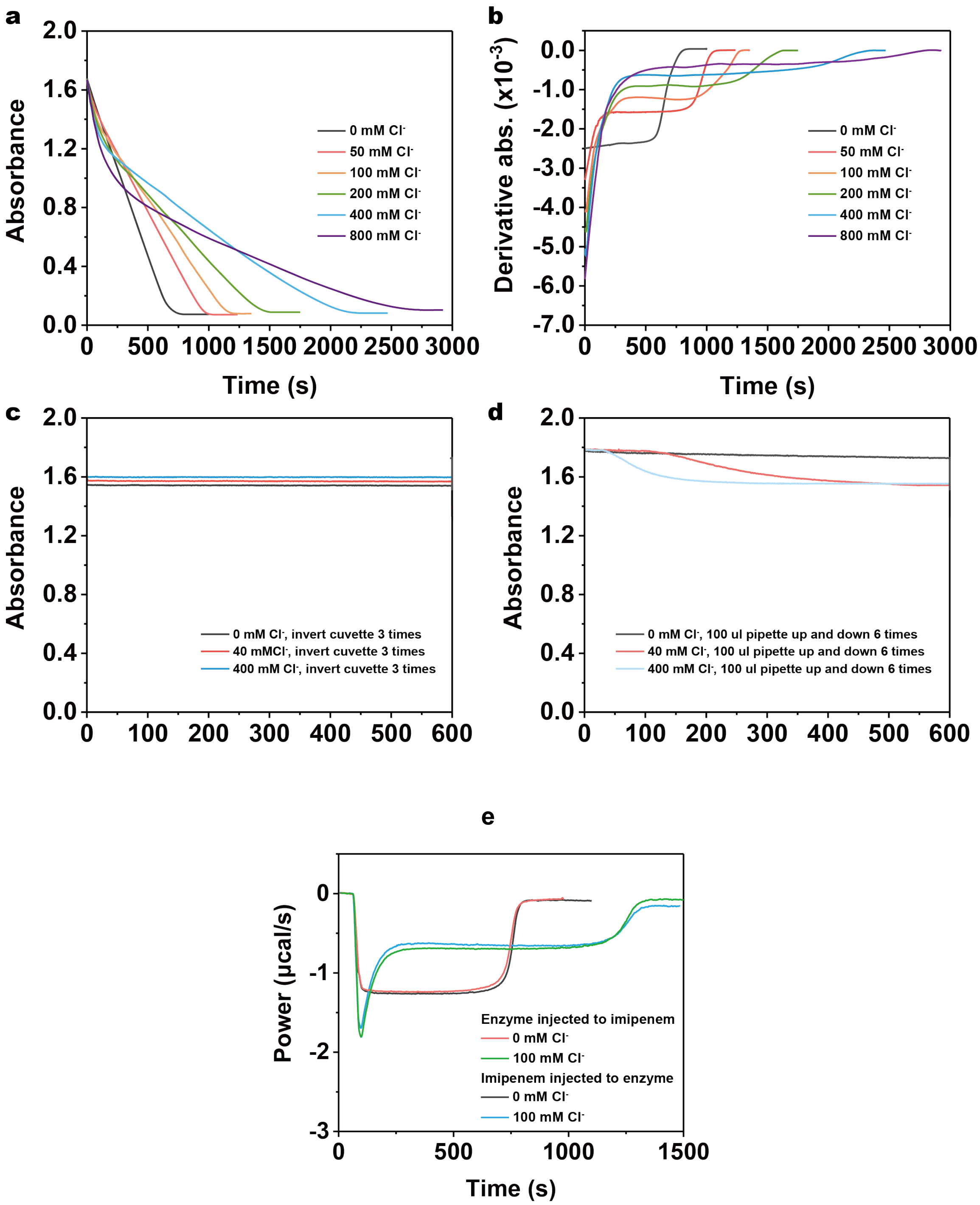

**Supplementary Figure 3** UV-Vis for monitoring imipenem hydrolyzed by OXA-48 to show it is compatible with ITC, and comparison of the ITC heat profile to show the fast-mixing deadtime is negligible when the order of injection was changed.

(**a**) UV-Vis spectra of OXA-48 hydrolyzing imipenem in the buffer with a series of chloride ion concentrations. Final concentration: 200 µM imipenem, 100 nM OXA-48 in 50 mM NaPi, pH 7.5, 1 mM $NaHCO_3$ with 0-800 mM chloride. (**b**) The first derivative d*A*/d*t* of the absorbance in (a) shows the rate of change of absorbance, which is compatible with the rate of change of heat measured by ITC. (**c**, **d**) Comparison of mixing methods of “inverting cuvette” with “pipetting” for the UV-Vis assay in the presence of imipenem and chloride, to show the UV-Vis absorbance spectra is highly sensitive to different mixing techniques. The absorbance of imipenem was monitored at 300 nm. Final concentration: 200 μM imipenem in 50 mM NaPi, pH 7.5 buffer supplemented with different chloride concentrations. All assays were carried out in a total volume of 1 mL with 100 nM OXA-48, 200 μM imipenem and 0-800 mM chloride. The decrease of imipenem absorbance was monitored continuously at 300 nm. Data were plotted using Origin 2017 Software. (**e**) Comparison of the order of injection in ITC assays is not affecting reproducibility. 200 μM imipenem solution and 100 nM OXA-$48_{WT}$ solution are fast mixed when one is injected into the other in ITC in the matched buffer of 50 mM NaPi, pH 7.5, 1 mM $NaHCO_3$ buffer with 0 or 100 mM chloride.

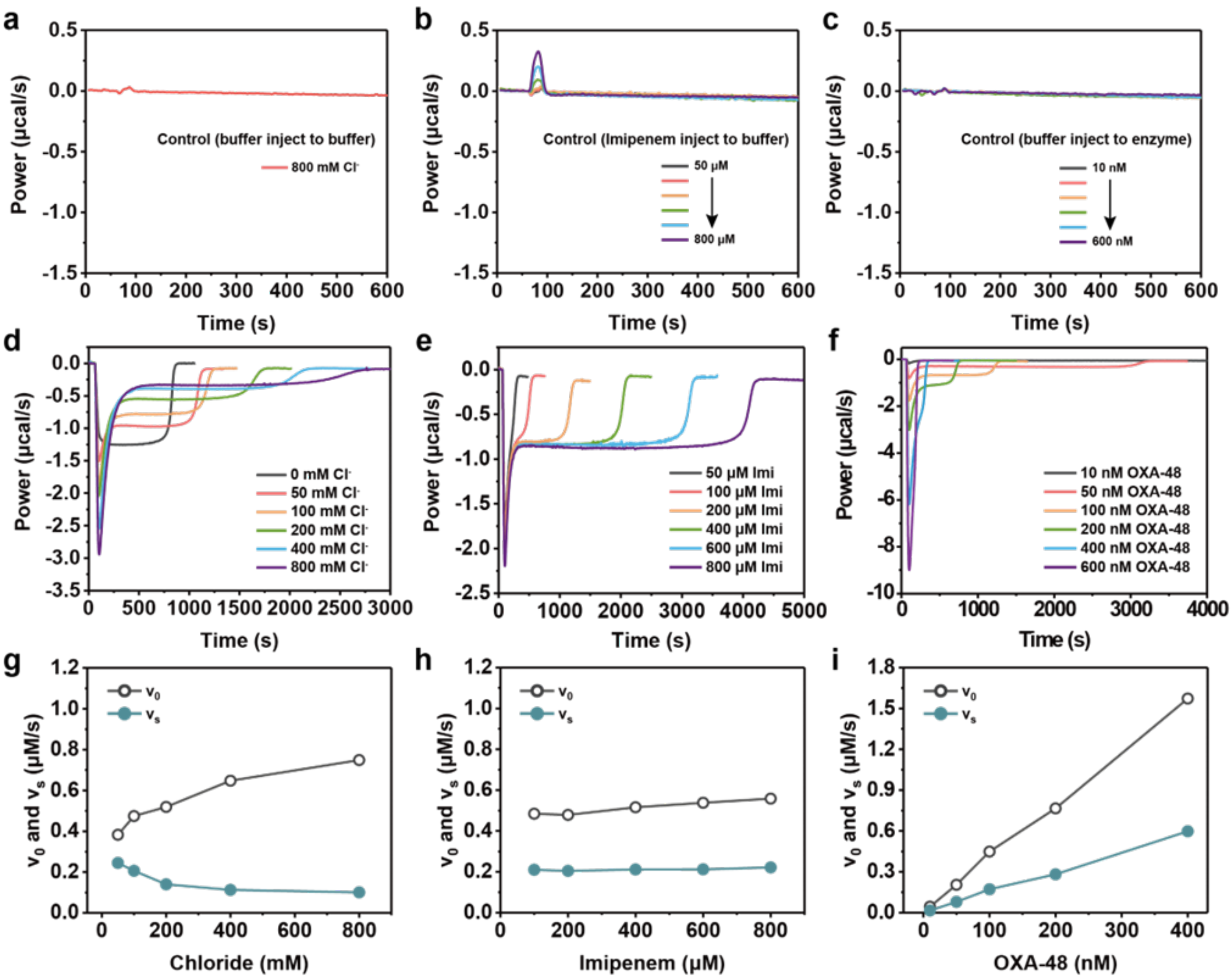

**Supplementary Figure 4** Effect of enzyme, substrate or chloride concentrations on biphasic kinetics in ITC assays.

(**a**) Buffer (50 mM NaPi, 800 mM NaCl, 1 mM $NaHCO_3$, pH 7.5) was injected into the same buffer as control; (**b**) Different concentrations of imipenem in buffer 50 mM NaPi, pH 7.5, 100 mM NaCl, 1 mM $NaHCO_3$ was injected into the same buffer as control; (**c**) Buffer (50 mM NaPi, 100 mM NaCl, 1 mM $NaHCO_3$, pH 7.5) was injected into different concentrations of OXA-48 in the same buffer as control; (**d**) Effect of chloride concentrations on biphasic kinetics. 200 μM substrate was injected into 100 nM OXA-$48_{WT}$ in the matched buffer (0-800 mM NaCl, 50 mM NaPi, pH 7.5, 1 mM $NaHCO_3$); (**e**) Effect of substrate concentrations on biphasic kinetics. 50-800 μM imipenem in the buffer of 50 mM NaPi, pH 7.5, 1 mM $NaHCO_3$, and 100 mM NaCl was injected into 100 nM enzyme in the matched buffer; (**f**) Effect of OXA-$48_{WT}$ concentrations on biphasic kinetics. 10 - 600 nM OXA-$48_{WT}$ in the buffer of 50 mM NaPi, pH 7.5, 1 mM $NaHCO_3$, and 100 mM NaCl was injected into 200 μM substrate in the matched buffer. Based on the ITC curves in (**d-f**), $v_0$ and $v_s$ for (**g**) different [$Cl^-$], (**h**) [substrate] and (**i**) [OXA-$48_{WT}$] were plotted to show $v_0/v_s$ is only [$Cl^-$]-dependent.

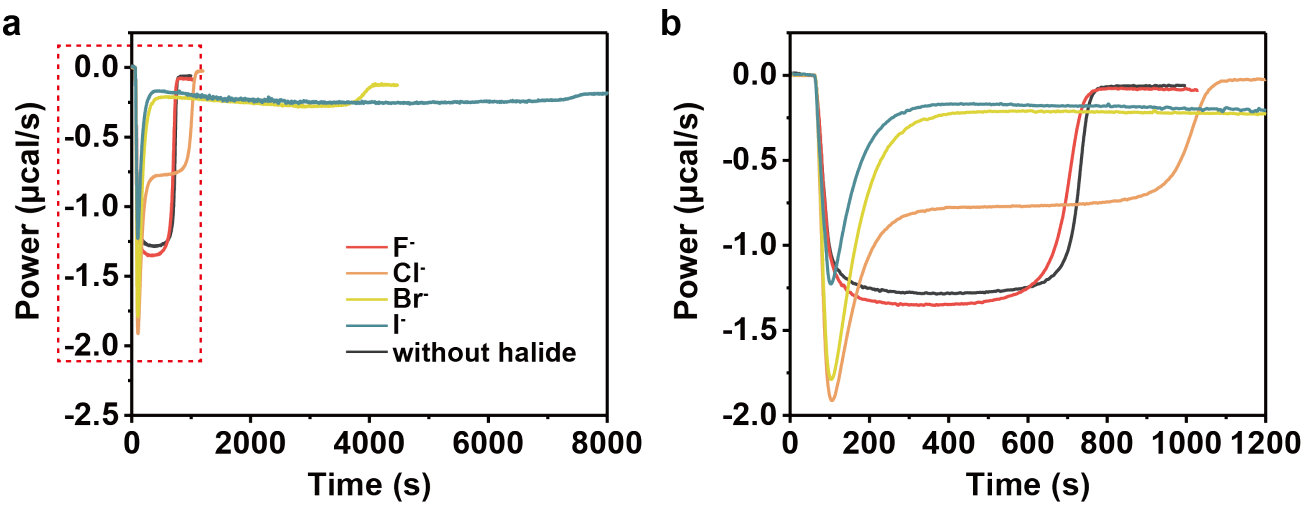

**Supplementary Figure 5** Effect of four halide ions at 100 mM on hydrolysis of imipenem by OXA-48. (**a**) 10 μL imipenem solution was injected into 210 μL of OXA-$48_{WT}$ (final concentration: 200 μM imipenem, 100 nM OXA-$48_{WT}$), both prepared in buffer of 50 mM NaPi, pH 7.5, 1 mM $NaHCO_3$, with 100 mM corresponding halide ions). (**b**) is a closeup of (a) from 0 to 1200 s.

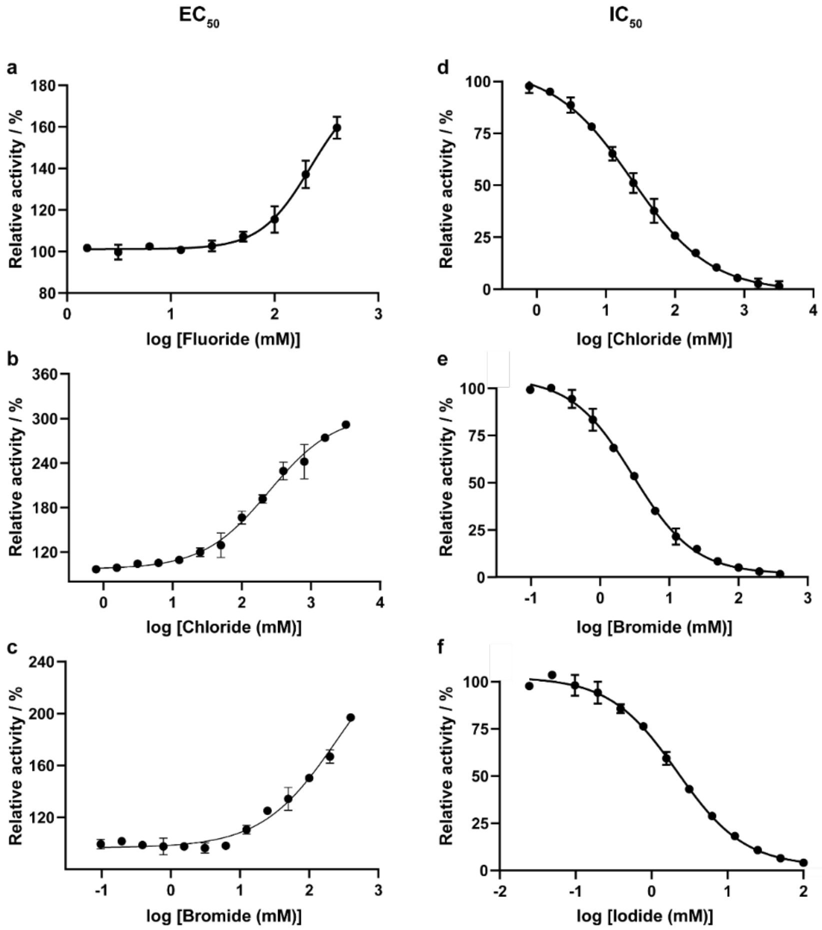

**Supplementary Figure 6** $EC_{50}$ plots of (**a**) fluoride, (**b**) chloride and (**c**) bromide to reflect the accelerating effect during imipenem hydrolysis by OXA-$48_{WT}$. $IC_{50}$ plots of (**d**) chloride, (**e**) bromide and (**f**) iodide to reflect the inhibiting effect during imipenem hydrolysis by OXA-48.

All 1 mL reactions were set up using 200 μM imipenem as the substrate in the buffer of 50 mM NaPi, pH 7.5, with various concentrations of halide ions by adding NaI, NaBr, NaCl, or NaF. Reactions were initiated by adding OXA-48$_{WT}$ to a final concentration of 100 nM and the absorbance of imipenem at 300 nm were recorded continuously. All curves are fitted with nonlinear regression analysis.

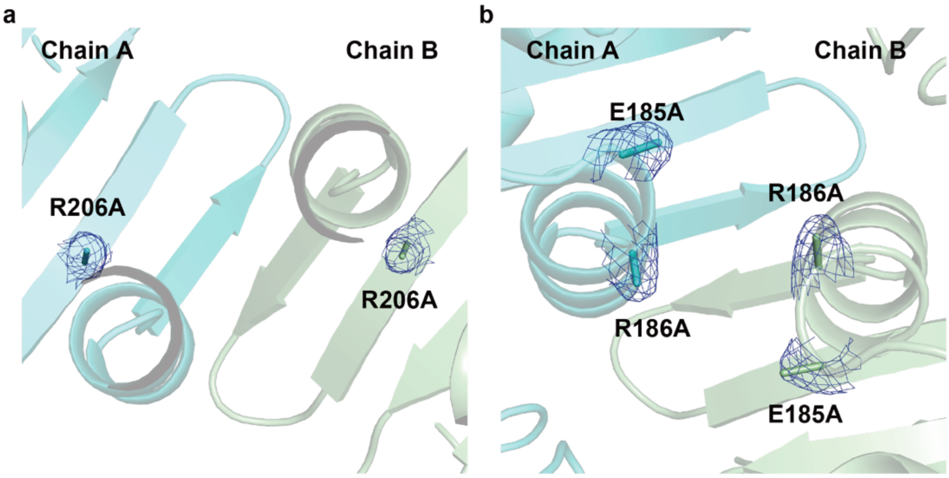

**Supplementary Figure 7** The crystal structure (7PEI) of OXA-48$_{E185A/R186A/R206A}$ with interface chloride binding site mutations to confirm the abolished bromide binding after mutating these residues to alanine.

(**a**) The interface of R206A-R206A' between two OXA-48$_{E185A/R186A/R206A}$ molecules. (**b**) The interface of E185A-E186A-E185A'-E186A' between two OXA-48$_{E185A/R186A/R206A}$ molecules. 2$F_o$−$F_c$ electron density map of mutated residual is shown as blue mesh, contoured at 1σ (0.2216 e$^-$/Å$^3$).

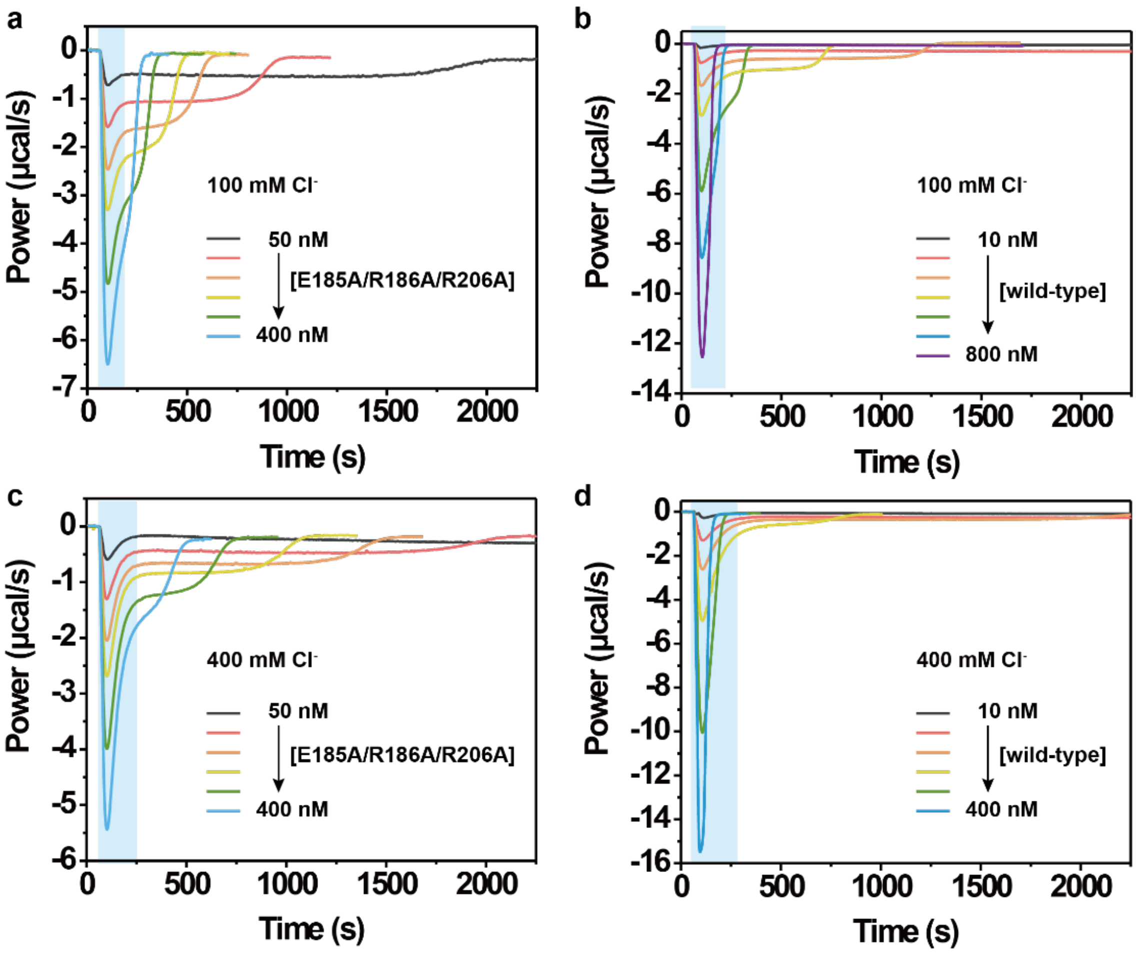

**Supplementary Figure 8** Comparing the OXA-48$_{WT}$ and OXA-48$_{E185A/R186A/R206A}$ for the burst phase for the level of allosteric effect.

Imipenem was hydrolyzed by various concentrations of interface variant OXA-48$_{E185A/R186A/R206A}$ and OXA-48$_{WT}$ in the presence of (**a, b**) 100 mM NaCl and (**c, d**) 400 mM NaCl. Experiments were performed by injecting 10 µL imipenem solution (final concentration 200 µM) into a sample cell filled with 208.7 µL enzyme (final concentration 50-400 nM for OXA-48$_{E185A/R186A/R206A}$, 10-800 nM for OXA-48$_{WT}$), both in the buffer of 50 mM NaPi, pH 7.5, 1 mM $NaHCO_3$, and 100 mM or 400 mM NaCl.

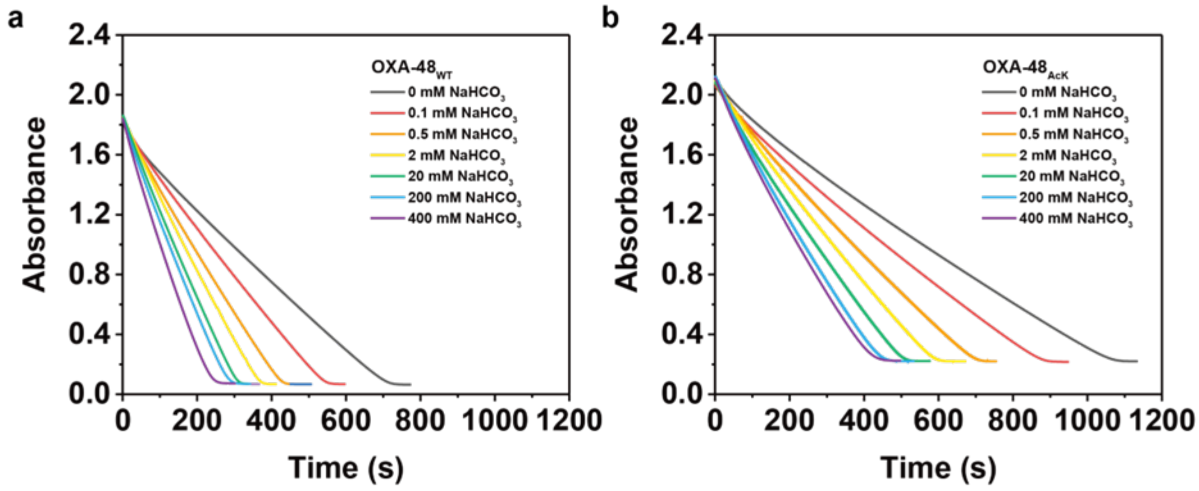

**Supplementary Figure 9** $HCO_3^-$ is a weak allosteric effector of OXA-48.

**(a, b)** UV-Vis spectra of 200 μM imipenem hydrolysis by (**a**) 100 nM OXA-$48_{WT}$ and (**b**) 8 μM OXA-$48_{AcK73}$ in 50 mM NaPi buffer, pH 7.5 supplemented with 0-400 mM $HCO_3^-$.

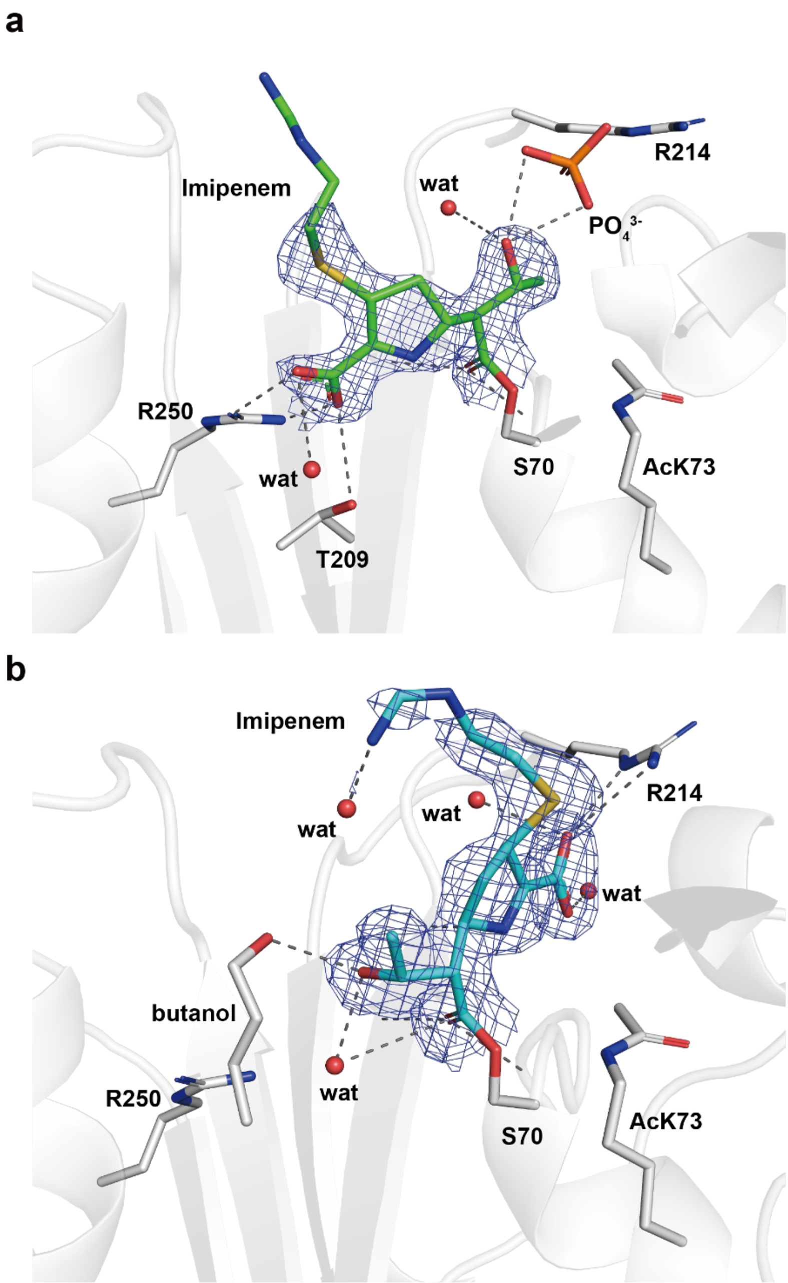

**Supplementary Figure 10** The crystal structure of chloride-free OXA-48AcK-imipenem acyl-intermediate complex.

(**a**) The active conformation and (**b**) the inactive conformation. Unbiased $2F_o - F_c$ electron density map of imipenem is shown as blue mesh, contoured at 1σ (0.3116 $e^-/Å^3$).

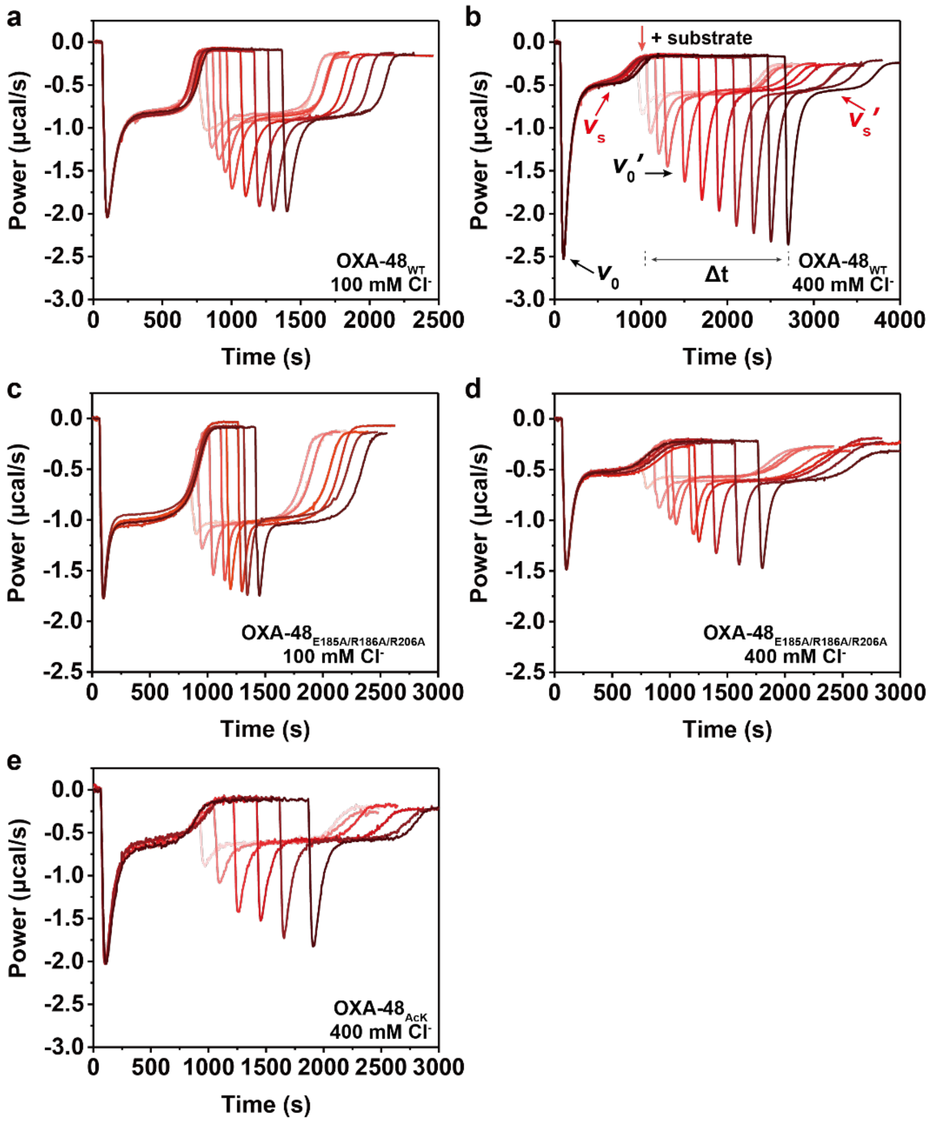

**Supplementary Figure 11** Overlays of kinetics curves for titrating different time intervals between two imipenem injections into OXA-48 in order to show the time required for the initial activity recovery before the 2nd injection.

(**a**) 180 µM imipenem x2 was hydrolyzed by 100 nM OXA-$48_{WT}$ in the buffer of 50 mM NaPi, pH 7.5, 1 mM $NaHCO_3$, supplemented with 100 mM $Cl^-$. (**b**) 140 µM imipenem x2 was hydrolyzed by 100 nM OXA-$48_{WT}$ in the buffer of 50 mM NaPi, pH 7.5, 1 mM $NaHCO_3$, supplemented with 400 mM $Cl^-$. (**c**) 200 µM imipenem x2 was hydrolyzed by 100 nM interface variant OXA-$48_{E185A/R186A/R206A}$ in the buffer of 50 mM NaPi, pH 7.5, 1 mM $NaHCO_3$, supplemented with 100 mM $Cl^-$. (**d**) 80 µM imipenem x2 was hydrolyzed by 100 nM interface variant OXA-$48_{E185A/R186A/R206A}$ in the buffer of 50 mM NaPi, pH 7.5, 1 mM $NaHCO_3$, supplemented with 400 mM $Cl^-$. (**e**) 150 µM imipenem x2 was hydrolyzed by 6.5 µM OXA-$48_{AcK}$ in the buffer of 50 mM NaPi, pH 7.5, containing 1 mM $NaHCO_3$, supplemented with 400 mM $Cl^-$. All concentrations are the final concentrations used in the reaction.

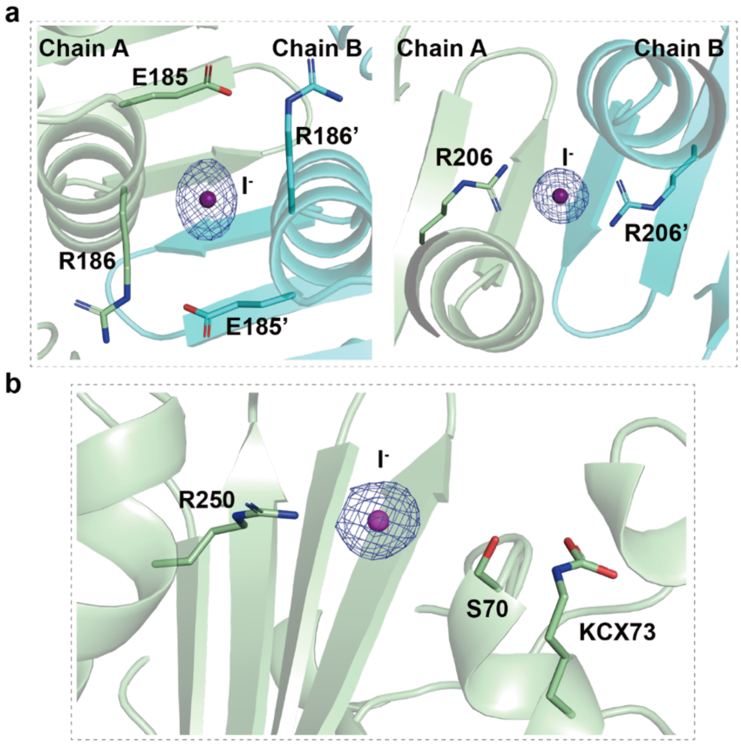

**Supplementary Figure 12** Crystal structure of OXA-48$_{WT}$ with iodide ions.

(**a**) Binding sites for iodide ions at the OXA-48$_{WT}$ dimer interface. (**b**) Binding sites for iodide ions in the OXA-48$_{WT}$ active site. Unbiased 2$F_o$−$F_c$ electron density map of iodide is shown as blue mesh, contoured at 1σ (0.3053 $e^-/A^3$).

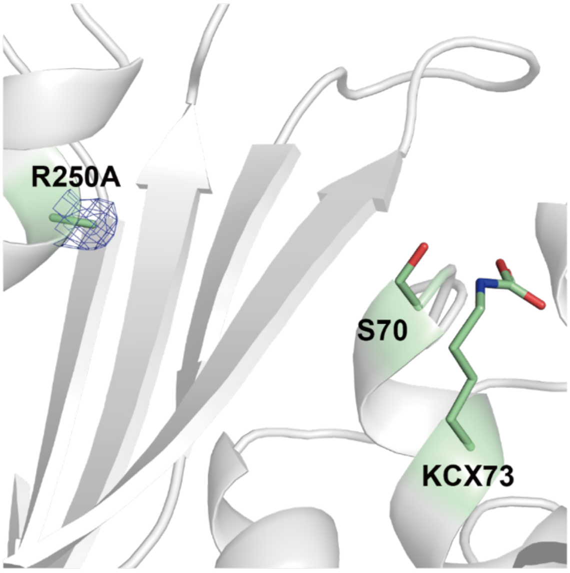

**Supplementary Figure 13** Crystal structure (7PGO) of OXA-48$_{R250A}$ soaked with $Br^-$, but no $Br^-$ density was found near R250A.

Unbiased 2$F_o$−$F_c$ electron density map of the mutated residual is shown as blue mesh, contoured at 1σ (0.3555 $e^-/Å^3$).

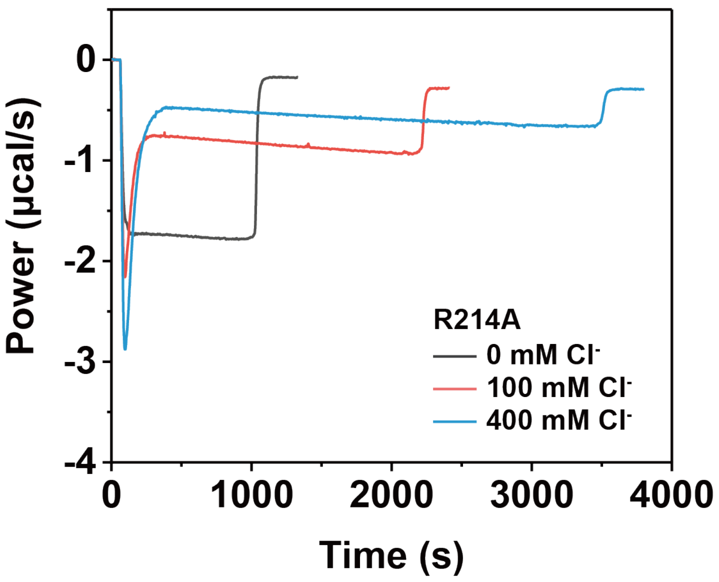

**Supplementary Figure 14** ITC kinetic curves of 400 µM imipenem hydrolysis by 3 µM OXA-48$_{R214A}$ variant with 0 mM, 100 mM or 400 mM chloride ion in the buffer of 50 mM NaPi, pH 7.5, and 1 mM $NaHCO_3$.

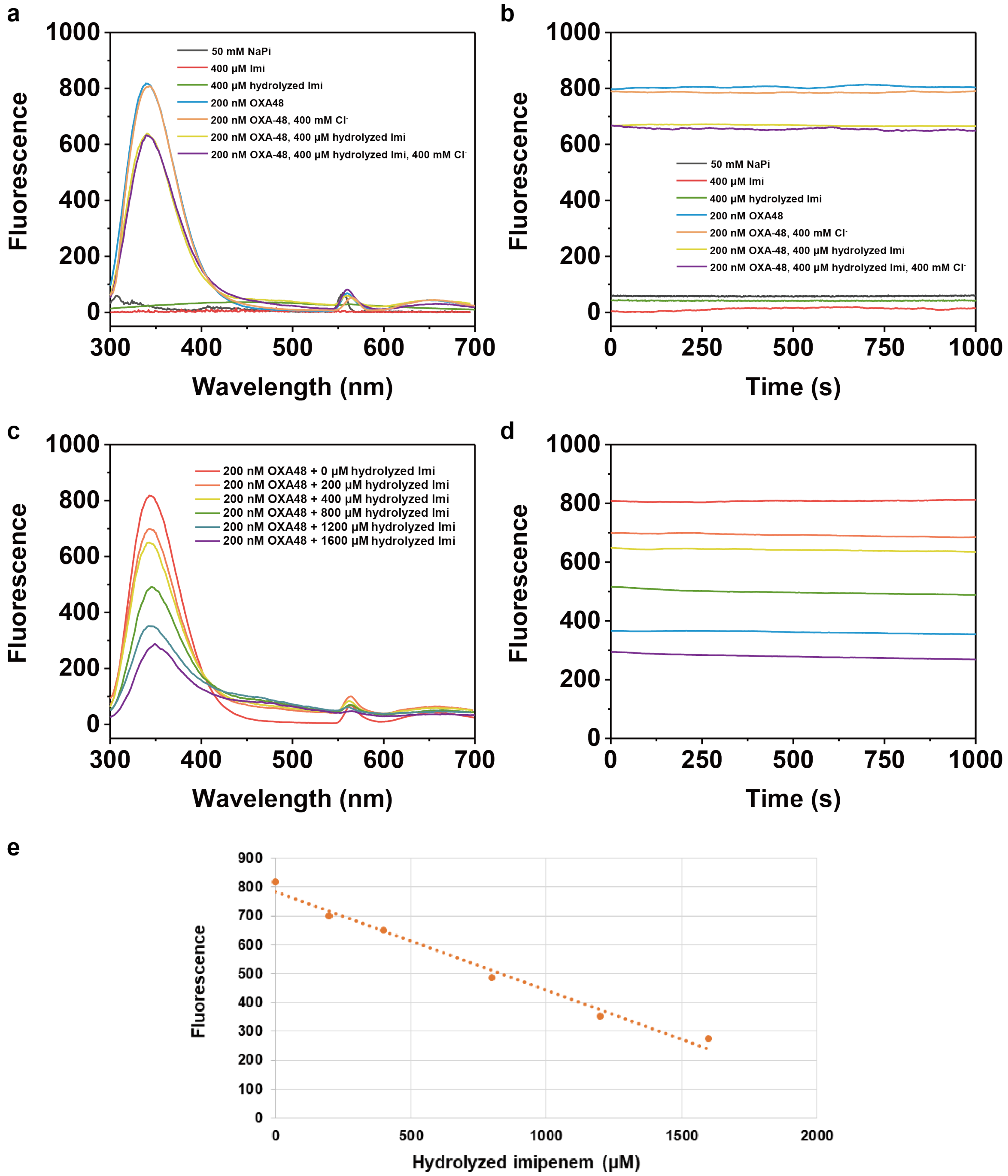

**Supplementary Figure 15** The intrinsic tryptophan fluorescence spectra of OXA-48$_{WT}$ to show the intensity at $\lambda_{em}$ = 340 nm is only affected by the concentration of hydrolyzed imipenem product, but not by other components in the assay.

(**a, b**) The intrinsic tryptophan fluorescence spectra ($\lambda_{ex}$ = 280 nm, $\lambda_{em}$ = 300-700 nm) of 200 nM OXA-48$_{WT}$ with the addition of 400 μM hydrolyzed imipenem product and other components in the buffer to demonstrate the fluorescence of the OXA-48 is not affected by other components in the assay but only the product of imipenem. Fluorescence is stable within the time frame of the activity assay. (**c, d**) The intrinsic fluorescence spectra of 200 nM OXA-48$_{WT}$ mixed with 0-1600 μM hydrolyzed imipenem at $\lambda_{ex}$ = 280 nm, $\lambda_{em}$ = 300-700 nm, showing the product quantitatively reduces the intrinsic fluorescence intensity. (**e**) Linear fitting plot of fluorescence from 200 nM OXA-48$_{WT}$ protein versus 0-

1600 μM hydrolyzed imipenem. All the reactions were performed in 2 mL buffer of 50 mM NaPi, pH 7.5, 1 mM $NaHCO_3$, with 0 or 400 mM chloride at 25 °C.

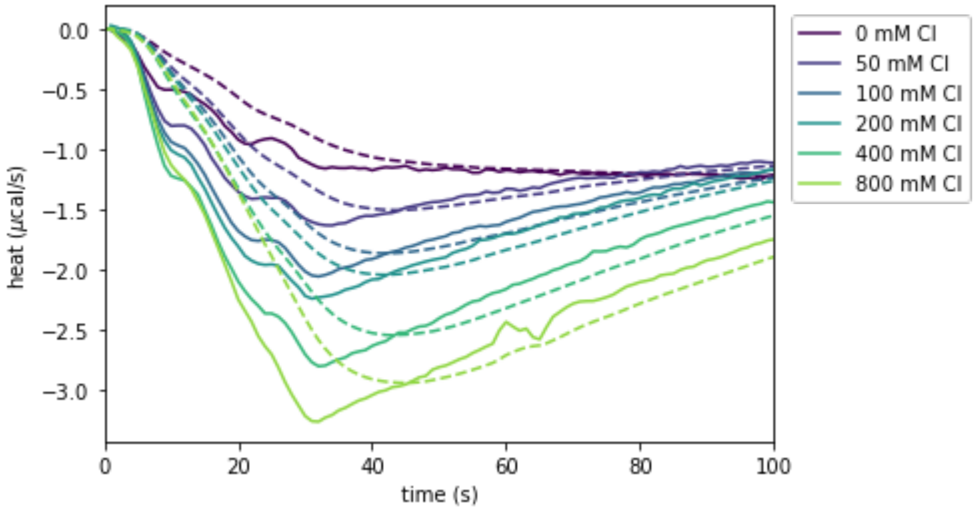

**Supplementary Figure 16** Converting between measured and real heat signal. Solid lines represent experimental data, while dashed lines are simulations of the model for 0 – 800 mM NaCl.

The measured signal in the experiment (dashed lines) and the real signal occurring at a given moment (solid lines) can be interconverted using Eq. 2. The plot shows a close-up view of the first seconds of the experiment, where the burst phase occurs. Note that the measured signal is much smoother than the real signal.

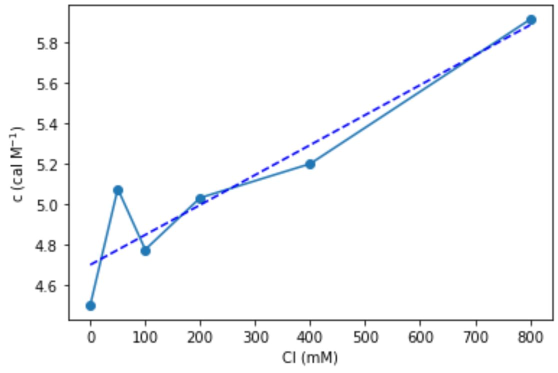

**Supplementary Figure 17** Numerically obtaining $c$, the conversion between moles of substrate and calories of heat generated.

This value was obtained for different chloride concentrations, obtaining a linear relationship between chloride concentration and *c*, as follows: $c = 4.7 + 0.0015 * [Cl]$, with an $R^2$ = 0.871.

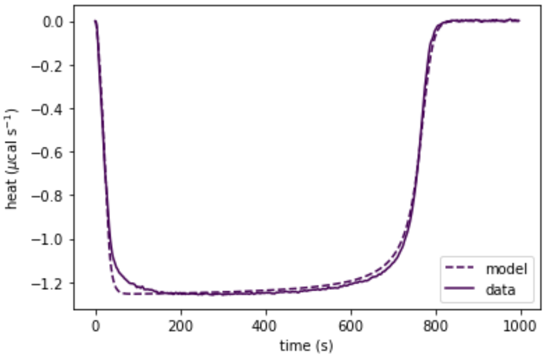

**Supplementary Figure 18** A numerical fit to the data confirms that $k_1$' and $k_2$' are of the same order of magnitude.

**a** **b** **c**

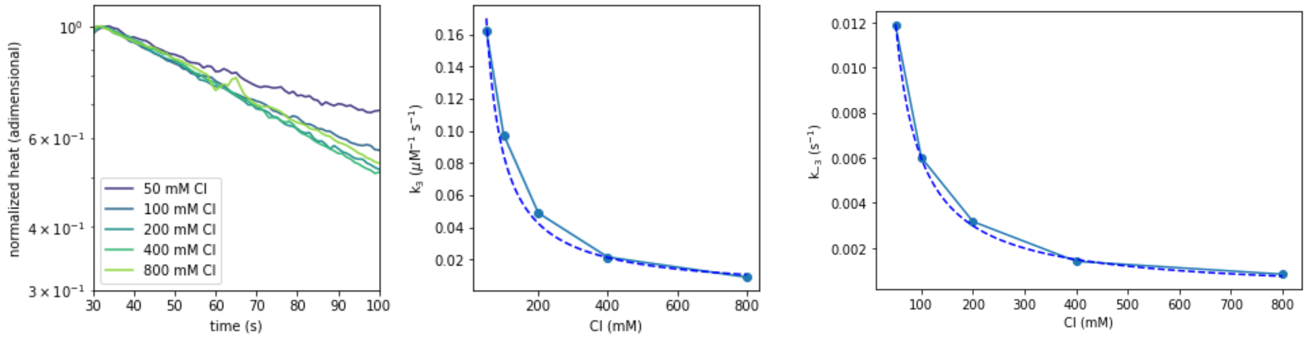

**Supplementary Figure 19** $k_3$' and $k_{-3}$' decrease with chloride.

(**a**) Dividing heat by its maximum value and representing the result in semilog axes, it is clear that the heat is decreasing exponentially in all chloride concentrations. (**b**) Fitting these data points to a line, we obtain almost constant values for the slope $k_3$'[Cl⁻] (values ranging between 0.0075 and 0.01 $s^{-1}$), which results in an inverse relationship between $k_3$' and chloride (solid circles, solid line added as a visual guide) which is well captured by the inverse curve 8.50/[Cl] (dashed line), with $R^2$ = 0.983. (**c**) Using the previous formula to obtain $k_{-3}$' (solid circles, solid line added as a visual guide), we observe an inverse relationship with chloride that can be captured with the equation 0.60/[Cl] (dashed line), with $R^2$ = 0.999. This relationship implies that $k_3$'/$k_{-3}$' ≈ 14.27 $\mu M^{-1}$.

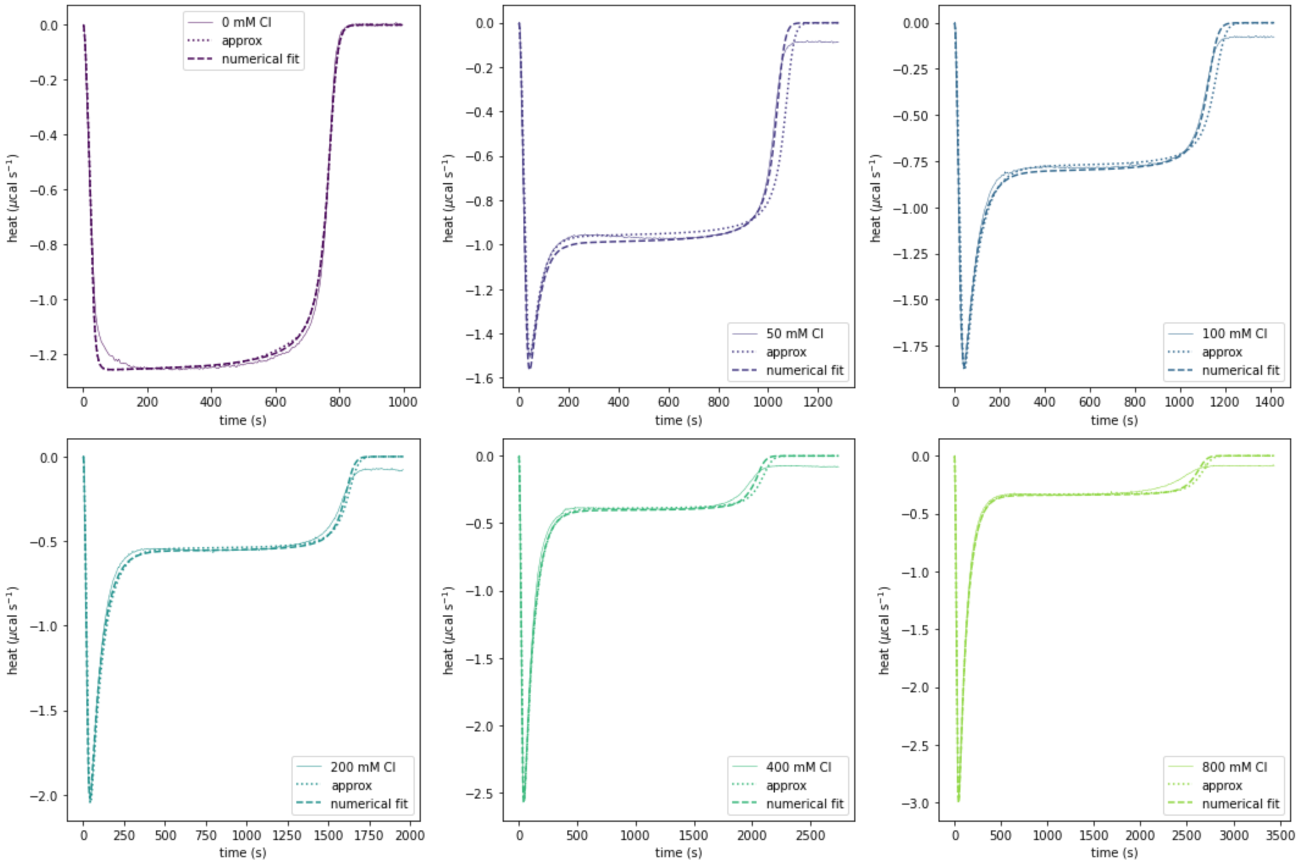

**Supplementary Figure 20** The analytical approximations are good fits to the data at different chloride concentrations. Solid lines represent experimental data, while dashed lines are simulations of the model.

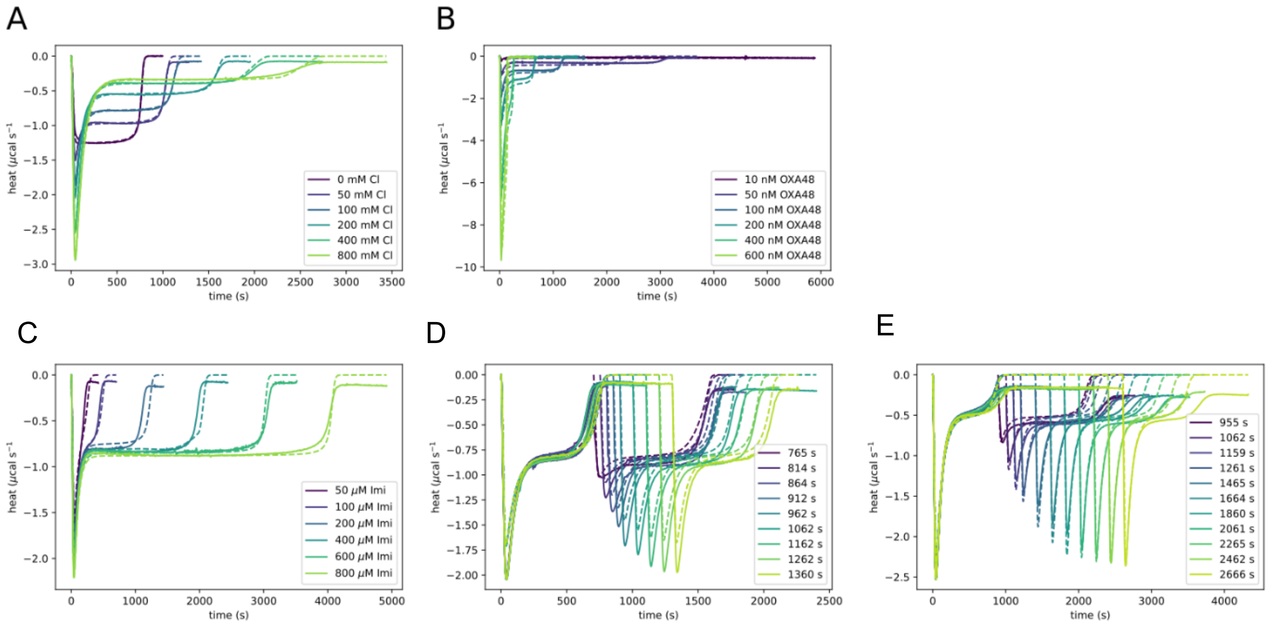

**Supplementary Figure 21** Our mathematical model accurately fits the experimental ITC data including single-injection and double-injection assays. Solid lines represent experimental data, while dashed lines are simulations of the model.

ITC curves were obtained with (**A**) different chloride concentrations, with initial enzyme at 100 nM and initial substrate at 200 μM, (**B**) different enzyme concentrations, with initial enzyme at 100 nM and chloride at 100 mM, (**C**) different substrate concentrations, with initial enzyme at 100 nM and chloride at 100 mM. Finally, the last two panels show the reactivation experiments, where new substrate is injected at different time points. In both cases there is enzyme at 100 nM, with (**D**) chloride at 100 mM and substrate at 180 μM and (**E**) chloride at 400 mM and substrate at 140 μM. In the double-titration experiments, we introduced a progressive delay for the activation of $k_3$ which helps to capture the peaks. For (D) we used a delay of $37(t_f - 705)/(t_f - 600)$, where $t_f$ is the reinjection time, while for (E) we used $37(t_f - 900)/(t_f - 700)$.

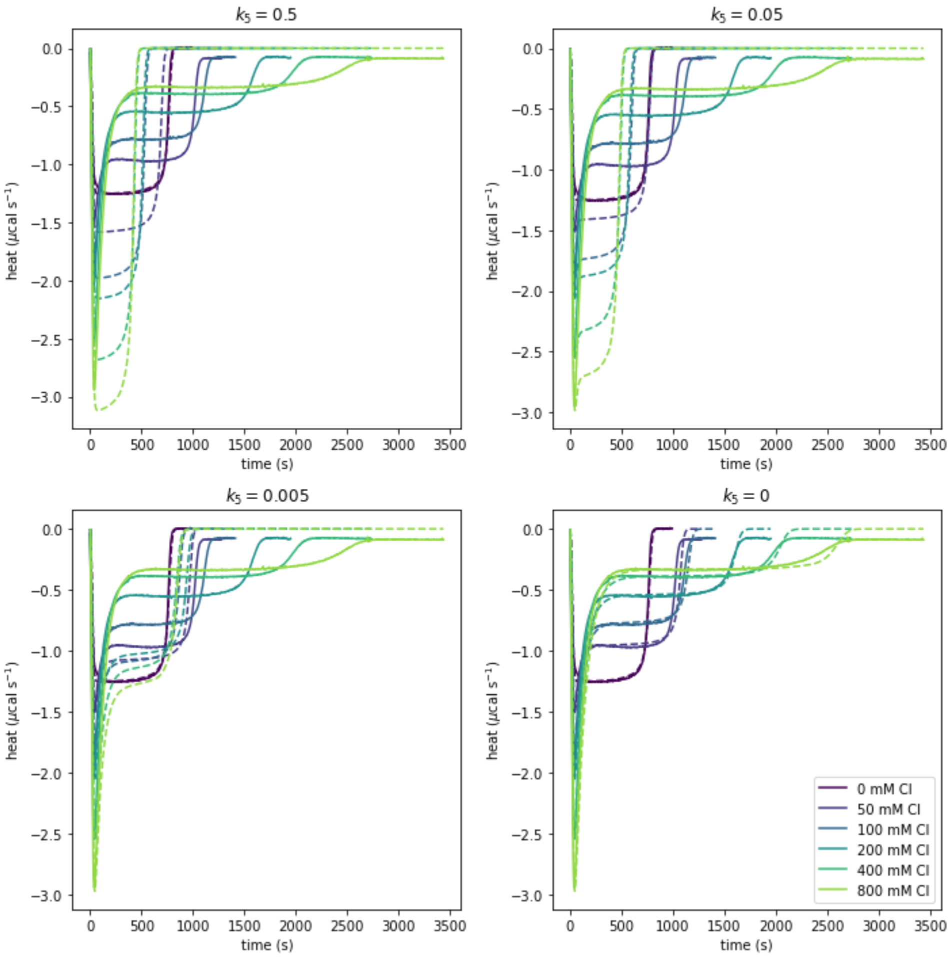

**Supplementary Figure 22** Introducing $k_5$ fails to capture the ITC data even for very low values from 0 to 0.5 s$^{-1}$, suggesting that there is no hydrolysis from the inactive intermediate leading to product.

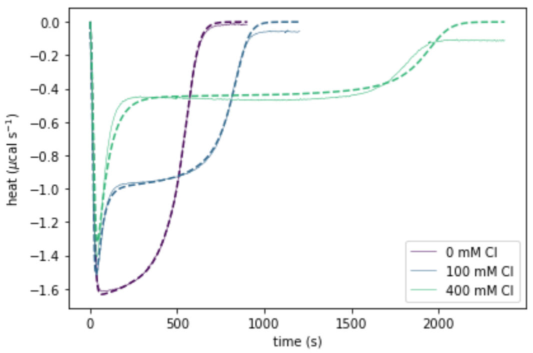

**Supplementary Figure 23** Simulated and experimental apparent reaction rates for the interface variant OXA-$48_{E185A/R186A/R206A}$.

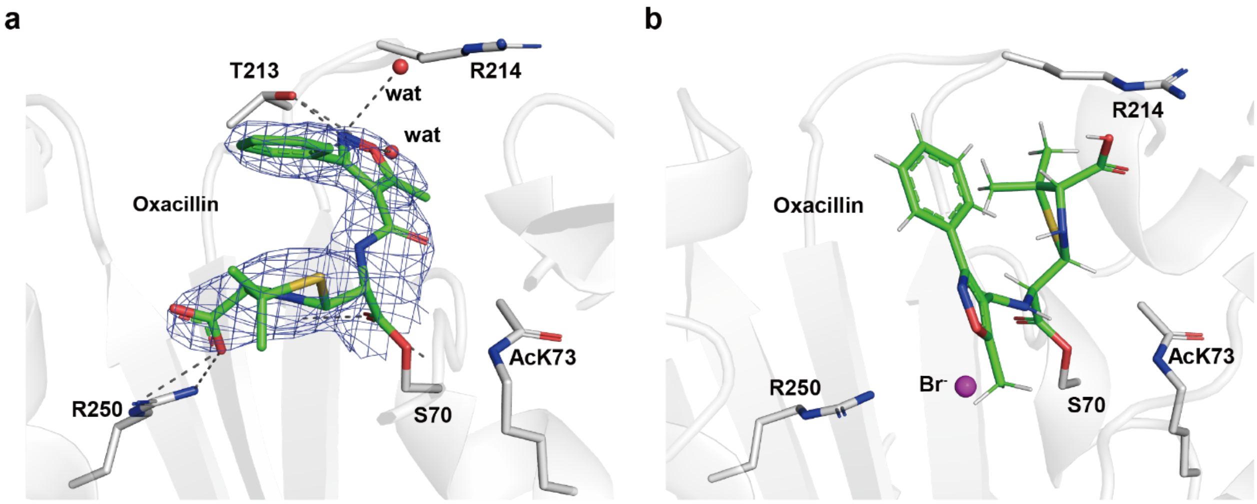

**Supplementary Figure 24 (a)** The crystal structures of OXA-$48_{AcK73}$-oxacillin acyl-intermediate complex captured in a crystallization condition free of chloride, and (**b**) the docked OXA-$48_{AcK73}$-oxacillin acyl-intermediate with bromide.

The initial unbiased $2F_o{-}F_c$ electron density map of oxacillin in (a) is shown as blue mesh, contoured at 1σ (0.2087 $e^-/Å^3$). The structure of oxacillin acyl-intermediate (in green) with bromide (in purple) binding at OXA-$48_{AcK}$ active site (in grey) in (b) was generated by molecular docking using MOE[1]. The docking was generated from the structure with OXA-$48_{AcK}$-imipenem-$Br^-$ complex H chain. Protein hydrogens are hidden for clarity.

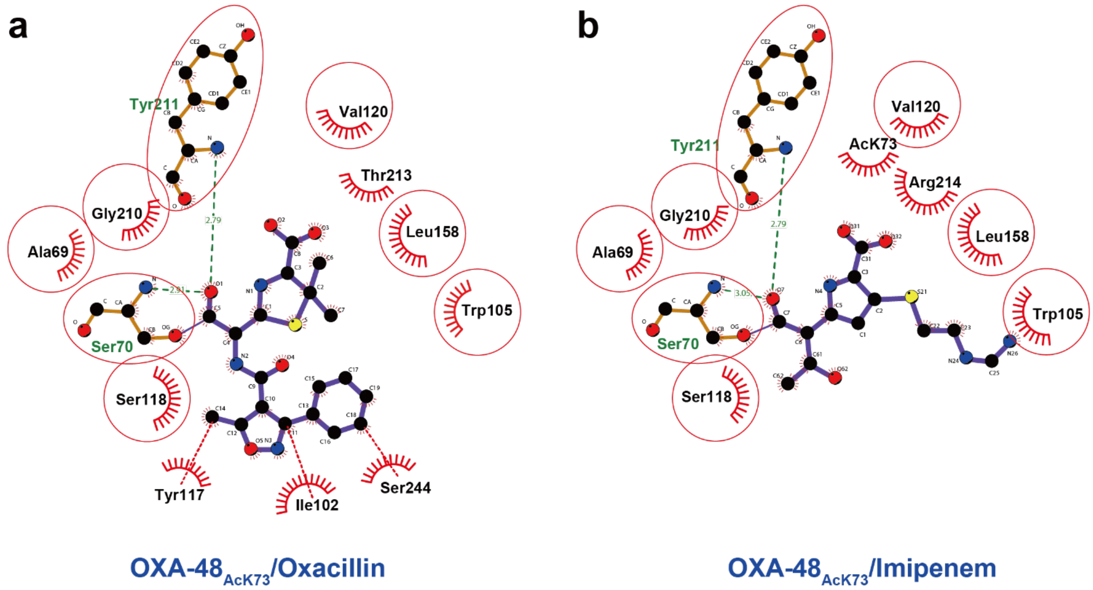

**Supplementary Figure 25** Ligplot diagrams show the hydrogen bonding and hydrophobic interactions between OXA-48$_{AcK73}$ with the inactive acyl-intermediate conformations of (**a**) oxacillin and (**b**) imipenem.

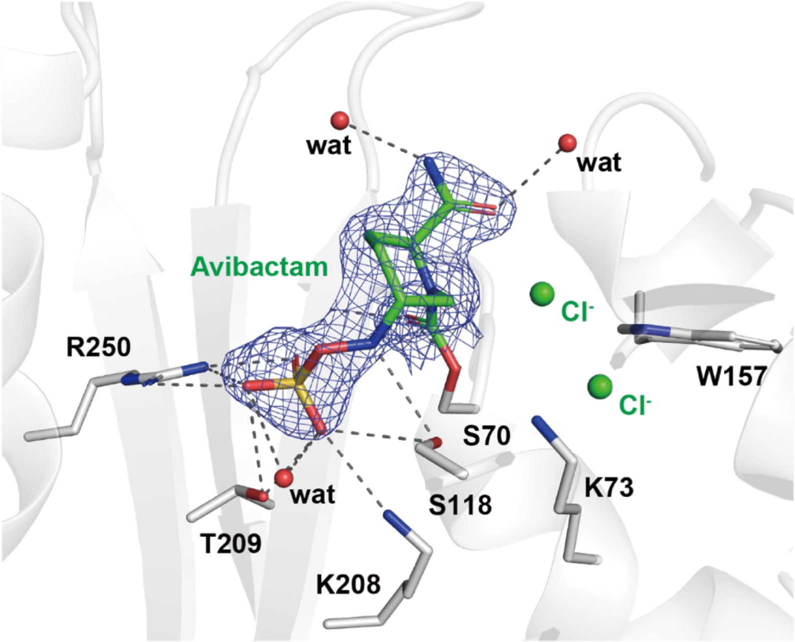

**Supplementary Figure 26** Crystal structure of OXA-48$_{WT}$-avibactam-Cl$^{-}$ complex.

The initial unbiased 2$F_o$−$F_c$ electron density map of avibactam is shown as blue mesh, contoured at 3σ (0.2687 e$^{-}$/Å$^{3}$).

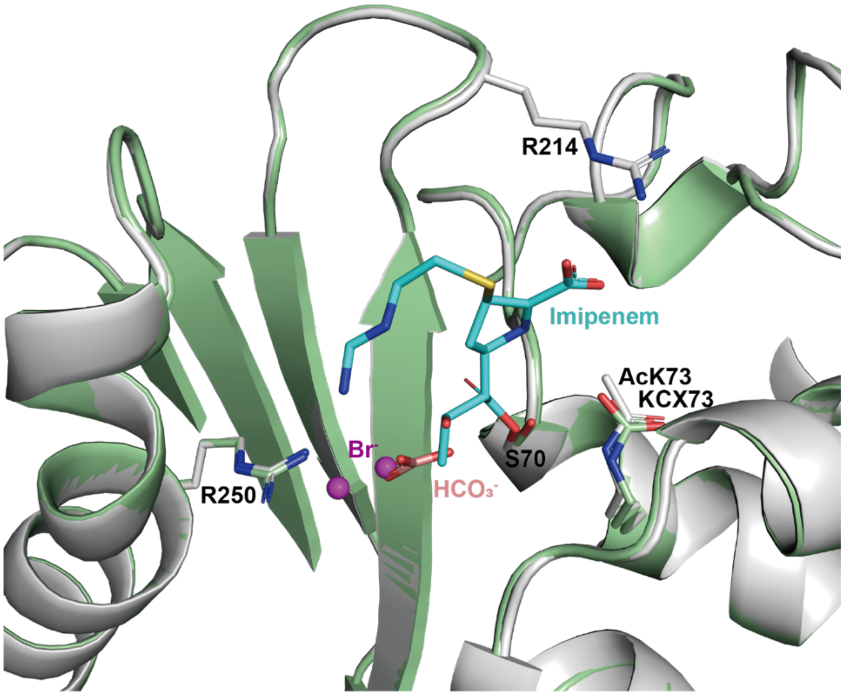

**Supplementary Figure 27** Superimposition of the OXA-48$_{AcK73}$-imipenem-Br$^{-}$ acyl-intermediate complex in the inactive conformation with the OXA-48$_{WT}$-$HCO_3^{-}$ complex.

OXA-48$_{AcK73}$-imipenem-Br$^{-}$ acyl-intermediate: protein in grey, imipenem in cyan, and bromide in purple. OXA-48$_{WT}$-$HCO_3^{-}$ complex: protein in green and $HCO_3^{-}$ in salmon.

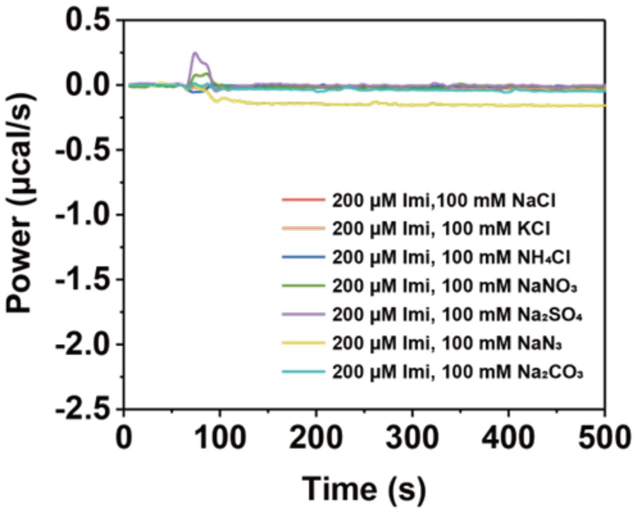

**Supplementary Figure 28** The control ITC heat profiles show there is only negligible heat detected when there is no OXA-48$_{WT}$.

Imipenem solution in the buffer of 50 mM NaPi, pH 7.5, 1 mM $NaHCO_3$ with various salts was injected into the sample cell filled with matched buffer.

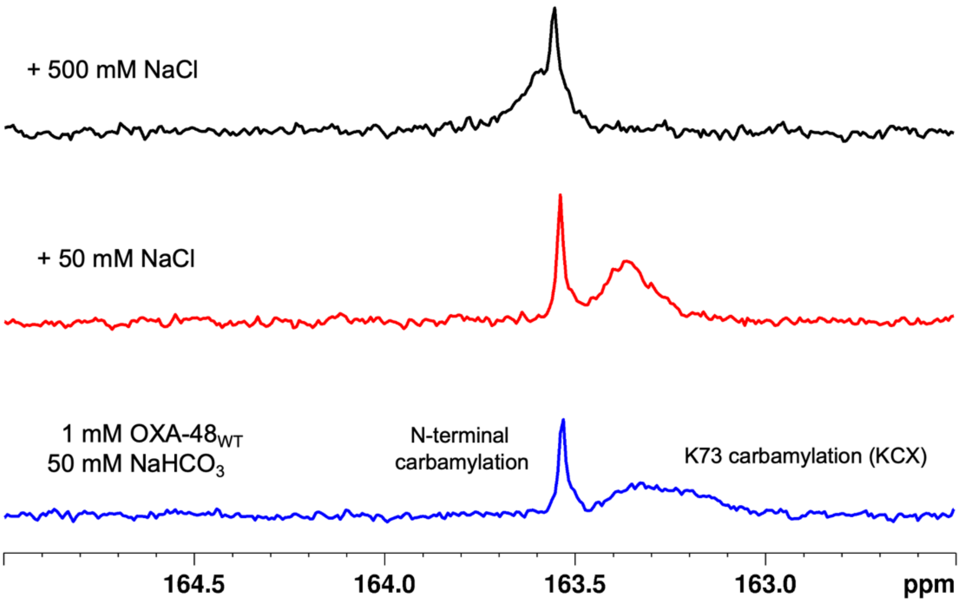

**Supplementary Figure 29** $^{13}C$ NMR spectra of carbamylated OXA-$48_{WT}$ and chemical shift changes caused by $Cl^{-}$.

A final concentration of 0, 50 mM and 500 mM NaCl was titrated into the sample of 1 mM OXA-$48_{WT}$ in 100 mM NaPi, pH 7.5, 50 mM $NaH^{13}CO_3$, 10% $D_2O$ buffer. It showed even 50 mM of $Cl^{-}$ can change the chemical shift of the KCX resonance. Please note, the sharp signal at 164 ppm is from N-terminal carbamylation and can only be observed at 50 mM not at 10 mM $NaH^{13}CO_3$ (data not shown).

CLUSTAL O(1.2.4) multiple sequence alignment

```
OXA-48          ----MRVLALSAV-FLVASIIGM---------PAVAKEWQENKSWNAHFTEHKSQGVVVL     46
OXA-2           -----MAIRIFAILFSIFSLATF---------AHAQEGTLERSDWRKFFSEFQAKGTIVV     46
OXA-10          -------MKTFAA-YVIIACLSS---------TALAGSITENTSWNKEFSAEAVNGVFVL     43
OXA-14          -------MKTFAA-YVIIACLSS---------TALAGSITENTSWNKEFSAEAVNGVFVL     43
OXA-16          -----------------IACLSS---------TALAGSITENTSWNKEFSAEAVNGVFVL     34
OXA-27          MNKYFTCYVVA--SLFLSGCTVQHNL---INETPSQIVQGHNQVIHQYFDEKNTSGVLVI     55
OXA-50          ----MRPLLFSA-LLLLSGH-------------TQASEWNDSQAVDKLFGAAGVKGTFVL     42
OXA-24          MKKFILPIFSISILVSLSACSSIKTK---SEDNFHISSQQHEKAIKSYFDEAQTQGVIII     57
OXA-25          MKKFILPIFSISILVSLSACSSIKTK---SEDNFHISSQQHEKAIKSYFDEAQTQGVIII     57
OXA-26          MKKFILPIFSISILVSLSACSSIKTK---SEDNFHISSQQHEKAIKSYFDEAQTQGVIII     57
OXA-40          MKKFILPIFSISILVSLSACSSIKTK---SEDNFHISSQQHEKAIKSYFDEAQTQGVIII     57
OXA-58          -MKLLKILSLVCLSISIGACAEHSMSRAKTSTIPQVNNSIIDQNVQALFNEISADAVFVT     59
OXA-163         ----MRVLALSAV-FLVASIIGM---------PAVAKEWQENKSWNAHFTEHKSQGVVVL     46
                                  .                             *     ....:

OXA-48          WNENKQQ--GFTNNLKRANQAFLPASTFKIPNSLIALDLGVVKDEHQVFKWDGQTRDIAT     104
OXA-2           ADERQADRAMLVFDPVRSKKRYSPASTFKIPHTLFALDAGAVRDEFQIFRWDGVNRGFAG     106
OXA-10          CKSSSKS--CATNDLARASKEYLPASTFKIPNAIIGLETGVIKNEHQVFKWDGKPRAMKQ     101
OXA-14          CKSSSKS--CATNDLARASKEYLPASTFKIPNAIIGLETGVIKNEHQVFKWDGKPRAMKQ     101
OXA-16          CKSSSKS--CATNDLARASKEYLPASTFKIPNAIIGLETGVIKNEHQVFKWDGKPRAMKQ     92
OXA-27          QTDKKIN--LYGNALSRANTEYVPASTFKMLNALIGLENQKA-DINEIFKWKGEKRSFTA     112
OXA-50          YDVQRQR--YVGHDRERAETRFVPASTYKVANSLIGLSTGAVRSADEVLSYGGKPQRFKA     100
OXA-24          KEGKNLS--TYGNALARANKEYVPASTFKMLNALIGLENHKA-TTNEIFKWDGKKRTYPM     114
OXA-25          KEGKNLS--TYGNALARANKEYVPASTFKMLNALIGLENHKA-TTNEIFKWDGKKRTYPM     114
OXA-26          KEGKNLS--TYGNALARANKEYVPASTFKMLNALIGLENHKA-TTNEIFKWDGKKRTYPM     114
OXA-40          KEGKNLS--TYGNALARANKEYVPASTFKMLNALIGLENHKA-TTNEIFKWDGKKRTYPM     114
OXA-58          YDGQNIK--KYGTHLDRAKTAYIPASTFKIANALIGLENHKA-TSTEIFKWDGKPRFFKA     116
OXA-163         WNENKQQ--GFTNNLKRANQAFLPASTFKIPNSLIALDLGVVKDEHQVFKWDGQTRDIAT     104
                                *:.  : ****:*: ::::.*.        ::: : *  :

OXA-48          WNRDHNLITAMKYSVVPVYQEFARQIGEARMSKMLHAFDYGNEDISGNVDSFWLDGGIRI     164
OXA-2           HNQDQDLRSAMRNSTVWVYELFAKEIGDDKARRYLKKIDYGNADPSTSNGDYWIEGSLAI     166
OXA-10          WERDLTLRGAIQVSAVPVFQQIAREVGEVRMQKYLKKFSYGNQNISGGIDKFWLEGQLRI     161
OXA-14          WERDLTLRGAIQVSAVPVFQQIAREVGEVRMQKYLKKFSYGNQNISGGIDKFWLEDQLRI     161
OXA-16          WERDLTLRGAIQVSAVPVFQQITREVGEVRMQKYLKKFSYGNQNISGGIDKFWLEDQLRI     152
OXA-27          WEKDMTLGEAMKLSAVPVYQELARRIGLDLMQKEVKRIGFGNAEIGQQVDNFWLVGPLKV     172
OXA-50          WEHDMSLRDAIKASNVPVYQELARRIGLERMRANVSRLGYGNAEIGQVVDNFWLVGPLKI     160
OXA-24          WEKDMTLGEAMALSAVPVYQELARRTGLELMQKEVKRVNFGNTNIGTQVDNFWLVGPLKI     174
OXA-25          WEKDMTLGEAMALSAVPVYQELARRTGLELMQKEVKRVNFGNTNIGTQVDNFWLVGPLKI     174
OXA-26          WEKDMTLGEAMALSAVPVYQELARRTGLELMQKEVKRVNFGNTNIGTQVDNFWLVGPLKI     174
OXA-40          WEKDMTLGEAMALSAVPVYQELARRTGLELMQKEVKRVNFGNTNIGTQVDNFWLVGPLKI     174
OXA-58          WDKDFTLGEAMQASTVPVYQELARRIGPSLMQSELQRIGYGNMQIGTEVDQFWLKGPLTI     176
OXA-163         WNRDHNLITAMKYSVVPVYQEFARQIGEARMSKMLHAFDYGNEDISGNVDSFWLDGGIRI     164
                 ::*  *  *:  * * *:: :::. *       :  ..:** : .   ..:*: . : :
```

```
OXA-48     SATEQISFLRKLYHNKLHVSERSQRIVKQAMLTEANGDYIIRAKTGYS---TRIEPKIGW    221
OXA-2      SAQEQIAFLRKLYRNELPFRVEHQRLVKDLMIVEAGRNWILRAKTGWEG-------RMGW    219
OXA-10     SAVNQVEFLESLYLNKLSASKENQLIVKEALVTEAAPEYLVHSKTGFSGVGTESNPGVAW    221
OXA-14     SAVNQVEFLESLYLNKLSASKENQLIVKEALVTEAAPEYLVHSKTGFSGVGTESNPGVAW    221
OXA-16     SAVNQVEFLESLYLNKLSASKENQLIVKEALVTEAAPEYLVHSKTGFSGVGTESNPGVAW    212
OXA-27     TPIQEVEFVSQLAHTQLPFSEKVQANVKNMLLLEESNGYKIFGKTGWA---MDIKPQVGW    229
OXA-50     SAMEQTRFLLRLAQGELPFPAPVQSTVRAMTLLESSPGWELHGKTGWC---FDCTPELGW    217
OXA-24     TPVQEVNFADDLAHNRLPFKLETQEEVKKMLLIKEVNGSKIYAKSGWG---MGVTPQVGW    231
OXA-25     TPVQEVNFADDLAHNRLPFKLETQEEVEKMLLIKEVNGSKIYAKSGWG---MGVTPQVGW    231
OXA-26     TPVQEVNFADDLAHNRLPFKLETQEEVKKMLLIKEVNGSKIYAKSGWG---MGVTPQVGW    231
OXA-40     TPVQEVNFADDLAHNRLPFKLETQEEVKKMLLIKEVNGSKIYAKSGWG---MGVTPQVGW    231
OXA-58     TPIQEVKFVYDLAQGQLPFKPEVQQQVKEMLYVERRGENRLYAKSGWG---MAVDPQVGW    233
OXA-163    SATEQISFLRKLYHNKLHVSERSQRIVKQAMLTEANGDYIIRAKTGYD---T----KIGW    217
           :  ::  *   *   .*      *  *.     :      : .*:*:          :.*

OXA-48     WVGWVELDD-NVWFFAMNMDMPTSDG-LGLRQAITKEVLKQEKIIP-----------  265
OXA-2      WVGWVEWPT-GSVFFALNIDTPNRMDDLFKREAIVRAILRSIEALPPNPAVNSDAAR  275
OXA-10     WVGWVEKET-EVYFFAFNMDIDNESK-LPLRKSIPTKIMESEGIIGG----------  266
OXA-14     WVGWVEKET-EVYFFAFNMDIDNESK-LPLRKSIPTKIMESEGIIGG----------  266
OXA-16     WVGWVEKET-EVYFFAFNMDIDNESK-LPLRKSIPTKIMESEGIIGG----------  257
OXA-27     LTGWVEQPDGKIVAFALKMEMRSEMP-ASIRNELLMKSLKQLNII------------  273
OXA-50     WVGWVKRN-ERLYGFALNIDMPGGEADIGKRVELGKASLKALGILP-----------  262
OXA-24     LTGWVEQANGKKIPFSLNLEMKEGMS-GSIRNEITYKSLENLGII------------  275
OXA-25     LTGWVEQANGKKIPFSLNLEMKEGMS-GSIRNEITYKLLENLGII------------  275
OXA-26     LTGWVEQANGKKIPFSLNLEMKEGMT-GSIRNEITYKSLENLGII------------  275
OXA-40     LTGWVEQANGKKIPFSLNLEMKEGMS-GSIRNEITYKSLENLGII------------  275
OXA-58     YVGFVEKADGQVVAFALNMQMKAGDD-IALRKQLSLDVLDKLGVFHYL---------  280
OXA-163    WVGWVELDD-NVWFFAMNMDMPTSDG-LGLRQAITKEVLKQEKIIP-----------  261
            .*:*:        *:::::          *  :    :     :
```

**Supplementary Figure 30** Sequence alignment of OXA-48 like carbapenemases to show R250 is strictly conserved.

It predicts that the biphasic kinetics involving R250 and chloride ion are probably highly common. By comparison, R214 is not conserved.

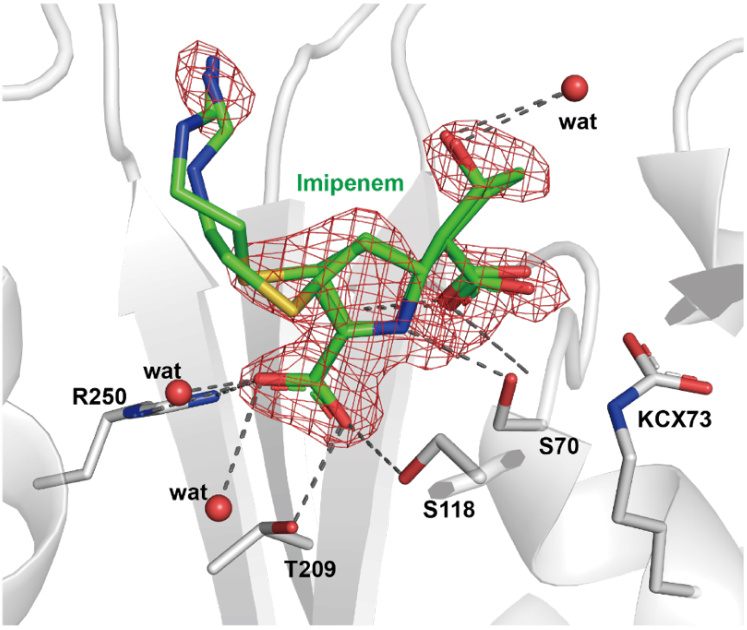

**Supplementary Figure 31** The crystal structure of OXA-$48_{WT}$-imipenem product complex.

The initial $F_o$−$F_c$ electron density map of imipenem is shown as red mesh, contoured at 3σ (0.3081 $e^-$/$A^3$). The crystal structure was obtained by soaking in 500 mM NaBr first, then 64 mM imipenem into the apo OXA-$48_{WT}$ crystal drop.

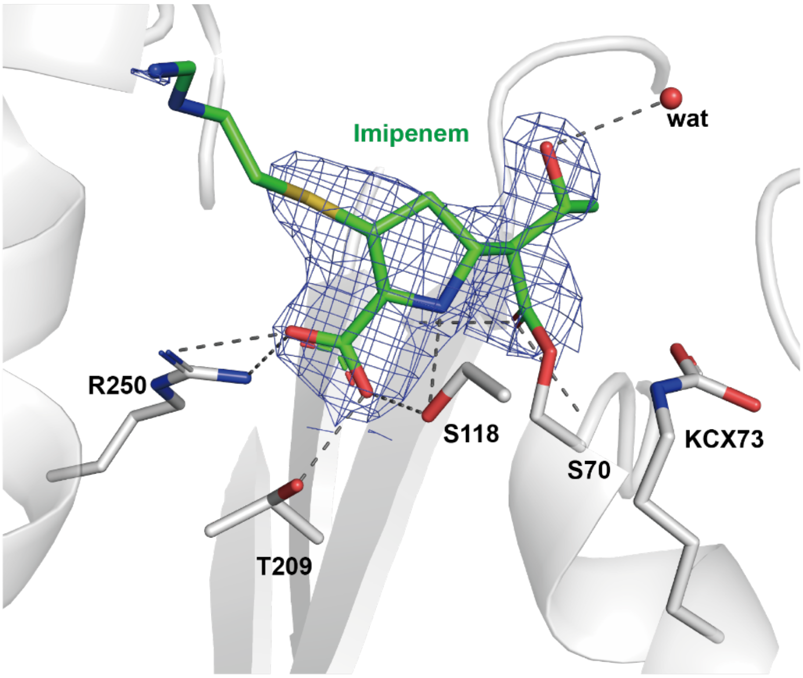

**Supplementary Figure 32** The crystal structure of OXA-$48_{WT}$-imipenem acyl-intermediate complex without Br bound.

The initial, unbiased 2$F_o$−$F_c$ electron density map of imipenem is shown as blue mesh, contoured at 1σ (0.2435 $e^-$/$A^3$). The crystal structure was obtained by soaking in 500 mM NaBr first, then 64 mM imipenem into the apo OXA-$48_{WT}$ crystal drop.

**General information.**

Imipenem was purchased from Chengdu Ai Keda Chemical Technology Co. Ltd. (China) and Merck (UK). All the other antibiotics were purchased from TCI (TCI Shanghai, China) and Merck (UK). All the non-antibiotic chemicals used in this study were analytical grade. The clinical isolates of carbapenemase-producing *Klebsiella pneumoniae* were kindly supplied by Dr Jin-E Lei from the First Affiliated Hospital of Xi'an Jiaotong University (Xi'an, China) and had been previously confirmed by DNA sequencing[2].

**Bacterial growth assay.**

Bacterial growth assays were performed using clinical *Klebsiella pneumoniae* strains expressing carbapenemases OXA-48, KPC or NDM gene, respectively. The spectrophotometric method was used to detect bacterial growth through the measurement of the opacity of the cultures (at $\lambda$ = 600 nm). The bacteria cultured overnight (same opacity in TB medium) were diluted 1000 times into TB medium containing antibiotics (20 μg/mL imipenem), chloride (400 mM) or both, respectively. These bacteria cultures were incubated at 37 °C, 180 rpm for 12 h. The optical density ($\lambda$ = 600 nm) of the bacteria was monitored using a spectrophotometer (Varioskan Flash, Thermo Fisher Scientific) every 1 h during the incubation.

**Calorimetric assays for living bacteria.**

Cell-based calorimetric assays were performed using clinical *Klebsiella pneumoniae* strains expressing OXA-48, KPC or NDM gene. Bacterial suspensions were prepared as described previously[2]. Experiments were performed by injecting 10 μL imipenem stock solution (final concentration 400 μM), preloaded in the syringe, into a sample cell filled with 210 μL of bacterial suspensions ($OD_{600}$ = 4.0 for OXA48-*Kp*, $OD_{600}$ = 1.5 for KPC-*Kp*, $OD_{600}$ = 0.1 for NDM-*Kp*), both prepared in the buffer (50 mM NaPi, pH = 7.5, 1 mM $NaHCO_3$, with 0-400 mM sodium chloride, 1 mM $ZnSO_4$ added for NDM-*Kp*). The control experiment was performed by injecting buffer into bacterial suspensions only. Real-time changes in heat-flow were recorded continuously until returning to baseline.

**Molecular cloning.**

The *bla*OXA-48 gene (GenBank: AAP70012.1) from residue K23 was codon-optimized for *E. coli*, synthesized (Sangon Biotech) and cloned into the pET9a expression vector using the *NdeI*/*BamHI* restriction sites. The resulting plasmid encodes an N-terminally $His_6$-tagged protein.

**Site-directed mutagenesis.**

The mutations in this work were introduced into $OXA\text{-}48_{WT}$ by site-directed mutagenesis. 20 ng of pET9a-*bla*OXA-48 template DNA was mixed with 10 μL 5x Phusion™ HF buffer, 0.2 μM forward and

0.2 µM reverse primer pairs (**Table S6**), 0.2 mM dNTP-mix, 1.5 µL DMSO, and 1 U Phusion™ High–Fidelity DNA Polymerase (Thermo Fisher Scientific; Waltham, USA) in a 50 µL reaction. The PCR reaction was performed with initial 3 min denaturation at 98 °C, followed by 30 cycles of 40 s denaturation at 98 °C, 15 s annealing at 63 °C, and 2.5 min extension at 72 °C. And a final extension was performed at 72 °C for 10 min before cooled to 4 °C. PCR products were immediately digested by *DpnI* (Thermo Fisher Scientific; Waltham, USA) for 2 h at 37 °C and transformed into *E. coli* TOP10 competent cells. The plasmids were extracted and purified, and subsequently verified by sequencing.

**Supplementary Table 6** Oligonucleotides used for sequencing and mutagenesis.

| Primer | Sequence |
|---|---|
| T7 | TAATACGACTCACTATAGGG |
| T7-term | GCTAGTTATTGCTCAGCGG |
| OXA48-R214A-f | GAAAACCGGTTACAGCACC**GCT**ATCGAACCGAAAATCGG |
| OXA48-R214A-r | CCGATTTTCGGTTCGAT**AGC**GGTGCTGTAACCGGTTTTC |
| OXA48-R250A-f | GATGGTCTGGGACTG**GCT**CAAGCGATCACCAAAGAAG |
| OXA48-R250A-r | CTTCTTTGGTGATCGCTTG**AGC**CAGTCCCAGACCATC |
| OXA48-E185A/R186A -f | CTGCACGTTAGC**GCGGCC**TCTCAG |
| OXA48-E185A/R186A-r | CTGAGA**GGCCGC**GCTAACGTGCAG |
| OXA48-R206A-f | GAACGGCGATTACATCATT**GCT**GCGAAAACC |
| OXA48-R206A-r | GGTTTTCGC**AGC**AATGATGTAATCGCCGTTC |
| OXA48-AcK73-f | GGCGTCTACTTTC**TAG**ATCCCGAACAGCCTG |
| OXA48-AcK73-r | CAGGCTGTTCGGGAT**CTA**GAAAGTAGACGCC |

**Gene expression and protein purification of OXA-48$_{WT}$ and its variants OXA-48$_{R250A}$, OXA-48$_{R214A}$, and OXA-48$_{E185A/R186A/R206A}$.**

All OXA-48$_{WT}$ and its variants OXA-48$_{R250A}$, OXA-48$_{R214A}$, and OXA-48$_{E185A/R186A/R206A}$ were expressed in *E. coli* BL21 Star (DE3) strain (New England Biolabs; Ipswich, USA). The transformed cells were selected with 100 µg/mL Kanamycin (TCI) on Terrific Broth (TB) agar medium (11.8 g/L tryptone, 23.6 g/L yeast extract, 9.4 g/L $K_2HPO_4$, 2.2 g/L $KH_2PO_4$ and 4 mL/L glycerin) by overnight incubation at 37 °C. Transformed cells were grown in TB media containing 100 µg/mL Kanamycin at 37 °C until the $OD_{600}$ reached 0.6. The culture was cooled to 18 °C before a final concentration of 0.1 mM isopropyl-β-D-thiogalactopyranoside (IPTG, Sigma-Aldrich) was added, and then further incubated for 24 h at 160 rpm. Cells were subsequently harvested by centrifugation at 8000 rpm for 10 min at 4 °C, resuspended in 20 mL buffer A (150 mM NaPi, 25 mM imidazole, pH 7.5) with lysozyme (Sangon Biotech, Shanghai, 0.5 mg/mL), and incubated at 4 °C for 30 min. The cells were lysed by sonication on ice (pulse interval, 2s/2s (on/off); duration, 30 min) and the lysate was centrifuged at 18000 rpm, 4 °C for 60 min. The supernatant was filtered through a 0.45 µm syringe filter before being loaded onto a 5 mL HisTrap™ column (GE Healthcare) which had been pre-equilibrated with 50 mL buffer A. Subsequently, the column was washed with 50 mL buffer A and the protein was eluted with 20 mL buffer B (150 mM NaPi, 250 mM imidazole, pH 7.5). The fractions containing the eluted protein, confirmed by sodium dodecyl sulfate-polyacrylamide gel electrophoresis (SDS-PAGE), was concentrated and further purified by size exclusion chromatography (SEC) on a HiLoad 16/600 Superdex 200pg column (GE Healthcare) with 100 mM HEPES buffer, pH 7.5. Purity of the collected sample was checked by SDS-PAGE before buffer exchange into required buffers for further experiments.

**Gene expression and protein purification of OXA-48$_{AcK73}$.**

BL21(DE3) was co-transformed with pET9a-OXA48-K73TAG plasmid and the plasmid of the tRNA synthetase by heat shock at 42 °C for 30 seconds. The cells were recovered in 950 μL SOC for 2 h at 37 °C before 100 μL was spread on an agar plate containing kanamycin (50 μg/mL) and spectinomycin (100 μg/mL). LB containing kanamycin (50 μg/mL) and spectinomycin (100 μg/mL) was inoculated with a single colony of transformed cells and incubated at 37 °C for 16 h at 180 rpm. Although AcK73 in the active site is virtually inaccessible for the deacetylases from the *E. coli* strain used, an extra step of caution was taken by adding nicotinamide to fully inhibit potential deacetylase activity in the culture. Thus, the inoculated LB media contained 50 μg/mL kanamycin and 100 μg/mL spectinomycin and were incubated at 37 °C. At $OD_{600}$ = 0.4, nicotinamide (20 mM) and L-acetyllysine (5 mM) were added to the media. After 30 mins, protein expression was induced with 0.5 mM IPTG and the cultures were incubated at 16 °C for 18 hours. The media was centrifuged at 6000 rpm for 20 mins at 4 °C. The cell paste was re-suspended in 30 mL of buffer (100 mM NaPi, 20 mM imidazole, pH 8.0) containing PMSF (0.5 mM). Cells were lysed using a sonication cycle of 10 s/50 s (on/off) for 6 minutes at 4 °C. Soluble material was separated from cell debris by centrifugation at 18000 rpm for 50 mins at 4 °C. The supernatant was applied to a pre-equilibrated Ni-NTA column and washed with five column volumes of the same buffer, and step gradients of increasing imidazole were used for washing and elution steps. Based on SDS-PAGE, elution fractions collected using 200 mM imidazole were concentrated with a centrifugal concentrator (10,000 MWCO). The mixture was diluted with buffer (100 mM NaPi, pH 7.5) to remove residual imidazole.

**General procedure for ITC assays.**

All the ITC calorimetric assays were carried out at 25 °C in the buffer, with a stirring speed of 750 rpm in an ITC-200 calorimeter (Malvern Instruments Ltd., UK). The reaction was initiated by injecting preloaded imipenem solution into the sample cell containing 210 μL enzyme solution; both are prepared in the same buffer. The reference cell of the calorimeter was loaded with the buffer only. Real-time thermal power changes were recorded continuously until the substrate was fully consumed. Data were plotted using Origin 2017 software.

**Effect of chloride concentrations on the hydrolysis of imipenem by OXA-48$_{WT}$ and its variants.**

To monitor the hydrolysis kinetic curves of imipenem by OXA-$48_{WT}$ and its variants in the presence of various chloride ion concentrations (as shown in **Fig. 2, Fig. S8**), experiments were performed by injecting 10 µL imipenem solution (400 µM for OXA-$48_{R214A}$, 200 µM for other enzymes) into the sample cell filled with 210 µL of OXA-$48_{WT}$ (100 nM), OXA-$48_{E185A/R186A/R206A}$ (100 nM), OXA-$48_{AcK}$ (8 µM), OXA-$48_{R250A}$ (2 µM) or OXA-$48_{R214A}$ (3 µM), both are prepared in the same buffer (50 mM NaPi, pH 7.5, 1 mM $NaHCO_3$, with 0, 100, 400 mM NaCl). The concentrations in parentheses are the final concentrations.

**UV-spectrometric assays.**

The UV-spectrometric assays were performed in buffer 50 mM NaPi, pH 7.5, 1 mM $NaHCO_3$, using a Mapada UV-3100 spectrophotometer (Shanghai Mapada Instruments Co. Ltd., China) at 25 °C. The assay was carried out in a total volume of 1 mL with 100 nM OXA-48, 200 µM imipenem and a series dilution of chloride (0-800 mM). The decrease of imipenem absorbance was monitored continuously at 300 nm. Data were plotted using Origin 2017 Software.

**Double injection assay.**

To monitor the influence of additional substrate or enzyme on the hydrolysis kinetic curve during the steady state phase of the reaction (**Fig. 3g, h**), experiments were conducted in "multi-injection" mode. Both imipenem and OXA-$48_{WT}$ stocks are in 50 mM NaPi, pH 7.5 buffer, containing 1 mM $NaHCO_3$ and 100 mM chloride. For the double injection assay of the imipenem substrate, 100 nM OXA-$48_{WT}$ was preadded into the sample cell and 4.38 mM imipenem stock was loaded in the syringe. Two successive injections of imipenem (2 x 10 µL) into the sample cell were made at 740 s intervals. For the double injection assay of the enzyme, 200 µM imipenem was preadded in the sample cell and 1.1 µM OXA-$48_{WT}$ stock was loaded in the syringe. Twice successive injections of enzyme (2 x 10 µL) into the sample cell were made at 510 s intervals.

**Enzyme activity recovery assay.**

A titration of different time intervals between two injections of imipenem was carried out to study the time required for the initial activity recovery before the 2nd injection of imipenem. Experiments were performed by injecting the first dose of imipenem solution into the sample cell filled with enzyme, both in the same buffer of 50 mM NaPi, pH 7.5, 1 mM $NaHCO_3$, with 100 or 400 mM NaCl. The same amount of imipenem is injected into the sample cell to initiate the second reaction after the first reaction at different intervals.

**Size exclusion chromatography of OXA-$48_{WT}$ and OXA-$48_{E185A/R186A/R206A}$.**

Protein dimerization determinations were made by size exclusion chromatography on the GE AKTA Purifier 10 FPLC System (GE Healthcare) using a HiLoad 16/600 Superdex 200 pg column (GE Healthcare). The column was run in NaPi buffer 50 mM, pH 7.5 at a flow rate of 1.0 mL/min. A 1 mL protein sample was injected and data analysis were performed using the UNICORN 5.20 software from GE. The retention time was converted to molecular weight according to the calibration curve.

**Protein thermal shift assays.**

Protein thermal shift assays were performed using the StepOne™ Real-Time PCR System (Applied Biosystems, Foster City, California, USA), with MicroAmp® Optical 8-Cap Strip (0.2 mL) from Applied Biosystems. Protein Thermal Shift™ Dye Kit (Thermo Fisher Scientific; Waltham, USA) was used. All pre-steps were setup on ice. A fresh Thermal Shift™ Dye (8x) was prepared by diluting the 1000x Thermal Shift™ with water. The reactions were prepared by pipetting all components into the wells in the following order: (1) 5.0 µL of Protein Thermal Shift™ buffer, (2) 2.5 µL protein stock (final concentration 12.5 µM), (3) 10 µL NaPi buffer (100 mM, pH 7.5, with 0, 80, 200 mM NaCl), (4) 2.5 µL fresh Protein Thermal Shift™ Dye (8x) to a total volume of 20.0 µL. The samples were assayed as triplicates in each measurement. The plate was centrifuged for 1 min at 1000 rpm to remove air bubbles and was kept on ice. The incubation was started at 25 °C for 2 min, followed by a temperature increase to 99 °C with a ramp-up rate of 1 °C/min. Melting curves were analyzed using the StepOne™ Software from Applied Biosystems. The melting temperature ($T_m$) was calculated from the inflexion point of the melting curve (derivative melting point).

**IC50 and EC50 of all halide ions on OXA-48$_{WT}$.**

The effect of halogen ions was examined with NaI, NaBr, NaCl, or NaF using 200 µM imipenem as substrate in a total volume of 1 mL buffer (50 mM NaPi, pH 7.5, with various concentrations of halide ions). Reactions were initiated by adding OXA-48$_{WT}$ to a final concentration of 100 nM and changes in absorbance of imipenem at 300 nm were recorded continuously by Mapada UV-3100 spectrophotometer (Shanghai Mapada Instruments Co. Ltd., China). The initial rate and the steady-state rate in the presence of a series of concentrations of halide ions were calculated. All experiments were done in triplicates. Half maximal inhibition concentration ($IC_{50}$) values in the steady-state reaction phase were calculated by plotting the percentage of inhibition against halide concentration; Half maximal effective concentration (EC50) values in the initial reaction phase were also determined by plotting the percentage of acceleration against halide concentration using GraphPad Prism 8 software[3].

**Protein crystallization.**

All OXA-48 proteins for crystallography were in 100 mM HEPES buffer, pH 7.5, and strictly no chloride present.

**OXA-48$_{WT}$-I$^-$ complex:**

OXA-48$_{WT}$ apo protein crystals were grown at 18 °C using the hanging drop vapor diffusion method by mixing 1 µL of the protein stock (10 mg/mL) and an equal volume of precipitant containing 0.1 M

HEPES, pH 8.0, 10% 1-butanol,10% PEG8000. Well-diffracted crystals were soaked with 100 mM NaI for 1 min at 20 °C before crystals were cryoprotected by precipitant supplemented with 25% v/v glycerol and flash-frozen in liquid nitrogen.

**OXA-48$_{E185A/R186A/R206A}$ apo crystal:**

OXA-48$_{E185A/R186A/R206A}$ variant crystals were grown at 18 °C using the hanging drop vapor diffusion method by mixing 1 µL of the protein stock (10 mg/mL) and an equal volume of precipitant containing 0.2 M $Li_2SO_4$, 21% PEG3350. Well-diffracted crystals were soaked with 100 mM NaBr for 5 min at 20 °C. Precipitant containing 25% v/v glycerol was used as cryoprotectant before crystals were flash frozen in liquid nitrogen.

**OXA-48$_{WT}$-avibactam-$Cl^-$ complexes:**

1 ratio of 500 mM avibactam (in HEPES buffer, pH 7.0) was pre-mixed with 49 ratios of 10 mg/mL OXA-48$_{WT}$ protein stock prior to crystallization (final concentration: 10 mM avibactam was mixed with 9.8 mg/mL OXA-48$_{WT}$ protein. The co-crystallization crystal was grown at 18 °C using the sitting drop vapor diffusion method by mixing 0.2 µL premixed protein solution and 0.2 µL precipitant against 75 µL of reservoir solutions containing 0.1 M HEPES, pH 8.0, 4% 1-butanol, and 10% w/v PEG8000. To obtain the OXA-48$_{WT}$-avibactam-$Cl^-$ complex, well-diffracted OXA-48$_{WT}$-avibactam crystals were soaked with 100 mM NaCl and an extra 5 mM avibactam for 15 min at 20 °C to ensure full occupancy of the ligand. Precipitant containing 25% v/v glycerol was used as cryoprotectant before crystals were flash frozen in liquid nitrogen.

**OXA-48$_{WT}$ apo, OXA-48$_{WT}$-$Br^-$, and OXA-48$_{WT}$-$HCO_3^-$ complexes:**

OXA-48$_{WT}$ apo crystals were grown at 18 °C using the sitting drop vapor diffusion method by mixing 1 µL of the protein stock (10 mg/mL) and an equal volume of precipitant containing 0.1 M HEPES, pH 7.5, 8% 1-butanol, and 11.6% w/v PEG8000. To obtain the OXA-48$_{WT}$-$Br^-$ complex, well-diffracted crystals were soaked with 500 mM NaBr at 20 °C for 5 min. To obtain the OXA-48$_{WT}$-$HCO_3^-$ complex, well-diffracted crystals were soaked with unbuffered 1 M $NaHCO_3$ at 20 °C for 30 min. Precipitant containing 20% v/v PEG400 was used as cryoprotectant before crystals were flash frozen in liquid nitrogen.

**OXA-48$_{R250A}$ apo and OXA-48$_{R250A}$-$Br^-$ complex:**

OXA-48$_{R250A}$ variant crystals were grown at 18 °C using the sitting drop vapor diffusion method by mixing 1 µL of the protein stock (10 mg/mL) and an equal volume of precipitant containing 0.1 M HEPES, pH 7.5, 8% 1-butanol, and 11.6% w/v PEG8000. Well-diffracted crystals were soaked with 250 mM NaBr at 20 °C. 20% v/v PEG400 was used as cryoprotectant before crystals were flash frozen in liquid nitrogen.

We deployed several strategies to capture an acyl-intermediate structure of OXA-48-imipenem with $Br^{-}$ bound. We tried multiple times to add 500 mM NaBr first followed by 64 mM imipenem to the apo OXA-$48_{WT}$ crystal droplet, but only either hydrolyzed product (**Fig S31**, 7PEP) or acyl-intermediate (**Fig S32**, 7PSF) could be observed, with no detection of an anomalous signal from $Br^{-}$ in the active site. But when we added 500 mM NaBr and 32 mM imipenem to the apo OXA-$48_{WT}$ crystal drop together, there was a clear $Br^{-}$ anomalous signal overlapping with the density of imipenem to show $Br^{-}$ and imipenem competing for binding to R250 (**Fig. 3a**, 1.53 Å resolution structure, undeposited). The details are as follows:

**OXA-$48_{WT}$-imipenem intermediate and OXA-$48_{WT}$-imipenem product complexes with $Br^{-}$ bound on the surface (but not in the active site):**

The OXA-$48_{WT}$-imipenem acyl-intermediate and product complexes were both obtained by soaking well-diffracted OXA-$48_{WT}$ apo crystals with 500 mM NaBr first, followed by 67.5 mM imipenem for 5-10 min. Please note, although no $Br^{-}$ anomalous density was observed in the active site, $Br^{-}$ density was consistently observed on the protein dimer interface near E185, R186, and R206, as well as on the protein surface near residues K137.

**OXA-48 $_{WT}$-imipenem-$Br^{-}$ product complexes with $Br^{-}$ bound in the active site:**

The OXA-$48_{WT}$-imipenem-$Br^{-}$ product complex was obtained by soaking well-diffracted crystals with 32 mM imipenem and 500 mM NaBr together as a premixed stock solution at 20 °C for 5-10 min. Precipitant containing 20% v/v PEG400 was used as cryoprotectant for crystals before crystals were flash frozen in liquid nitrogen. The anomalous density for $Br^{-}$ was consistently observed in the active site near R250. The $Br^{-}$ density was also observed on the protein dimer interface near E185, R186, and R206, as well as on the protein surface near residues R134, K137, and R174.

**OXA-$48_{AcK}$-imipenem, OXA-$48_{AcK}$-imipenem-$Br^{-}$ and OXA-$48_{AcK}$-oxacillin intermediate complexes:**

OXA-$48_{AcK}$ variant apo crystals were grown at 18 °C using the sitting drop vapor diffusion method by mixing 1 µL of the protein stock (10 mg/mL) and an equal volume of precipitant containing 0.1 M HEPES, pH 7.5, 8% 1-butanol, and 11.6% w/v PEG8000. To obtain the OXA-$48_{AcK}$-imipenem acyl-intermediate complex, well-diffracted crystals were soaked with 30 mM imipenem at 20 °C. The OXA-$48_{AcK}$-imipenem-$Br^{-}$ intermediate complex was obtained by soaking OXA-$48_{AcK}$ variant apo crystals directly with imipenem powder and 250 mM NaBr at 20 °C for 5 to 10 min. To obtain the OXA-$48_{AcK}$-oxacillin acyl-intermediate complex, well-diffracted crystals were soaked with 30 mM oxacillin at 20 °C. Precipitant containing 20% v/v PEG400 was used as cryoprotectant before crystals were flash frozen in liquid nitrogen.

**Crystallography data collection, processing and refinement.**

Crystal diffraction data for structures of OXA-$48_{WT}$-$I^-$ complex, OXA-$48_{E185A/R186A/R206A}$ apo, and OXA-$48_{WT}$-avibactam-$Cl^-$ complex were collected at 100 K on beamline 17U1 of the Shanghai Synchrotron Radiation Facility (SSRF, Shanghai). Data were indexed and integrated using XDS[4] and HKL-3000[5], scaled with CCP4i2 suite[6] of programs. Crystal diffraction data for structures of apo OXA-$48_{WT}$, OXA-$48_{WT}$-$Br^-$ complex, OXA-$48_{WT}$ -$HCO_3^-$ complex, apo OXA-$48_{R250A}$, OXA-$48_{AcK}$-imipenem, OXA-$48_{AcK}$-imipenem-$Br^-$ and OXA-$48_{AcK}$-oxacilin intermediate complexes, OXA-$48_{WT}$-imipenem intermediate and OXA-$48_{WT}$-imipenem product complexes, and OXA-48 $_{WT}$-imipenem-$Br^-$ product complex was collected at 100 K on beamlines i03 and i04 of the Diamond Light Source UK. Reflections were automatically processed with the *xia2* pipeline[7] of the CCP4i2 software suite[6]. All structures were solved by molecular replacement using MOLREP[8] or Phaser[9] using OXA-48 PDB ID 4S2P as the search model. The structures were built and refined using iterative cycles with Crystallography Object-Oriented Toolkit (Coot[10]) and REFMAC5[11]. The NCS restraint parameters were used at later stages. Data collection and refinement statistics are provided in **Table S3**.

**Michaelis-Menten kinetic for OXA-$48_{WT}$ and OXA-$48_{AcK73}$.**

Michaelis-Menten kinetics for OXA-$48_{WT}$ and OXA-$48_{AcK73}$ were measured for the enzyme-catalyzed hydrolysis of imipenem (Merck). The rates of hydrolysis of imipenem by 0.1 μM OXA-$48_{WT}$ were monitored by following the UV absorption at 300 nm for the initial 30 seconds. All reactions were performed at 25 °C in the buffer of 100 mM NaPi, 50 mM $NaHCO_3$, pH 7.5, with strictly no chloride. 0-175 mM imipenem and the buffer solution were premixed in a 1 cm cuvette, and then the enzyme stock was added to a total volume of 1 mL. Kinetic parameters ($k_{cat}$, $K_M$, $k_{cat}/K_M$) were calculated using the Michaelis-Menten equation $y = E_t * k_{cat} * x/(K_M + x)$, from the GraphPad Prism 6.01 Software[12]. All experiments were performed in triplicate.

**Molecular docking.**

Covalent docking was performed using Molecular Operating Environment (MOE)[1]. Oxacillin (CID6196) coordinates were retrieved from the PubChem compound database[13]. The protein template is obtained from the H chain of the OXA-$48_{AcK}$-imipenem-$Br^-$ structure. The imipenem ligand and water molecular were removed but the bromide ion was kept. Protein and ligand structures were prepared by using default parameters. The binding site was defined as S70 of the protein and the lactam ring-opening reaction was chosen as the reaction type. The figures of OXA-$48_{AcK}$-oxacillin complex in the inactive conformation were prepared using PyMOL[14].

**Synthesis of $N_α$-(*tert*-butoxycarbonyl)-L-acetyllysine** (**1**).

Commercially available $N_α$-(tert-butoxycarbonyl)-L-lysine (10 g, 40.6 mmol) was dissolved in NaOH (1 M, 100 mL) / THF (100 mL) and the mixture was cooled to 0 °C. Acetyl chloride (3.16 mL, 44.7 mmol) was added dropwise, and the reaction was stirred at room temperature. After 24 h, most of the THF was removed under reduced pressure. The resulting mixture was acidified to pH 4.0 with 1 M HCl and extracted with EtOAc (3 x 100 mL). The organic layer was dried over anhydrous $MgSO_4$, filtered and concentrated under reduced pressure to give the product $N_{\alpha}$-(*tert*-butoxycarbonyl)-L-acetyllysine (**1**) as a white solid (5.5 g, 47%). **$^1$H NMR** (400 MHz, DMSO-$d_6$) δ 7.78 (t, *J* = 4.88 Hz, 1H), 6.99 (d, *J* = 6.4 Hz, 1H), 3.8 (m, 1H), 2.98 (m, 2H), 1.77 (s, 3H), 1.68–1.48 (m, 2H), 1.37 (s, 9H), 1.36–1.21 (m, 4H) ppm. **$^{13}$C NMR** (100 MHz, DMSO-$d_6$) δ = 174.2, 172, 155.6, 77.9, 53.4, 38.3, 30.5, 28.8, 28.2, 23.1, 22.6 ppm. **HRMS**: $C_{13}H_{24}N_2O_5$ : [M - H]$^-$ = 287.1596.

**Synthesis of L-acetyllysine (2).**

$N_{\alpha}$-(*tert*-butoxycarbonyl)-L-acetyllysine (4.8 g, 16.6 mmol) (**1**) was dissolved in DCM (50 mL). TFA (5.76 mL, 83 mmol) was added dropwise, and the reaction was stirred at room temperature. After 18 h, most of the TFA was removed under nitrogen and then completely removed under reduced pressure. The red oil was re-dissolved in deionized water (25 mL) and extracted with DCM (3 x 30 mL). The aqueous phase was concentrated under reduced pressure to give L-acetyllysine (**2**) as a yellow oil (7 g, 99 %). **$^1$H NMR** (400 MHz, $D_2O$) δ = 4.05 (t, *J* = 6Hz, 1H) 3.18 (t, *J* = 6.4 Hz, 2H), 2–1.85 (m, 2H), 1.97 (s, 3H), 1.56 (m, 2H), 1.44 (m, 2H) ppm. **$^{13}$C NMR** (100 MHz, $D_2O$) δ = 174, 172.2, 52.8, 38.8, 29.3, 27.7, 21.7, 21.5 ppm. **HRMS**: $C_8H_{16}N_2O_3$ : [M + H]$^+$ = 189.1242.

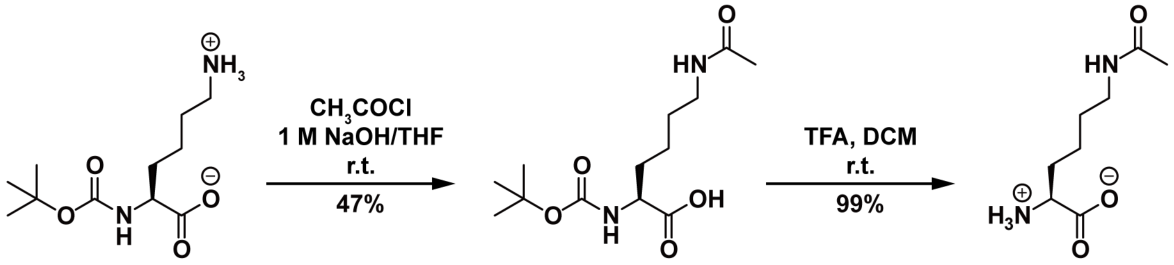

1 2

**Scheme 1.** Synthesis of L-acetyllysine (**2**).

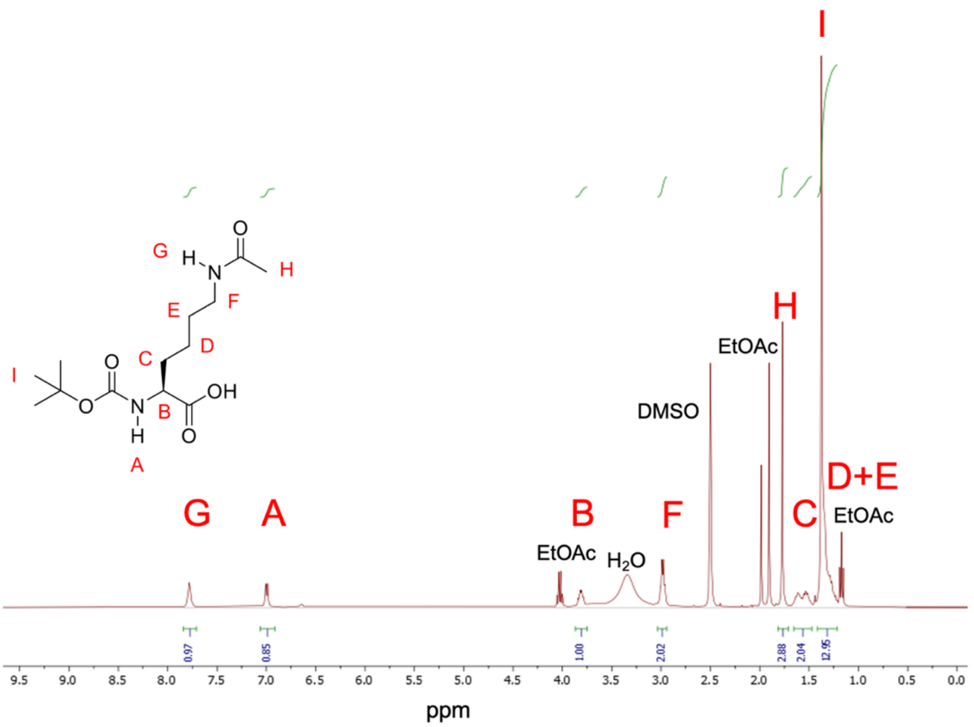

**Supplementary Figure 33** $^{1}$H NMR spectrum of $N_{\alpha}$-(*tert*-butoxycarbonyl)-L-acetyllysine (**1**) in DMSO-$d_6$ (400 MHz).

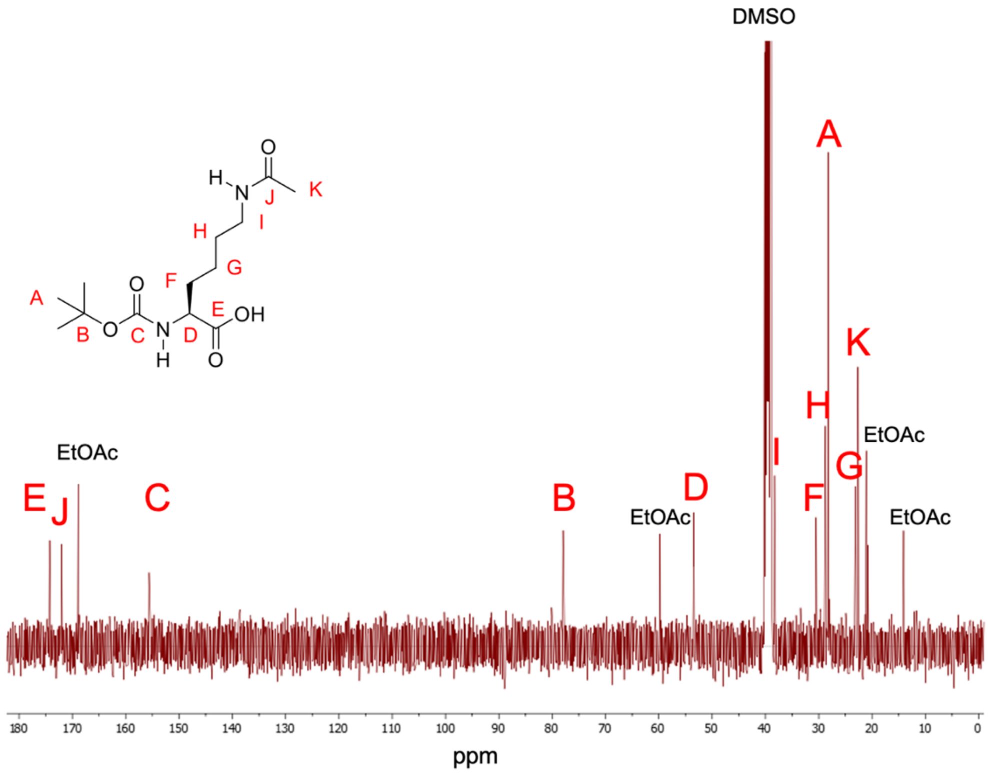

**Supplementary Figure 34** $^{13}$C NMR spectrum of $N_{\alpha}$-(*tert*-butoxycarbonyl)-L-acetyllysine (**1**) in DMSO-$d_6$ (100 MHz).

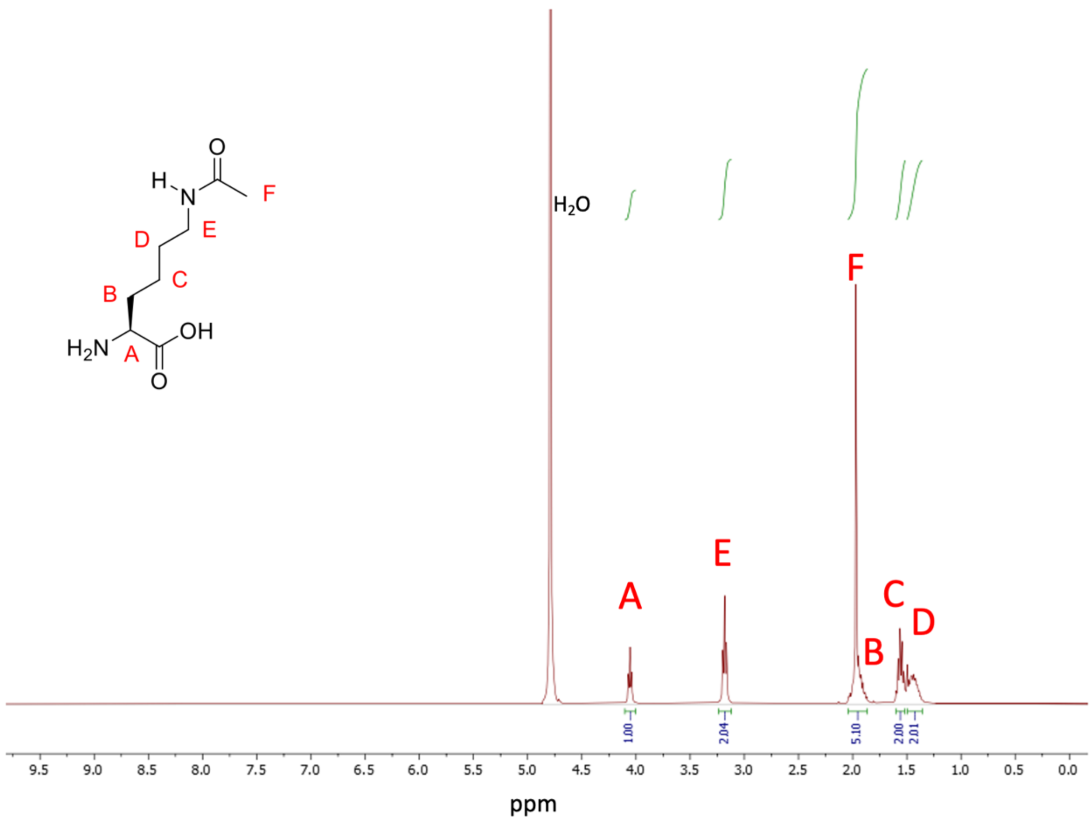

**Supplementary Figure 35** $^{1}$H NMR spectrum of L-acetyllysine (**2**) in $D_2O$ (400 MHz).

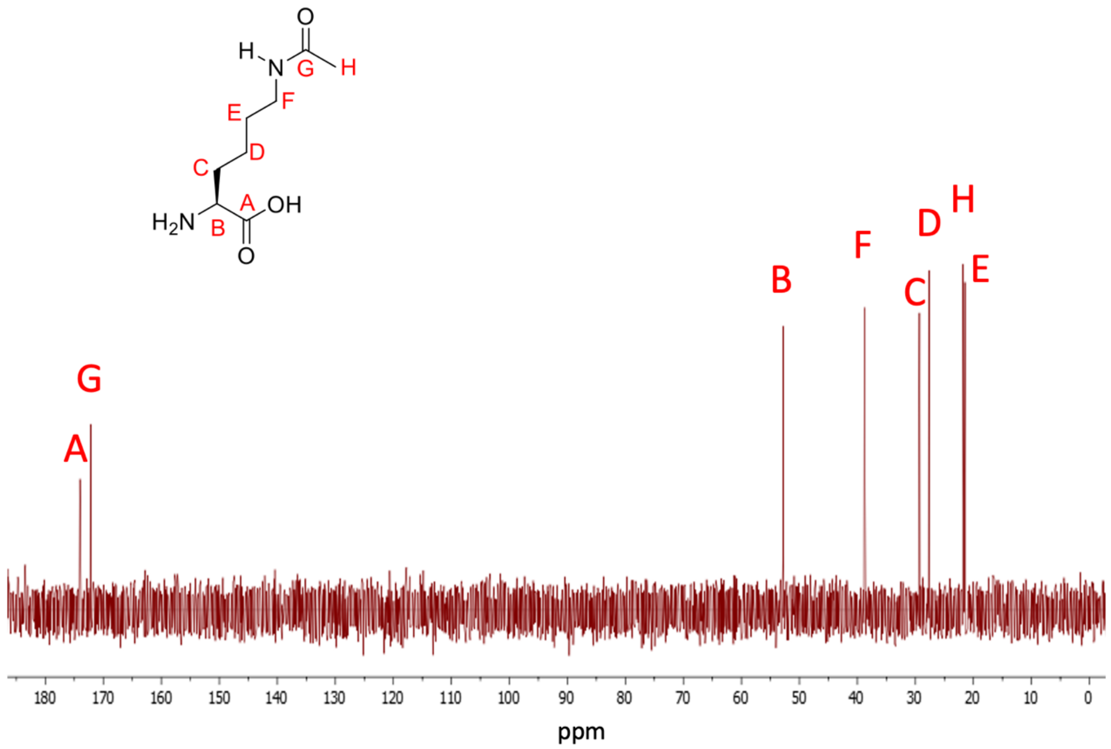

**Supplementary Figure 36** $^{13}$C NMR spectrum of L-acetyllysine (**2**) in $D_2O$ (100 MHz).

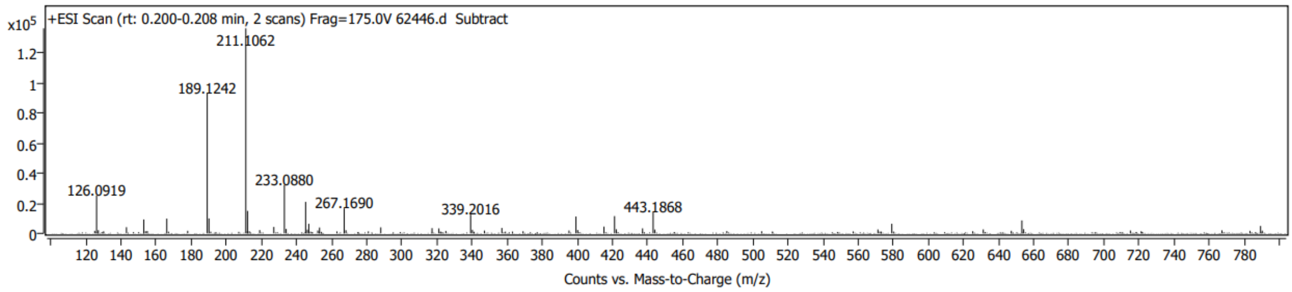

**Supplementary Figure 37** High-resolution MS of L-acetyllysine (**2**).

**Mathematical modelling.**

The standard enzymatic reactions for OXA-48 are as follows:

$E + S \xrightarrow{k_1} EA \xrightarrow{k_2} E + P + H,$ where $E$ is enzyme, $S$ is substrate, $EA$ is the intermediate enzyme complex, $P$ is product, $H$ is heat released by the reaction, and $k_1$ and $k_2$ are the kinetic rates for both reactions. Although these reactions are reversible, in practice the kinetic constants are so skewed that we will consider them irreversible for simplicity.

Deriving the corresponding ordinary differential equations from the law of mass action (which is warranted since the number of molecules in our experiments is very high), we obtain the following system in the absence of chloride:

$$\frac{d[E]}{dt} = -k_1[E][S] + k_2[EA],$$
$$\frac{d[S]}{dt} = -k_1[E][S],$$
$$\frac{d[EA]}{dt} = k_1[E][S] - k_2[EA],$$
$$\frac{dH}{dt} = -ck_2[EA].$$

Here $[X]$ is the concentration of molecule $X$, where $X = E, S$ or $EA$, and $c$ is the conversion constant between moles of product and calories of heat. The variable that is measured experimentally in the ITC experiments is a function of $dH/dt$, the instantaneous heat release (see next section).

We have shown that chloride binds OXA-48's dimer interface, which implies the reversible reaction $E + 2Cl^- \rightleftharpoons E^{Cl}$ where $E^{Cl}$ is the enzyme with more than two chloride ions in the interface and on the surface. Since the concentration of chloride is much larger than that of enzyme in our experiments, it is reasonable to suppose that most of the enzyme is in $E^{Cl}$ form, and we will consider only this form in the following.

This form of the enzyme undergoes the usual reactions, $E^{Cl} + S \xrightarrow{k_1'} E^{Cl}A \xrightarrow{k_2'} E^{Cl} + P + H$, where $k_1'$ and $k_2'$ could in principle be different from $k_1$ and $k_2$ above, which will depend on chloride.

In addition, $E^{Cl}A$ undergoes two extra reactions $E^{Cl}A + Cl^- \underset{k_{-3}'}{\overset{k_3'}{\rightleftharpoons}} E^{Cl}A.Cl$, forming an inactive intermediate complex $E^{Cl}A.Cl$ as the forward reaction, and Cl$^-$ dissociating from the inactive intermediate complex as the reverse reaction.

In the presence of chloride, the new mathematical model is then

$$\frac{d[E^{Cl}]}{dt} = -k_1'[E^{Cl}][S] + k_2'[E^{Cl}A],$$
$$\frac{d[S]}{dt} = -k_1'[E^{Cl}][S],$$
$$\frac{d[E^{Cl}A]}{dt} = k_1'[E^{Cl}][S] - k_2'[E^{Cl}A] - k_3'[E^{Cl}A][Cl^-] + k_{-3}'[E^{Cl}A.Cl],$$
$$\frac{d[E^{Cl}A.Cl]}{dt} = k_3'[E^{Cl}A][Cl^-] - k_{-3}'[E^{Cl}A.Cl],$$
$$\frac{dH}{dt} = -ck_2'[E^{Cl}A].$$

For simplicity of notation, we will drop the bracket notation and merge the two models, using $E$ for both forms of the enzyme, $C$ for the active intermediate $E^{Cl}A$ and $C_2$ for the inactive intermediate $E^{Cl}A.Cl$. The model then becomes

$$\frac{dE}{dt} = -k_1'ES + k_2'C,$$
$$\frac{dS}{dt} = -k_1'ES,$$
$$\frac{dC}{dt} = k_1'ES - k_2'C - k_3'\mathrm{Cl}^-C + k_{-3}'C_2,$$
$$\frac{dC_2}{dt} = k_3'\mathrm{Cl}^-C - k_{-3}'C_2,$$
$$\frac{dH}{dt} = -ck_2'C,$$

Where $k_1$', $k_2$', $k_3$', and $k_{-3}$' are apparent reaction rate constants is that would be taken as functions of chloride. In the following, we will use mathematical arguments to discuss the effect of chloride ion concentration on them.

Working with the "real" heat curve

The shape of titration curves is affected by the calorimeter response time[15]. The signal measured by the calorimeter decreases exponentially with a characteristic time $t_{ITC}$, and so the measured heat release $P_m(t)$ is a convolution between the real signal $P_s(t) = dH/dt$ and an exponential kernel $R(t) = (t_{ITC})^{-1} \exp(-t/t_{ITC})$. Thus,

$$P_m(t) = \frac{1}{t_{ITC}} \int_0^t \exp(-(t-\tau)/t_{ITC}) P_s(\tau)\, d\tau, \qquad (1)$$

which can be solved either by differentiating once or using the Laplace transform, yielding

$$t_{ITC} \frac{dP_m}{dt} + P_m = P_s(t). \qquad (2)$$

Given a measured signal $P_m(s)$, we can numerically obtain from this equation the "real" signal that corresponds to our dynamic equations. In our experiments, $t_{ITC} = 8\,s$, given by the manufacturer of ITC-200 calorimeter (Malvern Instruments Ltd., UK). The "real" and "measured" signals are quite similar, but there are relevant differences especially when there are spikes of heat release, such as at the beginning of the titration experiment (**Fig. S16**).

In the following, we will use the "real" signal $P_s(t)$ to obtain kinetic rates and then use these to simulate our model. In order to compare with the experimental datasets, we will then transform our mathematical variable $dH/dt$ to a measured $P_m$, using the differential equation above. As seen in **Fig. 2**, the ITC data shows a baseline drift, but we will not correct it, as it will not significantly alter our main conclusions.

Dealing with the finite injection time of the substrate

A common problem when modelling ITC curves comes from the fact that the injection of the substrate into the sample cell is not instantaneous, but rather requires a finite amount of time. Typically, this injection is also combined with fast mixing, so that a homogeneous concentration of substrate is quickly attained in the sample cell. Writing this out mathematically is a challenge: ideally, we could derive a partial differential equation that would incorporate the diffusion into the sample cell but deriving a mathematical expression for the mixing term seems rather hard.

Since our main purpose is to extract the reaction rates of imipenem by OXA-48, we have found a simple, alternative way to model the diffusion of the substrate throughout the ITC sample cell by phenomenologically modelling a linear injection into the system. The equation for $E$ then becomes

$$\frac{dE}{dt} = \frac{E_T}{k_0}(1 - \Theta(t - k_0)) - k_1 ES + k_2 C,$$

where $\Theta(t)$ is the Heaviside function that has value 0 if $t < 0$, and 1 if $t > 0$. $k_0$ is the time needed for the substrate to diffuse, and we will set it to values between 20 and 30 s depending on the experiment, which fits the purpose of capturing this initial injection and mixing.

Finding out $c$

Integrating the area under any of our experimental ITC curves, we obtained the total heat released when all the substrate has been hydrolysed. We would get the same value by multiplying the initial substrate concentration ($S_0 = 200\mu M$) times $c$, the conversion rate. Thus, we can estimate $c$ by dividing the total heat released by $S_0$. Using this method, in our ITC experiments with different chloride concentrations, we got values of $c$ between 4.5 and 5.5 calories per mole, increasing linearly with chloride concentration (**Fig. S17**).

As there is no binding heat from chloride, the variability could then come from an intrinsic drift of the baseline inherent from ITC as a technique. Therefore, we will use the estimated value of $c$ for each chloride concentration in the following sections.

No chloride: finding out $k_2$

In the absence of chloride, OXA-48 follows typical Michaelis-Menten kinetics (single phase), meaning the heat release is constant (and maximum). If heat release is constant, then as $dH/dt = -ck_2C$ and both $c$ and $k_2$ are constants, $C$ must also be a constant. Therefore, $dC/dt = 0$, which implies that $k_1ES = k_2C$. But $E + C = E_T$ is the total enzyme concentration, so solving for $E$ and substituting in the above equation we get

$$k_1 E_T S - k_1 C S = k_2 C \implies C = \frac{k_1 E_T S}{k_1 S + k_2} = \frac{E_T}{1 + \frac{k_2}{k_1 S}}.$$

$S \gg E_t$ at this point in the experiment, as the initial substrate concentration is $200 \mu M$, while $E_T = 100\, nM$, three orders of magnitude less. If we assume $k_1$ and $k_2$ are in the same order of magnitude (an assumption that will be checked later), we deduce from this that, in the constant release stage, almost all the OXA-48 enzyme molecules are in $C$ form.

At this point of maximum heat release, we have

$$k_2 = -\frac{dH/dt}{cC} \approx -\frac{dH/dt}{cE_T}$$

and as we know both $c = 4.5 cal M^{-1}$ and $E_T = 100 nM$ we can derive $k_2$ from measuring the value of $dH/dt$ at the constant heat release stage. For the reaction condition with no chloride, we get $k_2 = 2.83 s^{-1}$.

<u>Estimating the order of magnitude of $k_1$</u>

It is not easy to obtain estimates for $k_1$ from simple mathematical arguments like the previous one. We can, however, perform a numerical fit of the model to the no chloride curve. This yields a value of $k_1 = 1.06\, \mu M^{-1} s^{-1}$ and $k_2 = 2.83 s^{-1}$, confirming our previous argument with a good fit to the data (**Fig. S18**).

When there is no chloride, our model only has three parameters, namely $k_0$, $k_1$ and $k_2$. Fixing $k_0 = 30 s$, our numerical routine (using Python's minimize function, see the Github code) yields $k_1 = 1.06\, \mu M^{-1} s^{-1}$ and $k_2 = 2.83 s^{-1}$ with a good fit for the data. The actual value of $k_1$ can change between 1 and 1.6 $\mu M^{-1} s^{-1}$ if we allow the minimization routine to vary $k_0$.

<u>Relationship between $k_2$' and chloride</u>

When repeating the experiment with nonzero concentrations of chloride in solution, we find that the maximum heat release increases as chloride increases. As both $c$ and $E_T$ are the same throughout all experimental conditions, it must be the case that $k_2$' is increasing with chloride.

The maximum heat release in all cases is reached at $t \approx 34\, s$ and by that time the 0 mM chloride treatment has almost reached the maximum heat release. More importantly, by simulating our equations with $k_1 = 1.06\, \mu M^{-1} s^{-1}$ and $k_2 = 2.83 s^{-1}$ (as above) by $t = 34\, s$ the treatment with no chloride has already reached the maximum heat release. If we assume that $k_1$ is not greatly affected by chloride (we will check this assumption later), we deduce that by $t = 34\, s$ almost all the enzyme molecules are in $C$ form in all reactions. That is

$$(dH/dt)_{max} = ck_2' E_T \implies k_2' = \frac{(dH/dt)_{max}}{cE_T}$$ as before.

From this equation we can infer the values of $k_2$' from the data, obtaining a logarithmic relationship $k_2' = 0.55 + 0.93 \log(\mathrm{Cl} + 34.6)$ between $k_2$' and chloride, $R^2$= 0.94 (**Fig. 4d**).

<u>Finding out $k_3$'</u>

After the initial spike in heat release, most of the enzyme is in $C$ state, and suddenly heat starts going down. This is due to the reaction $C \to C_2$ kicking in, which has an apparent rate constant $k_3$'. Once most of the enzyme is in $C$ state, this means that $dE/dt = 0$ and so $k_1' ES = k_2' C$, which means that in the equation $dC/dt$ the first two terms are negligible. But also, because there is still no $C_2$ whatsoever, the last term $k_{-3}' C_2$ is also close to zero. So, at this spike, we have $dC/dt \approx -k_3' Cl^- C$. In other words, $C$ will decrease exponentially at a rate $k_3' Cl^-$, very shortly after the spike. Because $dH/dt$ is proportional to $C$, the heat will also decrease exponentially. We can check this by plotting the data in semilog axes: if $dH/dt = A \exp(-k_3' Cl^- t)$ then $\log dH/dt = \log A - k_3' Cl^- t$, and so we should see a linear relationship between $t$ and $\log dH/dt$, which is the case (**Fig. S19**). Fitting this equation to the data, an inverse relationship between $k_3$' and chloride is obtained (**Fig. 4e**).

Notice that this exponential decrease does not start immediately, but only after a delay of around 37 seconds. This means that the reaction $C + Cl^- \to C_2$ does not start immediately (perhaps because of the mixing). As a result, in order to capture the data, we will set $k_3$' to zero in our equations if $t < 37 s$.

<u>Finding out $k_{-3}$'</u>

Now, although the curves start going down at the same speed, they soon diverge. This is because, as $C_2$ starts to increase, the previous approximation $dC/dt \approx -k_3' Cl^- C$ is no longer valid and the change in heat is no longer exponential, which can be appreciated because the curves are no longer linear in the semilog plot.

The curves eventually reach a new stationary value $(dH/dt)_s$, where $k_3' Cl^- (C)_s = k_{-3}'(C_2)_s$. At this point almost all the enzyme is in either $C$ or $C_2$ form, and so $(C)_s + (C_2)_s = E_T$. From these two equations, we get $k_{-3}' = Cl^- k_3' \frac{C_s}{E_T - C_s}$, so we only need to obtain $C_s$. That is done by measuring the stationary value $(dH/dt)_s = ck_2' C_s$, which leads us to

$$k_{-3}' = Cl^- k_3' \frac{(dH/dt)_s}{ck_2' E_T - (dH/dt)_s}.$$

But $ck_2' E_t = (dH/dt)_{\max}$ , so finally we have

$$k_{-3}' = Cl^- k_3' \frac{(dH/dt)_s}{(dH/dt)_{\max} - (dH/dt)_s}.$$

That is, finding out $k_{-3}$' depends only on the value of $dH/dt$ at its peak ($(dH/dt)_{\max}$) and the value of $dH/dt$ at the plateau ($(dH/dt)_s$). Using this formula, we obtain an inverse relationship between $k_{-3}$' and chloride (**Fig. 4f**).

<u>The relationship between $k_1$' and chloride</u>

As pointed out before, there is no easy way to obtain information about $k_1$' from the equations, and therefore we have turned to numerical optimization. We have fitted the parameters of each of the ITC curves using standard numerical routines in Python, obtaining very similar results to the ones described above. **Fig. S20** shows each data curve and two simulations of the model, one obtained using our analytic approximations (dotted lines) and the other using the numerical fit (dashed lines). The agreement between them is very good. From the numerical fits, we can obtain values for $k_1$' and confirm that it does not depend on chloride (**Fig. S20**).

Using Python's minimize function, we have fitted each ITC curve for every chloride concentration to our model, obtaining the curves above (dashed lines). Comparing them against our analytical approximations (dotted lines), the correspondence between the two is very high. The values of the parameters for each curve can be found in the GitHub code at github.com/pablocatalan/oxa48.

Plotting the values of $k_1$' obtained in the numerical fit to the ITC data versus chloride, we observe there is no relationship between the two variables. In fact, changing the values of this parameter between 1.04 and 1.10 $\mu M^{-1} s^{-1}$ does not lead to significant changes in our simulations (**Fig. 4c**).

Fitting the ITC curves

For each ITC curve in each of the experiments, we applied the previously described methods to obtain the apparent rates (**Fig. S21**). For the exact values of the parameters used in these simulations, see github.com/pablocatalan/oxa48.

Simulating a non-zero hydrolysis rate from the inactive intermediate

We asked whether the inactive intermediate is also able to hydrolyse imipenem. This new model would include a new reaction $C_2 \xrightarrow{k_5} E + P + H$ with a rate $k_5$. The model then becomes

$$
\begin{aligned}
\frac{dE}{dt} &= \frac{E_T}{k_0}(1 - \Theta(t - k_0)) - k_1' ES + k_2' C + k_5 C_2, \\
\frac{dS}{dt} &= -k_1' ES, \\
\frac{dC}{dt} &= k_1' ES - k_2' C - k_3' \mathrm{Cl}^- C + k_{-3}' C_2, \\
\frac{dC_2}{dt} &= k_3' \mathrm{Cl}^- C - k_{-3}' C_2 - k_5 C_2, \\
\frac{dH}{dt} &= -c(k_2' C + k_5 C_2),
\end{aligned}
$$

We then simulated this model for different values of $k_5$ and the values of the other parameters. We see that this new model fails to capture the ITC data even for very low values of $k_5$ from 0 to 0.5 $s^{-1}$, which suggests that there is no hydrolysis from the inactive intermediate (**Fig. S22**).

Simulating the dynamics for the interface mutant

With the data in **Fig S11c, d**, we use our mathematical analysis to study the apparent reaction rates and their relationship to chloride concentration in the interface mutant OXA-48$_{E185A/R186A/R206A}$ as well (**Fig. S23**). Simulation plots using the same equations (dashed lines) agree quite well with the data (solid lines). Parameter values are shown in **Table 1**.
